# Exploring correlations in genetic and cultural variation across language families in Northeast Asia

**DOI:** 10.1101/513929

**Authors:** Hiromi Matsumae, Peter Ranacher, Patrick E. Savage, Damián E. Blasi, Thomas E. Currie, Kae Kognebuchi, Nao Nishida, Takehiro Sato, Hideyuki Tanabe, Atsushi Tajima, Steven Brown, Mark Stoneking, Kentaro K. Shimizu, Hiroki Oota, Balthasar Bickel

## Abstract

Culture evolves in ways that are analogous to, but distinct from, genomes. Previous studies examined similarities between cultural variation and genetic variation (population history) at small scales within language families, but few studies empirically investigated such parallels across language families using diverse cultural data. We report an analysis comparing culture and genomes from in and around Northeast Asia spanning 11 language families. We extract and summarize the variation in language (grammar, phonology, lexicon), music (song structure, performance style), and genomes (genome-wide SNPs) and test for correlations. We find that grammatical structure correlates with population history. Recent contact and shared descent fail to explain the signal, suggesting relationships that arose before the formation of current families. Our results suggest that grammar might be a cultural indicator of population history, while also demonstrating differences among cultural and genetic relationships that highlight the complex nature of human history.

## Introduction

The history of our species has involved many examples of large-scale migrations and other movements of people. These processes have helped shape both our genetic and cultural diversity(*1*). While humans are relatively homogenous genetically, compared to other species, there are subtle population-level differences in genetic variation that can be observed at different geographical scales(*2*). Furthermore, while there are universal features of human behavior (e.g., all known societies have language and music(*3*–*5*)), our cultural diversity is immense. For example, we speak or sign over 7,000 mutually unintelligible languages(*6*), and for each ethno-linguistic group there tend to be many different musical styles(*7*). Researchers have long been interested in reconstructing the history of global migrations and diversification by combining historical and archaeological data with patterns of present-day biological and cultural diversity. Going back as far as Darwin, many researchers have argued that cultural evolutionary histories will tend to mirror biological evolutionary histories(*8*–*18*). However, differences in the ways that cultural traits and genomes are transmitted mean that genetic and cultural variation may in fact be explained by different historical processes(*19*–*22*). Major advances in both population genetics and cultural evolution since the second half of the 20^th^ century now allow us to test these ideas more readily by matching genetic and cultural data(*19*, *23*–*25*).

The cultural evolution of language has proven particularly fruitful for understanding past population history(*26*–*28*). A classic approach involves identifying and analyzing sets of homologous (cognate) words among languages. This lexical approach allows for the reconstruction of evolutionary lineages and relationships within a single language family, such as Indo-European or Austronesian (*26*, *27*, *29*). However, lexical methods cannot usually be applied to multiple language families(*28*) as they do not share robustly identifiable cognates due to a time limit of approximately 10,000 years, after which phylogenetic signals are generally lost(*30*–*32*). An alternative approach is to study the distribution of features of grammar and phonology, such as the relative order of word classes in sentences or the presence of nasal consonants. While structural data in language tends to evolve too fast to preserve phylogenetic signals of language families(*33*, *34*) and the history of lexica and structure might be partially independent as for example in the emergence of creole languages(*22*), the geographical distribution of language structure often points to contact-induced parallels in the evolution of entire sets of language families beyond their individual time depths(*35*, *36*).

Yet language is only one out of many complex cultural traits that could serve as a proxy for deep history. It has been proposed that music may preserve even deeper cultural history than language(*37*–*43*). Standardized musical classification schemes (based on features such as rhythm, pitch, and singing style) can be used to quantify patterns of musical diversity among populations for the sake of comparison with genetic and linguistic differences(*37*, *38*, *41*, *44*). Among indigenous Taiwanese populations speaking Austronesian languages, such analyses revealed significant correlations between music, mitochondrial DNA, and the lexicon(*38*), suggesting that music may indeed preserve population history. However, whether such relationships extend beyond the level of language families remains unknown.

To address this gap, we focus on populations in and around Northeast Asia (Fig. 1). Northeast Asia provides a useful test region because it contains high levels of genetic and cultural diversity – including a large number of small language families or linguistic isolates (e.g., Tungusic, Chukuto-Kamchatkan, Eskimo-Aleut, Yukagir, Ainu, Nivkh, Korean, Japanese)(*35*). Crucially, while genetic and linguistic data throughout much of the world have been published, Northeast Asia is the only region for which published musical data allow for direct matched comparison of musical, genetic, and linguistic diversity(*41*, *45*, *46*).

**Figure 1:**
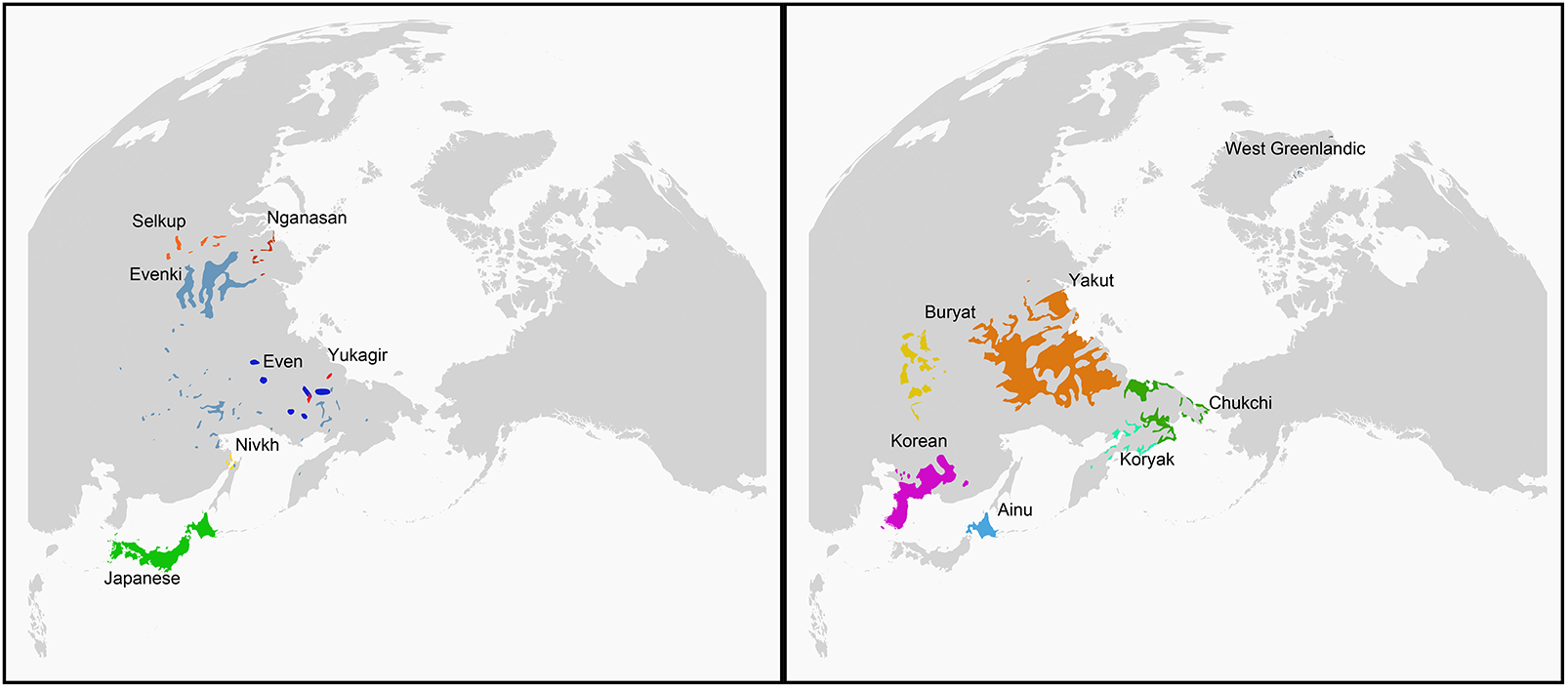
Geographical loactions of the fourteen languages. Language polygons are plotted on two panels because they partly overlap in spatial distributions. Similarly-colored pairs of languages belong to the same family: Even and Evenki belong to the Tungusic family, Selkup and Nganasan to the Uralic family, and Koryak and Chukchi to the Chukotko-Kamchatkan family.

We here use these matched comparisons to test competing hypotheses about the extent to which different forms of cultural data reflect population history at a level beyond the limits of language families. Specifically, we aim to test whether patterns of cultural evolution are significantly correlated with patterns of genetic evolution (population history), and if so, whether music or language (lexicon(*47*), grammar(*48*, *49*), or phonology(*50*–*52*)) would show the highest correlation with patterns of genetic diversity, after controlling for the influence of recent contact between languages (spatial autocorrelation) and shared inheritance within individual language families.

## Results

We selected all available populations from in and around Northeast Asia (14 populations, encompassing 11 language families/isolates) for which all four sources of data (genome-wide single nucleotide polymorphisms (SNPs), grammars, phonology, and music) were available (Fig. 1, Tables S1-S2, Materials and Methods)(*41*). For genetic data, we newly genotyped 22 Nivkh individuals from Sakhalin Island in Russia using the Illumina Human Omni 2.5-8 BeadChip array (Materials and Methods). First, we investigated the similarity between populations in each of the dimensions of enquiry. For this purpose, we used split networks(*53*), which display multiple sources of similarity in a consistent manner (Fig. 2; and Supporting Information Fig. S10-14, Tables S3-S7). Distance analysis of lexical data resulted in a network topology with an overall star-shaped structure (Fig.2C) (where we exclude Nganasan for lack of data in our source database (*47*)). Exceptions are given by the two pairs of languages that are related to one another and that stand out as proximate (Even and Evenk both belong to the Tungusic family, while Chuckchi and Koryak both belong to the Chukotko-Kamchatkan family). The results of this distance analysis are consistent with the fact that lexical material is able to detect relationships within language families, but cannot resolve historical relations between families. Therefore, we excluded the lexical data from subsequent analyses.

**Figure 2.**
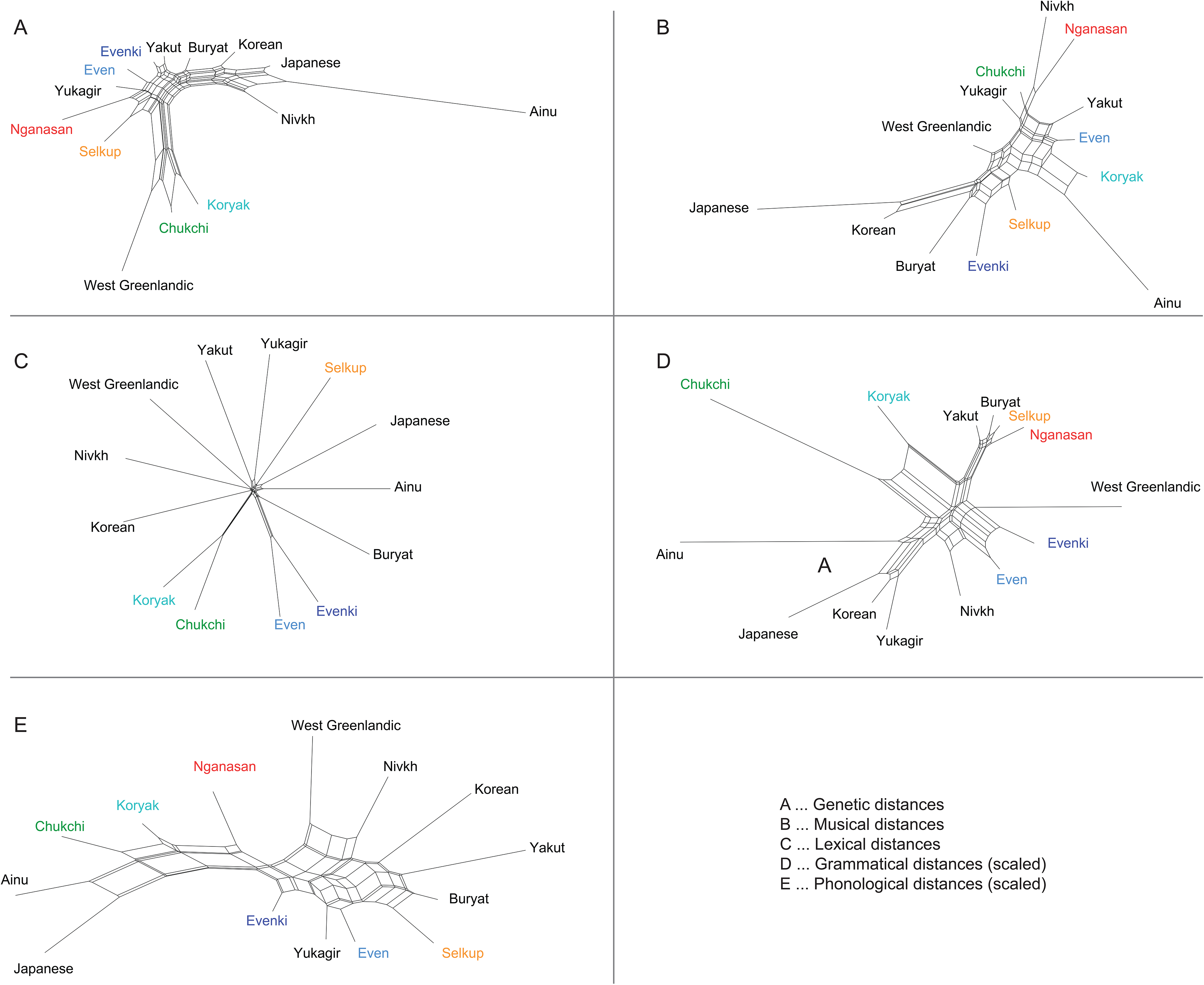
Neighbornet networks of the populations based on dimensionality-reduced distance matrices in SNPs, Lexicon, Grammar, Phonology, and Music (see Materials and Methods). Colors indicate language families: Selkup and Nganasan belong both to Uralic; Even and Evenk to Tungusic; Koryak and Chukchi to Chukotko-Kamchatkan.

Distance analyses of grammatical, phonological, genetic, and musical distances reveal potentially more informative structure. Importantly, and in agreement with the claim that language structure does not identify family relationships(*30*, *33*), the clustering emerging from the distances do not generally coincide with languages families, except perhaps for Tungusic (Even and Evenk) in the domain of grammar (with a distance of 17.37 vs. a distance of 21.45 to their next unrelated neighbor Nivkh, Table S6) and Chukotko-Kamchatkan (Chukchi and Koryak) in the domain of phonology (with a distance of 23.45 vs. a distance of 27.81 to the next unrelated neighbor Nganasan). Most of the clustering instead points to inter-family relations: for example, Korean and Japanese are neighbors in the networks based on grammar, SNPs, and music, but not phonology(*54*–*56*). Buryat and Yakut are close together in SNPs, grammar, and phonology, but not in music (*41*, *57*). The music-based network is consistent with a previous study showing the uniqueness of Ainu music and a distinction of East Asian music from circumpolar music based on cluster analysis of musical components(*41*). Nivkh shows different patterns for each factor. For example, Nivkh is genetically closer to Korean, Japanese, and Buryat than the others and shows the second highest affinity with Ainu in all populations in the distance matrix, reflecting the tree’s branch position. However, music, grammar, and phonology do not follow such relationships in Nivkh. Taken together, these results suggest that similarities in the grammatical and phonological structure are not primarily associated with simple vertical descent within language families. Instead, the different cultural features might be associated (between themselves and with genetic history) in the way in which they are transmitted, developed or adopted across families.

We implemented a redundancy analysis (RDA) on the principle components (or coordinates) of the data to explore this idea further (Materials and Methods, Supplementary Information). RDA summarizes the variation in a response variable that can be explained by an explanatory variable and finds directed associations. The RDA analysis reveals two associations that are significant under a permutation test (Fig. 3): grammatical similarity predicts genetic similarity (grammar -> genetics, variance=0.44), and genetic similarity predicts grammatical similarity (genetics -> grammar, variance=0.56).

**Figure 3:**
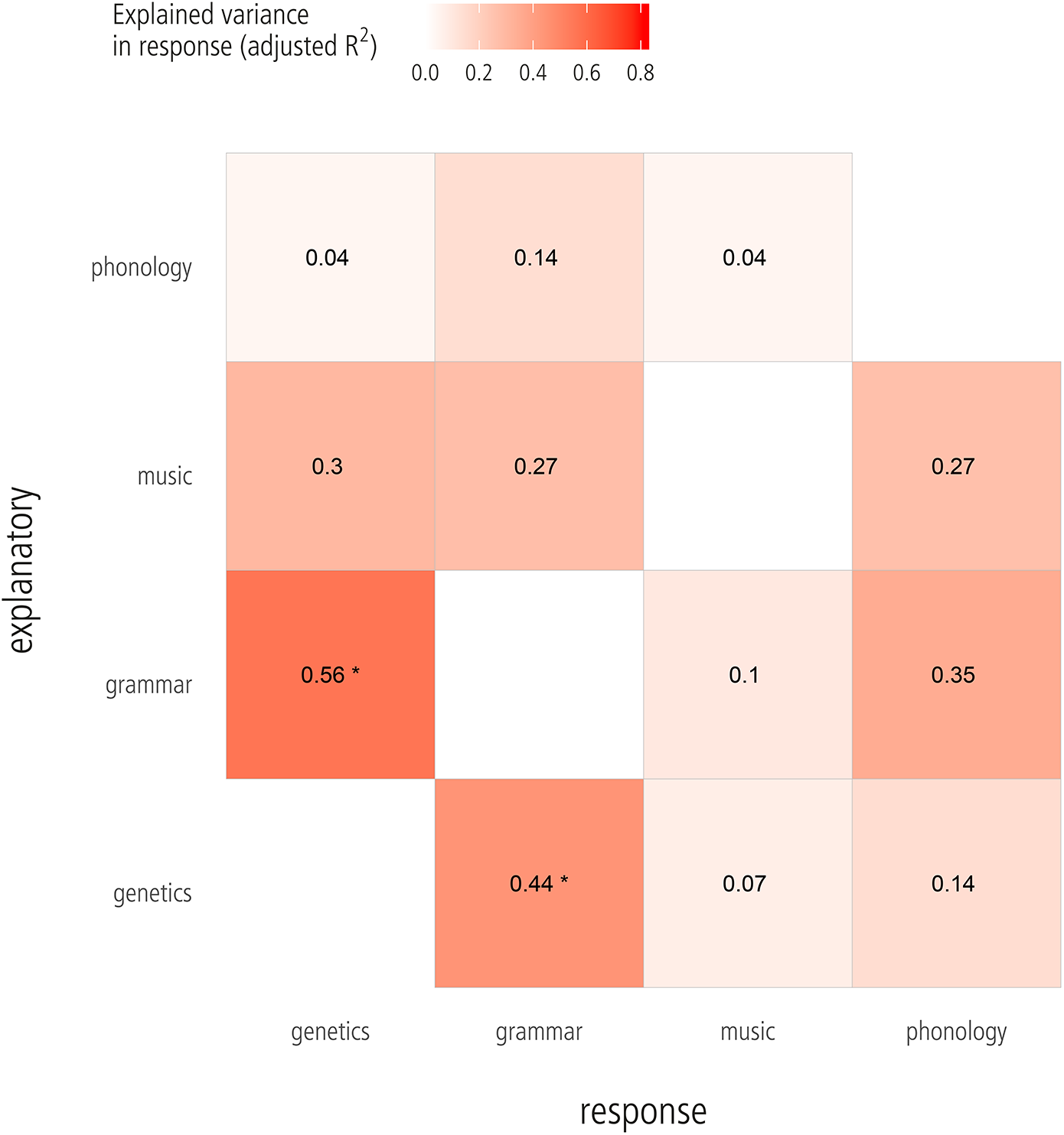
Redundancy Analysis (RDA) between four pairs of factors (genetics, grammar, music, and phonology). Variance in the response explained by each explanatory variable; * indicates a significant association (*p* ≤ 0.05).

While distance analysis suggests that both associations reflect deep-time correspondences, dating back to before the formation of current language families (as identifiably by cognate words), spatial proximity and contact between societies might lead to similar patterns of association that are relatively recent and shallow. In response to this, we evaluated three possible scenarios to explain the signal in the data: 1.) Recent contact scenario: the associations reflect recent and current contact and, hence, can be explained by spatial autocorrelation in the current data; that is, societies that are currently close to each other tend to have similar grammars and population history. 2.) Inheritance scenario: the associations reflect common ancestry. The associations result from vertical descent within the remaining linguistic families for which our sample contains more than one member (Tungusic, Chukotko-Kamchatkan, and Uralic). 3.) Deep-time correspondence scenario: the associations reflect a non-shallow correspondence between language and genetics that cannot be explained by recent contact nor phylogenetic inheritance within known families. To distinguish between the three scenarios we treated spatial proximity and inheritance as potential confounds and carried out a partial RDA to control their effect (Supplementary Information). As societies and languages placed far from the equator tend to display larger spatial ranges(*58*), we represented the territory of each society with areas rather than points and sample random spatial locations from within these areas. The partial RDA reveals strong evidence against the recent contact scenario: spatial proximity fails to explain both associations (Fig. S18). When controlling for spatial autocorrelation (1,000 random samples allowing for the uncertainty of people’s locations) the observed explained variance is still greater than that of random permutations (normalized differences between observed and permuted explained variance z > 1 SD in over 99% of spatial samples; Kullback-Leibler Divergence KLD > 2.5; Fig. S16-18). When controlling for both recent contact and phylogenetic inheritance of language in partial RDA, still both associations show stronger evidence than the other relationships (Fig. 4, and S19-21). Of them, genetics -> grammar association is stronger (z > 1 SD in 92.8% of samples, KLD > 1.5) than grammar -> genetics (z > 1 SD in 75.2% of samples; KLD = 1.1).

**Figure 4:**
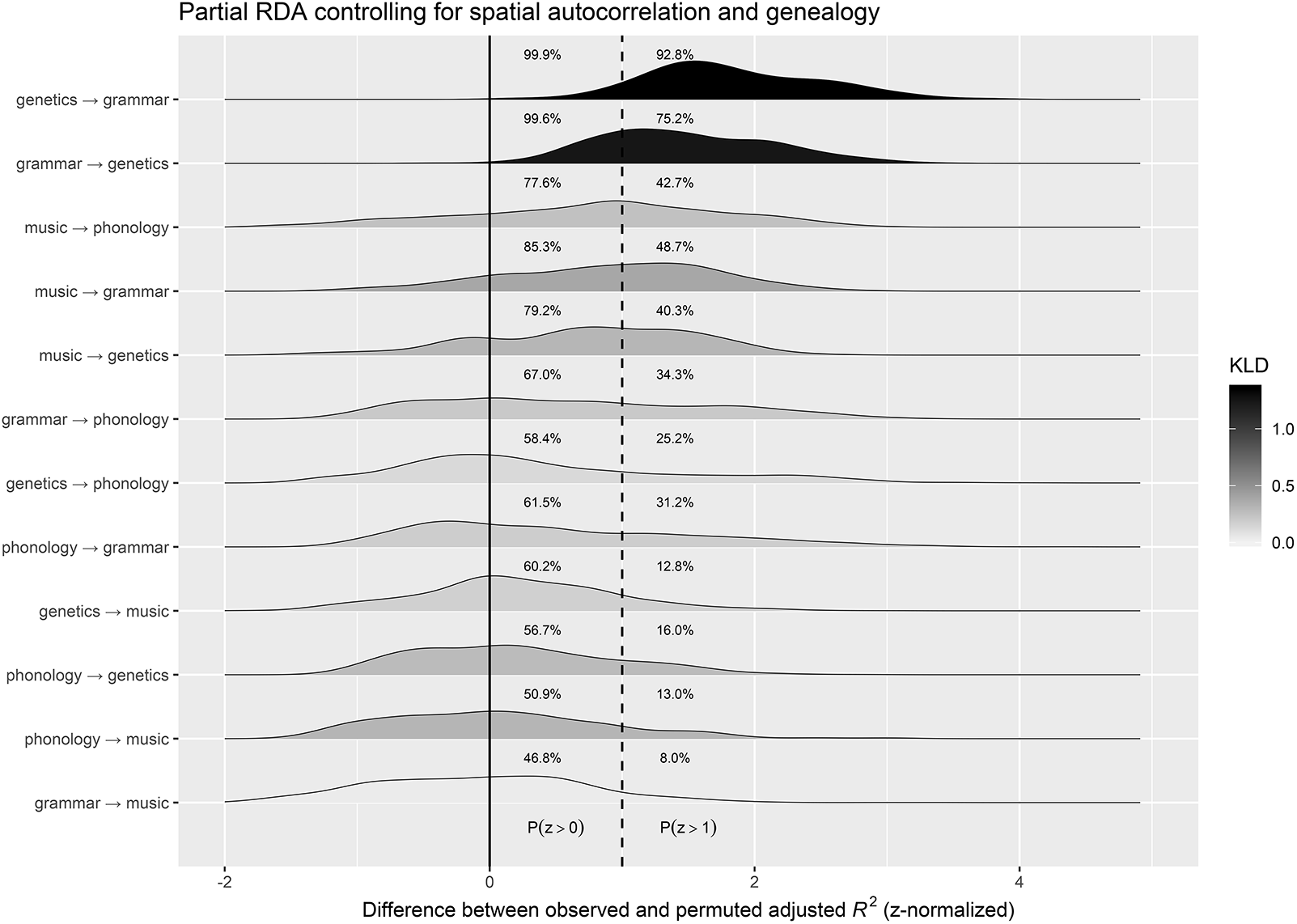
Partial RDA controlling for spatial autocorrelation and genealogy: Densities of the difference between observed and permuted adjusted *R*^2^ (z-normalized) in the partial RDA. All input components contribute at least 15% to the explained variance. Numbers right to the solid and dashed line show the proportion of simple with a positive difference (*P*(*z* > 0 SD) and a strong positive difference (*P*(*z* > 1 SD), respectively. Grey shading reflects the Kullback-Leibler divergence (KLD) between the observed and permuted adjusted *R*^2^. The larger the KLD (dark color), the more the distribution of the observed R^2^ differs from that of the random permutations. KLD=0 (transparent color) suggests that there is no difference between the distribution of the observed and the permuted R^2^.

To evaluate the robustness of the associations, we furthermore performed three types of sensitivity analyses. First, we varied the number of principal components (or coordinates) passed to the RDA and, thus, the amount of variance in both the response and the predictor. Different thresholds of how much variance a component needs to explain in order to be included (10%, 15%, 18%) show little effect on the results, although the association genetics -> grammar is far more robust (all z > 1 SD in at least 91% of samples, all KLD > 1.5) than its inverse (z > 1 SD between 74% and 99% of samples, KLD between 1.2 and 1.4; Fig. S22-S23). Second, we varied the language sample passed to the RDA. While most languages have little to no effect on the signal, this is not true for Ainu, as removing Ainu from the analysis weakens the support for the associations of grammar and genetics (z > 1 SD in only 25-31%, KLD < 0.2, when controlling for spatial proximity and inheritance; Fig. S25-S27). Third, in the partial RDA, some spatial samples happen to explain the variance in the response better than others (lower tail of observed adjusted *R*^*2*^ in Fig. S16, S17, S19, and S20). The sampled locations that produce R^2^ estimates in the 20% upper and lower tails of the total do not cluster in specific regions in the genetics -> grammar association (Fig. S28-S29), while its inverse tends towards some clustering in the polygons of Buryat, Evenki, and Selkup (Fig. S30-S31). This might be indicative of recent language contact.

To summarize, we found significant correlations between genetics and grammar by the basic RDA using the complete set of genomes, music, and language in Northeast Asia. The partial RDA controlling for geography and linguistic inheritance as well as sensitivity analyses suggest that the genetics -> grammar association is stronger than its inverse, and the relationships may trace back to earlier relationships between languages before the recent contacts and inheritance.

## Discussion

We have simultaneously explored the relations among genetic, linguistic, and musical data beyond the level of language families for the first time after Cavalli-Sforza’s works(*9*, *12*). We find remarkable evidence for the relationships between population history and grammatical similarity while genomes and grammar might be evolved by different forces.

A possible interpretation of our findings is that the relationship between grammar and population history was exceptionally well preserved over the recent contact beyond language families whether or not evolutionary mechanisms of grammar are the same as genomes. Population genetics detect gene flows between populations beyond phylogenetic relationships. Our dataset covers a phylogenetically-broad range of populations; three lineages to the present-day East Eurasian (Ainu, East Asian, and Northeast Asian) and one to North American (Greenlandic Inuit) (*59*, *60*), including gene flows beyond the lineages, such as Japanese-Ainu(*61*) and Buryat-Yakut(*57*). While the evolutionary forces that influence population history are fairly well understood, determining to what extent the genetic relationships of particular populations reflect shared ancestry vs. prehistoric contact in culture is still challenging. Moreover, the evolutionary processes that influence culture and language are under debate(*62*) but can obviously be very different from those influencing genomes. For example, cultural replacement and language shift can occur within a single generation due to colonization or other sociopolitical factors. Our results removing the influence of the proximity in cultural similarities give support to the notion that these different data reveal different historical patterns, yet show that some cultural features still can preserve relationships extending even beyond the boundaries of language families. Therefore, grammar was more conservative than the lexicon because the lexical distances are not associated with genetic distances across language families in our dataset (Fig. 2). This contrasts with expectations from historical linguistics(*33*) and also from recent findings that suggest that grammar evolves faster than the lexicon in Austronesian(*63*) and also shows rapid evolution in Indo-European(*64*). The statistical power to detect a signal is weakened when Ainu was removed in the sensitivity analysis (Fig. S24-27). While this might suggest a special position of Ainu in the larger context, we need larger samples of languages and populations inside and outside of Northeast Asia to resolve this question.

The genetics --> grammar correlation is stronger and more robust than its inverse (Fig. S15), indicating that grammatical variation preserves genetic variation better than vice versa. When controlling only for spatial autocorrelation, both are equally strong (Fig. S18), but when controlling for both spatial autocorrelation and common descent (Fig. S21), only genetics --> grammar remains robust. Furthermore, the grammar -> genetics correlation reveals more spatial clustering effects than its inverse. Together, these observations suggest that common descent and contact affect grammatical and genetic variation differently: while they can strongly affect variation in grammar, this is not the case for genes. Genetic variation is more predictable and happens - to some degree - independently of spatial neighborhood and family. This difference between grammar and genes makes it easier to predict grammatical variation from genetics than its inverse.

Interestingly, our results are qualitatively different from the only previous study to quantitatively compare genetic, linguistic, and musical relationships(*38*). Among indigenous Austronesian-speaking populations in Taiwan, music was significantly correlated with genetics but not language, while we find here that music is not robustly associated with either language or genetics. However, there are several methodological differences that might underlie these differences. In particular, the two studies looked at different types of data (genome-wide SNPs, structural linguistic features, and both group and solo songs here vs. mitochondrial DNA, lexical data, and only group songs previously). Further research with larger samples and different types of data may help to elucidate general relationships among language, music, and genetics.

The recent studies highlight Northeast Asian populations as one of major genetic components of basal East Eurasians(*23*, *65*). However, our knowledge about relationships between culture and local population history is limited in Northeast Asia. In addition to revealing an association between genetic and grammatical patterns, our results also reveal complex dissociations in which these data reflect different local histories. For example, while previous studies suggest specific genetic and cultural relationships between Korean and mainland Japanese populations(*56*, *66*) or posit a shared origin (*67*–*69*), our findings support similarities in SNPs, music, and grammar, but not in lexicon and phonology (Fig. 2, Supporting Information, Tables S3-S7). Even the Ainu show particular genetic similarity to the Japanese, their music clusters most closely with that of the Koryak (Fig. 2, Tables S3-S5). This may reflect different levels of genetic, linguistic, and musical exchange at different points of history. Musical patterns may reflect more recent cultural diffusion and gene flow from the Okhotsk and other “circumpolar” populations that interacted with the Ainu from the north within the past 1,500 years(*70*), as we previously proposed in our “triple structure” model of Japanese archipelago history(*41*). Newly genotyped Nivkh samples showed the closeness to Ainu in SNPs but not in others (Fig. 2A), suggesting that historical relationships in the coastal region of Northeast Asia. Nivkh might be a key population connecting Ainu and other Northeast Asians, however, the population history of Nivkh is not well understood. Therefore, further analyses are necessary to investigate in more detail of local population history in Northeast Asia including Nivkh.

In conclusion, we have demonstrated a relationship between grammar and genome-wide SNPs across a variety of diverse Northeast Asian language families. Our results suggest that grammatical structure may reflect population history more closely than other cultural (including lexical) data, but we also find that different aspects of genetic and cultural data reveal different aspects of our complex human histories. In other words, cultural relationships cannot be completely predicted by human population histories. Alternative interpretations of such mismatches would be historical events (e.g., language shift in local history) or culture-specific evolution independent from genetic evolution. Future analyses of such relationships at broader scales using more explicit models should help improve our understanding of the complex nature of human cultural and genetic evolution.

## Materials and Methods

### Experimental Design

#### Selection of populations in this study

We selected 14 populations for which matching musical (Cantometrics/CantoCore), genetic (genome-wide SNP) and linguistic (grammatical/phonological features) data were available (Tables S1-S2 and Fig. 1). These represented a subset of 35 Northeast Asian populations whose musical relationships were previously published and analyzed in detail(*41*). Linguistically, these 13 populations fall into 11 language families/isolates(*71*) (*72*). Korean, Ainu, Nivkh and Yukaghir are language isolates. Buryat, Japanese, Yakut, West Greenland Inuit are the sole representatives in our sample of the Mongolic, Japonic, Turkic, and Eskimo-Aleut language families, respectively. The remaining languages are classified into three language families: Koryak and Chukchi are Chukotko–Kamchatkan languages; Even and Evenk are Tungusic languages; and Selkup and Nganasan are Uralic languages.

#### Music Data

All music data and metadata are detailed in our previous report of circumpolar music(*41*). For the present analysis, we used a subset of 14 of the original 35 populations with matching genetic and linguistic data; these 14 populations are represented by 264 audio recordings of traditional songs. Each song was analyzed manually by P.E.S. using the same 41 classification characters used in (30) (from Cantometrics (29) and CantoCore (35)).

#### Genetic Data

##### Nivkh DNA samples from the Horai Collection

We used the DNAs of Nivkh maintained by the Asian DNA Repository Consortium (ADRC). The DNA samples were originally collected in Sakhalin, Russia by Dr. Satoshi Horai in the 2000s and were kept at 4℃ in Sokendai. We genotyped 32 Nivkh individuals (14 females and 18 males) with the Illumina Omni 2.5-8 BeadChip Array at the National Center for Global Health and Medicine (Table_S8_SampleID_Nivkh.xlsx). Two DNA samples were removed because of their poor quality. We selected 2,246,124 sites for SNPs with a call rate greater than 95%. Using PLINK(*73*), we performed a Hardy-Weinberg equilibrium test to exclude sites with P<10^−6^, resulting in 2,246,123 sites. Then we calculated inbreeding coefficients using sites with Minor Allele Frequency (MAF) > 0.01, confirming that none of the cousin equivalents exceeded F = 0.0625. Using the same threshold of MAF, we found kinship between 12 pairs (involving 14 individuals) with PI_HAT >0.125 (third-degree relative or closer). Eight samples were removed; 22 individuals thereby passed the quality control and kinship tests. Then we carried out strand checks between the Illumina Human Omni 2.5-8 BeadChip SNPs and JPT+CHB in 1,000 Genomes using BEAGLE 4.0(*74*). In the Nivkh data, 2,041,779 sites passed the strand check and 114,077 sites were flipped using PLINK. After the strand check, all sites that did not have an allele match were removed. We converted the Illumina unique IDs to rsIDs.

##### Merging Nivkh and public data

Publicly-available genome-wide SNP array data for 14 populations, including three Nivkh individuals (Table S1)(*56*, *75*–*79*), were obtained and curated as follows. As several genotyping platforms were used, to avoid discordancy of alleles on +/− strands we used the strand check utility in BEAGLE 4.0(*80*) for a dataset of Ainu against JPT and CHB in 1000 Genomes using BEAGLE 4.0 (*74*). To obtain shared SNPs among different platforms, genotype datasets including our Nivkh data were merged into a single dataset in PLINK file format by PLINK 1.9(*81*).

##### Removing outlier individuals

We manually removed outlier individuals from the merged dataset based on results of PCA and ADMIXTURE(*82*–*84*). Finally, we used 15 individuals of Nivkh (13 individuals from our data and 2 individuals from public data) in the population genetics analysis (Table S1, S8). The final merged genotype dataset included 245 individuals and 37,093 SNPs (total genotyping rate was 0.999). The merged dataset in PLINK format was converted to Genepop format using PGDSpider(*85*).

#### Language data

##### Lexical data

Lexical distances between populations were provided by Søren Wichmann on September 13, 2014 using version 16 of the ASJP (Automated Similarity Judgment Program) database(*47*). This program automatically calculates a matrix of pairwise distances between languages by comparing phonetic similarities across 40 categories of basic vocabulary (“Swadesh lists”). Distances are calculated as LDND (“Levenshtein Distance Normalized Divided”) values, which corrects for both differences in word lengths and the possibility of chance similarities between languages (*86*). This analysis did not attempt to identify and remove loanwords.

##### Grammar and phonology data

We combined data on grammatical and phonological traits from AUTOTYP (*52*), WALS(*48*), the ANU Phonotactics database(*50*), and PHOIBLE(*51*) and extracted a set of 21 grammar and 84 phonological features with coverage over 80% in each language, and in most cases 100% (Supporting Information Section 2).

#### Statistical Analysis

In contrast to population history, standardized methods for modeling cultural evolution across different types of data are not yet established. Therefore, we matched population history to cultural similarities to analyze both genetic and cultural data in a common framework. We obtained distance matrices representing differences between populations/languages for a subsequent comparative analysis using the following procedures for music and language, because musical and linguistic (grammatical and phonological) data have different data structures.

#### Genetic analysis

To estimate population differentiations, pairwise *F*_*st*_ values between populations were calculated with Genepop version 4.2(*87*). Pairwise *F*_*st*_ is the proportion of the total genetic variance due to between population differences, and is a convenient measure because it does not depend on the actual magnitude of the genetic variance. In other words, genetic markers that evolve slowly are expected to have the same F_st_ value as markers that evolve more rapidly, because the total variance is decomposed into within population and between population components.

#### Music analysis

A previously published matrix of pairwise distances among all 283 songs was calculated using normalized Hamming distances(*88*) to calculate the weighted average similarity across all 41 musical features(*41*). This distance matrix was then used to compute a distance matrix of pairwise musical *ϕ*_st_ values among the 14 populations using Arlequin(*89*) and the lingoes function of the ade4 package in R(*90*). φ_st_ is analogous to *F*_*st*_ but takes into account distances between individual items, making it more appropriate for analysis of cultural diversity(*44*, *88*, *90*). Further details concerning the calculations can be found elsewhere (*90*).

#### Language analysis

##### Grammar and phonology data

In contrast to songs and individual genotypes, language data do not represent individuals for each population. In view of the fact that the data are partly numerical and partly categorical, we used a balanced mix of Principle Component Analysis (PCA) and Multiple Correspondence Analysis (MCA) to calculate differences between languages (*91*) (Supplementary Information Section 3). Empty values were imputed using the R package missMDA (*92*).

#### Comparative analysis of music, SNPs, and language structure

##### PCoA for SNPs and music

We performed a principal coordinate analysis (PCoA) on the distance matrices of pairwise F_st_ for SNPs and pairwise *ϕ* _st_ for music (F_st_ and *ϕ* _st_ matrices are available from github, Supporting Information Section 3)(*93*). Similar to a PCA, a PCoA produces a set of orthogonal axes whose importance is measured by eigenvalues (Fig S2-S5). However, in contrast to the PCA, non-Euclidean distance matrices can be used. Heatplots of PCo and PC were visualized by ggplot2 in R (Fig S6-9) (*94*).

##### Split network graphs

Distances were visualized using the SplitsTree neighbornet algorithm (version 4, (*95*)) and are reported in detail in Supporting Information Tables S3-S7. In order to control for multicollinearity, we used PCA/MCAs and PCoAs as input rather than the raw data.

##### Geographic distances

The geographical polygons were taken from the Ethnologue(*96*), supplemented by a hand-drawn polygon estimate for Ainu. In view of the mobility of speakers over time, we sampled 1,000 random locations from within the polygons and used these for assessing correlations. The random point samples were generated in PostGIS https://postgis.net/ (Supporting Information Section 2.4). For each of the 1,000 samples, we computed the spherical distance between all random locations, which we store in a distance matrix. Then we perform a distance-based Moran’s eigenvector map analysis (dbMEM) to decompose the spatial structure of each of the resulting 1,000 distance matrices (Supporting Information Section 3.3)(*97*). Similar to a PCoA, dbMEM reveals the principal coordinates of the spatial locations from which the distance matrix was generated. We only return those eigenfunctions that correspond to positive spatial autocorrelation.

##### (Partial) Redundancy Analysis

Redundancy analysis (RDA) was carried out to explore the linear relationship between SNPs, grammar, phonology and music. Partial redundancy analysis (pRDA) was used to control for spatial dependence (Supporting Information Section 5). (p)RDA is an alternative to the traditionally used Mantel test, which was found to yield severely underdispersed correlation coefficients and a high false positive rate in the presence of spatially correlated data (*98*). RDA performs a regression of multiple response variables on multiple predictor variables (*99*), while pRDA also allows to control for the influence of confounders. Redundancy analysis yields an adjusted coefficient of determination (adjusted *R*^*2*^), which captures the variation in the response that can be explained by the predictors. We compare the observed adjusted *R*^*2*^ values against a distribution under random permutations (Fig. 4, Fig. S15-27). To assess robustness, we z-normalize the difference between observed and permuted adjusted *R*^*2*^ and report the proportion of samples for which the observed adjusted *R*^*2*^ is one standard deviation larger than the permuted (z > 1 SD). Moreover, we compute the Kullback-Leibler divergence (KLD) between the distribution of observed adjusted *R*^*2*^ and permuted adjusted *R*^*2*^. The KLD allows to assess the overall divergence of the two distributions, z>1 SD reports the proportion of samples with a strong positive difference. (p)RDA and subsequent analyses were performed in R using the vegan package (64).

## General

We thank the Asian DNA Repository Consortium for the use of Ainu SNPs data and URPPs Evolution in Action and Language and Space University of Zurich for support for an interdisciplinary workshop “Frontiers of early human expansion in Asia: linguistic and genetic perspectives on Ainu, Japan and the North Pacific Rim” to help study linguistic and genetic histories in North Asia. Computations were partially performed on the NIG supercomputer at ROIS National Institute of Genetics, Japan. We thank Søren Wichmann for providing the ASJP distance matrix and Brigitte Pakendorf, Johanna Nichols, Juha Janhunen, Ekaterina Gruzdeva, Anna Bugaeva, Anna Berge, Dagmar Jung, and Marcelo Sánchez, and Chiara Barbieri for discussion.

## Data and material availability

Data and analysis code are available at https://github.com/derpetermann/music_languages_genes. All raw data were previously published and can be accessed from the original publications and/or public data repositories, with the exception of the Ainu and Nivkh DNA data which was provided by the Asian DNA Repository Consortium. All data and analysis code needed to evaluate the conclusions in the paper are present in the paper and/or the Supplementary Materials.

## Author Contributions

P.E.S., H.M., H.O., B.B. and K.K.S. initially designed the research with advice from M.S. and S.B.. K.K. N.N. and H.T. genotyped Nivkh DNA samples. T.S. A.T. H.O. and H.M. analyzed genetic data. P.R., D.E.B., B.B., and T.E.C. analyzed language data. P.E.S., H.M. and P.R. analyzed music data. H.M., D.E.B, P.R, and B.B designed and implemented the statistical analysis. H.M, P.E.S., P.R., M.S., B.B., K.K.S., and D.E.B. wrote the manuscript. P.R., D.E.B. and B.B wrote the supporting information.

## Figure Legends

Figure 1. Geographic areas of 14 languages/populations. Because some of the areas overlap in space, they are plotted in two separate maps.

Figure 2. Neighbornet networks of the populations based on dimensionality-reduced distance matrices in SNPs, Lexicon, Grammar, Phonology, and Music (see Materials and Methods). Colors indicate language families: Selkup and Nganasan belong both to Uralic; Even and Evenk to Tungusic; Koryak and Chukchi to Chukotko-Kamchatkan.

Figure 3. Redundancy Analysis (RDA) between four pairs of factors (genetics, grammar, music, and phonology). Variance in the response explained by each explanatory variable; * indicates a significant association (p ≦ 0.05).

Figure 4. The larger the KLD (dark color), the more the distribution of the observed R^2^ differs from that of the random permutations. KLD=0 (transparent color) suggests that there is no difference between the distribution of the observed and the permuted R^2^.

