## Supplementary Information for "Exploring correlations in genetic and cultural variation across language families in Northeast Asia"

November 11, 2020

**References** **59**

### S1 Packages and functions

#### S1.1 Packages

For the analysis we use the following three main packages:

- `ade4` provides tools for multivariate data analysis
- `adespatial` provides tools for multiscale spatial analysis of multivariate data
- `vegan` provides tools for ordination methods and diversity analysis
- `FactoMinor` provides tools for dimensionality reduction of mixed data (categorical and continuous)

```r
# Data analysis (main)
library(ade4)
library(vegan)
library(adespatial)
library(FactoMineR)

# Data analysis (additional)
library(missMDA)
library(ape)
library(LaplacesDemon)
library(phangorn)

# Data handling and manipulation
library(dendextend)
library(broom)
library(reshape2)
library(plyr)
library(dplyr)

# Plotting and knitting
library(knitr)
library(kableExtra)
library(ggplot2)
library(ggpolypath)
library(cowplot)
library(ggridges)
library(RColorBrewer)
library(scales)

# Spatial Analysis and mapping
library(sp)
library(spdep)
library(rgdal)
library(rgeos)
library(mapproj)
```

#### S1.2 Functions

In this section we document all custom-defined functions for performing the redundancy analysis and for data preprocessing.

##### S1.2.1 Redundancy Analysis

```r
# Distance-based Moran's Eigenvector (dbMEM) Analysis
random_points_to_dbmem <- function(r_points, print_count=FALSE){
#' Computes dbMEMs for each random point sample in r_points
#' @param r_points: the random points (SpatialPointsDataFrame)
#' @param print_count: Print the number of processed samples when iterating over r_points?
#' @return a list comprising the dbMEMs for each sample

  if (class(r_points)[1] != "SpatialPointsDataFrame") {
    stop("please provide a SpatialPointsDataFrame")
  }
  n_sample <- unique(r_points$sample_id)
  #n_sample <- max(r_points$sample_id)

  epsilon <- 0.1
  geo_pco  <- list()

  for (j in n_sample) {

      # Get all points of sample j
      points <- r_points[r_points$sample_id == j,]

      # Compute distances between all points in the sample
      mat <- spDists(points, points)

      # Compute the mst, find its longest edge and use as a threshold
      mst_1 <- spantree(mat)
      mst_le <- max(mst_1$dist)

      # Add a small epsilon (for numerical stability)
      thresh <- mst_le + epsilon

      # Find all nearest neighbors within the distance threshold
      nb <- dnearneigh(points, 0, thresh)

      # Normalize the data
      spwt <- lapply(nbdists(nb, points), function(x) 1 - (x/(4 * thresh))^2)

      # Compute weighted neighbor list
      lw <- nb2listw(nb, style = "B", glist = spwt, zero.policy = TRUE)

      # Compute MEMs with a corresponding positive autocorrelation
      res <- as.data.frame(scores.listw(lw, MEM.autocor = "positive"))

      rownames(res) <- points$nam_label
      colnames(res) <- paste("geo_pco_", seq(1,ncol(res)), sep="")
      res <- res[order(rownames(res)), drop = FALSE, ]
      geo_pco[[paste('sample_', j, sep="")]] <- res

      if (print_count) {
        if (j%%1000 == 0) {
          print(paste(j, " samples processed"))}}}
```

```r
  return (geo_pco)}

# Redundancy Analysis

rda_wrapper_setup_0 <- function(response, explanatory, n_perm){
#' Performs an RDA without controlling for confounding factors
#' @param response: the repsonse variable
#' @param explanatory: the explanatory variable
#' @param n_perm number of permutations for anova-like permutation tes

  if (is.null(rownames(response)) & is.null(rownames(explanatory))) {
    stop("Row names of the response and the explanatory variable must be defined!")
  }

  if (any(rownames(response) != rownames(explanatory))) {
    stop("Row names of the response and the
         explanatory variable must be identical and match in order!")
  }

  ex_name <- sub('\\_.*', '', colnames(explanatory)[1])
  re_name <- sub('\\_.*', '', colnames(response)[1])

  # Run the RDA (in vegan package explanatory variables are Y!)
  rda <- rda(Y = explanatory, X = response)

  # Compute the adjusted explained variance and the significance
  r2 <- RsquareAdj(rda)$r.squared
  r2_adj <- RsquareAdj(rda)$adj.r.squared
  sig <- anova.cca(rda, step = n_perm)$`Pr(>F)`[1]
  rda_results <- list(r2=r2, r2_adj=r2_adj, sig=sig,
                      explanatory=ex_name, response=re_name)}

rda_wrapper_setup_1 <- function (response, explanatory, random_geo_pco=NULL,
                                 n_perm=100, print_count=FALSE) {
#' Performs a partial RDA and controls for spatial autocorrelation
#' @param response: the repsonse variable
#' @param explanatory: the explanatory variable
#' @param random_geo_pco dbMEMs of random spatial point patterns
#' @param n_perm number of permutations per random geo sample
#' @param print_count: print the number of processed samples when iterating over r_points?
#' @return a list comprising the rda results for each sample

  if (is.null(rownames(response)) & is.null(rownames(explanatory))) {
    stop("Row names of the response and the explanatory variable must be defined!")
  }

  if (any(rownames(response) != rownames(explanatory))) {
    stop("Row names of the response and the
         explanatory variable must be identical and match in order!")
  }

  ex_name <- sub('\\_.*', '', colnames(explanatory)[1])
```

```r
  re_name <- sub('\\_.*', '', colnames(response)[1])

  rda_results <- list()
  progress = 0

  for (sample in names(geo_mem)) {
    progress = progress + 1
    constraint <- random_geo_pco[[sample]]

    # Run the RDA (in vegan package explanatory variables are Y!)
    rda <- rda(Y = explanatory, X = response, Z = constraint)

    # Compute the (adjusted) explained variance
    r2_res = RsquareAdj2_part(rda)
    r2_semi <- r2_res$r_squared_a
    r2_partial <- r2_res$r_squared_b
    r2_adj_semi <- r2_res$adj_r_squared_a
    r2_adj_partial <- r2_res$adj_r_squared_b

    # Perform permutations
    perm <- permute_rda(n_perm, explanatory, response, constraint)

    rda_results[[sample]] <- list(r2_semi=r2_semi, r2_adj_semi=r2_adj_semi,
                                  r2_partial=r2_partial, r2_adj_partial=r2_adj_partial,
                                  perm_r2_semi=perm$r2_semi,
                                  perm_r2_adj_semi=perm$r2_adj_semi,
                                  perm_r2_partial=perm$r2_partial,
                                  perm_r2_adj_partial=perm$r2_adj_partial,
                                  explanatory=ex_name, response=re_name, geo=TRUE)}
    if (print_count){if (i%%10 == 0) {print(paste(i, " samples processed"))}}

  return(rda_results)}

rda_wrapper_setup_2 <- function (response, explanatory, random_geo_pco=NULL,
                                 n_perm=100, print_count=FALSE) {
#' Performs a partial RDA and controls for spatial autocorrelation  and genealogy
#' @param response: the repsonse variable
#' @param explanatory: the explanatory variable
#' @param random_geo_pco dbMEMs of random spatial point patterns
#' @param n_perm number of permutations per random geo sample
#' @param print_count: print the number of processed samples when iterating over r_points?
#' @return a list comprising the rda results for each sample

  if (is.null(rownames(response)) & is.null(rownames(explanatory))) {
    stop("Row names of the response and the explanatory variable must be defined!")
  }

  if (any(rownames(response) != rownames(explanatory))) {
    stop("Row names of the response and the
         explanatory variable must be identical and match in order!")
  }

  ex_name <- sub('\\_.*', '', colnames(explanatory)[1])
```

```r
  re_name <- sub('\\_.*', '', colnames(response)[1])

  rda_results <- list()
  progress = 0

  for (sample in names(geo_mem)) {
    progress = progress + 1

    constraint <- random_geo_pco[[sample]]

    #  Sample one language from each language family with two members
    sample_lgs <- c(single_lgs, sample(uralic,1), sample(chkkat, 1), sample(tungus, 1))
    explanatory_sample = explanatory[rownames(explanatory) %in% sample_lgs, , drop=F]
    response_sample = response[rownames(response) %in% sample_lgs, , drop=F]
    constraint_sample = constraint[rownames(constraint) %in% sample_lgs, ,drop=F]

    # Run the RDA (in vegan package explanatory variables are Y!)
    rda <- rda(Y = explanatory_sample, X = response_sample,
               Z = constraint_sample)

    # Compute the (adjusted) explained variance
    r2_res = RsquareAdj2_part(rda)

    r2_semi <- r2_res$r_squared_a
    r2_partial <- r2_res$r_squared_b
    r2_adj_semi <- r2_res$adj_r_squared_a
    r2_adj_partial <- r2_res$adj_r_squared_b

    # Perform permutations
    perm <- permute_rda(n_perm, explanatory_sample, response_sample, constraint_sample)

    rda_results[[sample]] <- list(r2_semi=r2_semi, r2_adj_semi=r2_adj_semi,
                                  r2_partial=r2_partial, r2_adj_partial=r2_adj_partial,
                                  perm_r2_semi=perm$r2_semi,
                                  perm_r2_adj_semi=perm$r2_adj_semi,
                                  perm_r2_partial=perm$r2_partial,
                                  perm_r2_adj_partial=perm$r2_adj_partial,
                                  explanatory=ex_name, response=re_name, geo=TRUE)}
   if (print_count){
    if (i%%100 == 0) {
      print(paste(i, " samples processed"))}}
 return(rda_results)}

rda_wrapper_sensitivity_1 <- function (response, explanatory, random_geo_pco=NULL, exclude_site,
                                       n_perm=100, print_count=FALSE) {
#' Performs a sensitivity analysis for RDA and controlling for spatial autocorrelation
#' @param response: the repsonse variable
#' @param explanatory: the explanatory variable
#' @param random_geo_pco dbMEMs of random spatial point patterns
#' @param exclude_site Name of the society that is excluded from the analysis
#' @param n_perm number of permutations per random geo sample
#' @param print_count: print the number of processed samples when iterating over r_points?
```

```r
#' @return a list comprising the rda results for each sample

  if (is.null(rownames(response)) & is.null(rownames(explanatory))) {
    stop("Row names of the response and the explanatory variable must be defined!")
  }

  if (any(rownames(response) != rownames(explanatory))) {
    stop("Row names of the response and the
         explanatory variable must be identical and match in order!")
  }

  ex_name <- sub('\\_.*', '', colnames(explanatory)[1])
  re_name <- sub('\\_.*', '', colnames(response)[1])

  rda_results <- list()
  progress = 0

  for (sample in names(geo_mem)) {
    progress = progress + 1

    constraint <- random_geo_pco[[sample]]

    # Exclude one society: the society might hide in each of the families or the isolates
    single_lgs_ex <- single_lgs[single_lgs!=exclude_site]
    uralic_ex <- uralic[uralic!=exclude_site]
    chkkat_ex <- chkkat[chkkat!=exclude_site]
    tungus_ex <- tungus[tungus!=exclude_site]

    #  Samples: only use societies that are not excluded
    sample_lgs <- c(single_lgs_ex, uralic_ex, chkkat_ex, tungus_ex)
    explanatory_sample = explanatory[rownames(explanatory) %in% sample_lgs, , drop=F]
    response_sample = response[rownames(response) %in% sample_lgs, , drop=F]
    constraint_sample = constraint[rownames(constraint) %in% sample_lgs, ,drop=F]

    # Run the RDA (in vegan package explanatory variables are Y!)
    rda <- rda(Y = explanatory_sample, X = response_sample,
               Z = constraint_sample)

    # Compute the (adjusted) explained variance
    r2_res = RsquareAdj2_part(rda)

    r2_semi <- r2_res$r_squared_a
    r2_partial <- r2_res$r_squared_b
    r2_adj_semi <- r2_res$adj_r_squared_a
    r2_adj_partial <- r2_res$adj_r_squared_b

    # Perform permutations
    perm <- permute_rda(n_perm, explanatory_sample, response_sample, constraint_sample)

    rda_results[[sample]] <- list(r2_semi=r2_semi, r2_adj_semi=r2_adj_semi,
                                  r2_partial=r2_partial, r2_adj_partial=r2_adj_partial,
                                  perm_r2_semi=perm$r2_semi,
                                  perm_r2_adj_semi=perm$r2_adj_semi,
```

```
                                    perm_r2_partial=perm$r2_partial,
                                    perm_r2_adj_partial=perm$r2_adj_partial,
                                    explanatory=ex_name, response=re_name,
                                    n_sites=nrow(explanatory_sample))}
   if (print_count){
    if (i%%100 == 0) {
      print(paste(i, " samples processed"))}}
 return(rda_results)}

rda_wrapper_sensitivity_2 <- function (response, explanatory, random_geo_pco=NULL, exclude_site,
                                          n_perm=100, print_count=FALSE) {
#' Performs a sensitivity analysis for partial RDA
#' controlling for spatial autocorrelation and genealogy
#' @param response: the repsonse variable
#' @param explanatory: the explanatory variable
#' @param random_geo_pco dbMEMs of random spatial point patterns
#' @param exclude_site Name of the society that is excluded from the analysis
#' @param n_perm number of permutations per random geo sample
#' @param print_count: print the number of processed samples when iterating over r_points?
#' @return a list comprising the rda results for each sample

  if (is.null(rownames(response)) & is.null(rownames(explanatory))) {
    stop("Row names of the response and the explanatory variable must be defined!")
  }

  if (any(rownames(response) != rownames(explanatory))) {
    stop("Row names of the response and the
         explanatory variable must be identical and match in order!")
  }

  ex_name <- sub('\\_.*', '', colnames(explanatory)[1])
  re_name <- sub('\\_.*', '', colnames(response)[1])

  rda_results <- list()
  progress = 0

  for (sample in names(geo_mem)) {
    progress = progress + 1

    constraint <- random_geo_pco[[sample]]

    # Exclude one society: the society might hide in each of the families or the isolates
    single_lgs_ex <- single_lgs[single_lgs!=exclude_site]
    uralic_ex <- uralic[uralic!=exclude_site]
    chkkat_ex <- chkkat[chkkat!=exclude_site]
    tungus_ex <- tungus[tungus!=exclude_site]

    #  Sample one language from each language family with two members
    sample_lgs <- c(single_lgs_ex, sample(uralic_ex,1), sample(chkkat_ex, 1),
                    sample(tungus_ex, 1))
    explanatory_sample = explanatory[rownames(explanatory) %in% sample_lgs, , drop=F]
    response_sample = response[rownames(response) %in% sample_lgs, , drop=F]
```

```r
    constraint_sample = constraint[rownames(constraint) %in% sample_lgs, ,drop=F]

    # Run the RDA (in vegan package explanatory variables are Y!)
    rda <- rda(Y = explanatory_sample, X = response_sample,
               Z = constraint_sample)

    # Compute the (adjusted) explained variance
    r2_res = RsquareAdj2_part(rda)

    r2_semi <- r2_res$r_squared_a
    r2_partial <- r2_res$r_squared_b
    r2_adj_semi <- r2_res$adj_r_squared_a
    r2_adj_partial <- r2_res$adj_r_squared_b

    # Perform permutations
    perm <- permute_rda(n_perm, explanatory_sample, response_sample, constraint_sample)

    rda_results[[sample]] <- list(r2_semi=r2_semi, r2_adj_semi=r2_adj_semi,
                                  r2_partial=r2_partial, r2_adj_partial=r2_adj_partial,
                                  perm_r2_semi=perm$r2_semi,
                                  perm_r2_adj_semi=perm$r2_adj_semi,
                                  perm_r2_partial=perm$r2_partial,
                                  perm_r2_adj_partial=perm$r2_adj_partial,
                                  explanatory=ex_name, response=re_name,
                                  n_sites=nrow(explanatory_sample))}
   if (print_count){
    if (i%%100 == 0) {
      print(paste(i, " samples processed"))}}
 return(rda_results)}

# Permute RDA
permute_rda <- function(n_perm, explanatory, response, constraint=NULL) {
#' Runs n_perm RDAs with permuted data
#' @param n_perm: the number of permutations
#' @param response: the repsonse variable
#' @param explanatory: the explanatory variable
#' @param constraint the constraint
#' @return a list comprising the RDA results for each permutation

  if (!is.numeric(n_perm)){stop("The number of permutations must
                                be .. well... a number.")}
  permutation_results <- data.frame(r2_adj=rep(NA, n_perm))

  for (i in 1:n_perm){
    # permute the response
    permutation_order <- sample(1:nrow(response))
    response <- response[permutation_order, , drop=F]

    # Geo-constraint?
    # Run the RDA (in vegan package explanatory variables are Y!)
    if (is.null(constraint)) {rda <- rda(Y = explanatory, X = response)}
    else {rda <- rda(Y = explanatory, X = response, Z = constraint)}
```

```r
    # Compute the (adjusted) explained variance
    r2_res = RsquareAdj2_part(rda)

    r2_semi <- r2_res$r_squared_a
    r2_partial <- r2_res$r_squared_b
    r2_adj_semi <- r2_res$adj_r_squared_a
    r2_adj_partial <- r2_res$adj_r_squared_b

    permutation_results[i, c("r2_semi")] <- r2_semi
    permutation_results[i, c("r2_partial")] <- r2_partial
    permutation_results[i, c("r2_adj_semi")] <- r2_adj_semi
    permutation_results[i, c("r2_adj_partial")] <- r2_adj_partial}

  return(permutation_results)}

# Compute adjusted R-squared
RsquareAdj2_part <- function (x) {

  #' Computes the (adjusted) R-squared of an RDA model using either
  #' - vegan's semipartial method
  #' - CONOCO's partial method
  #' code adapted from:
  #' https://davidzeleny.net/blog/2016/09/08/adjusted-r2-in-partial-constrained-
  #' ordination-the-difference-between-r-vegan-and-canoco-5/
  #' @param x: RDA result
  #' @return a list comprising the R-squared / adjusted R-squared

  m <- x$CCA$qrank
  n <- nrow(x$CCA$u)
  R2_a <- x$CCA$tot.chi/x$tot.chi
  R2p_a <- x$pCCA$tot.chi/x$tot.chi
  p_a <- x$pCCA$rank
  radj_a <- RsquareAdj(R2_a + R2p_a, n, m + p_a) - RsquareAdj(R2p_a, n, p_a)

  R2_b <- x$CCA$tot.chi/(x$tot.chi - x$pCCA$tot.chi)
  p_b <- x$pCCA$rank
  radj_b <- 1 - (1 - R2_b)*(n - p_b - 1)/(n - m - p_b - 1)

  if (any(na <- m >= n - 1)) radj_b[na] <- NA

  return (list(r_squared_a = R2_a, adj_r_squared_a = radj_a, r_squared_b = R2_b,
               adj_r_squared_b = radj_b))}
```

##### S1.2.2 Helper functions for data preprocessing

```r
trim_data <- function(data.list, trim.to=siberia_metadata_all, extra.coverage=.8) {
 #' Trims the linguistic input data and only keeps variables with
 #' data points for languages in the study area and with non-constant values

 #' @param data.list the data to be trimmed
 #' @param trim.to contains the ids of those languages that are retained after trimming
```

```r
 #' @param extra.coverage the percentage of covered data
 #' @return the trimmed data

  lgs <- lapply(data.list, function(l) {
      l$UULID <- ifelse(l$isocode %in% c('bxm','bxr'),
               '[i-bua][a-1095][g-buri1258]',
              paste(l$UULID))
      subset(l, UULID %in% trim.to$UULID)
     })
  vars <- lgs[sapply(lgs, function(l) {
       length(l$UULID)==length(unique(l$UULID)) &
       length(unique(l[,1]))>1 &
         length(unique(l$UULID)) >= floor(extra.coverage*length(trim.to$UULID))
      })]
  return(lapply(vars, function(l) l[,c(1,3)]))}

compute_coverage <- function(data.list, gg=siberia_metadata) {
 #' Computes the coverage of all languages
 #' @param data.list the input data
 #' @param gg a data.frame comprising the languages for which the coverage is computed
 #' @return the input data with the coverage added as a separate column

  x <- sapply(gg$UULID, function(l) {
         coverage <- round(mean(sapply(data.list, function(v) { l %in% v$UULID })),2)*100
        })
  df <- data.frame(UULID=names(x), Coverage=x)
  gg$Language <- rownames(gg)
  df.g <- merge(df, gg)
  return(df.g)}

flatten <- function(data.list, gg=siberia_metadata) {
 #' Flattens the nested linguistic data
 #' @param data.list: the input data
 #' @param gg: a data.frame comprising the languages for which the data are flattened
 #' @return the flattened data

  df.list <- lapply(seq_along(data.list), function(v) {
    df <- data.list[[v]]
    var.name <- gsub('(.*\\$)', '\\2', names(data.list)[v])
    names(df)[1] <- var.name
    return(dplyr::select(df, UULID, dplyr::everything()))
    })
  df.flat <- Reduce(function(x,y) dplyr::full_join(x, y, by='UULID'), df.list)
  rownames(df.flat) <- sapply(df.flat$UULID, function(x) rownames(gg[gg$UULID %in% x,]))
  return(df.flat)}

print_dist <- function(d, caption) {
  #' Prints the distance matrices in a table
  #' @param d: the input distance matrix
  #' @param caption: the caption of the table
```

```r
  d.m <- as.matrix(sort_dist_mat(d))
  d.m[upper.tri(d.m, diag=T)] <- NA
  colnames(d.m) <- abbreviate(colnames(d.m), minlength=8)
  rownames(d.m) <- abbreviate(rownames(d.m), minlength=8)
  options(knitr.kable.NA = '')
  d.m <- d.m[2:nrow(d.m),1:ncol(d.m)-1]

  kable(d.m, digits=3, format = 'latex', caption=caption) %>%
    kable_styling(latex_options = c("scale_down", "hold_position")) %>%
                  column_spec(1, border_left=T) %>%
                  column_spec(ncol(d.m)+1, border_right=T)}

pcos_to_factors <- function(var_th, genetics, genetics_ev, grammar, grammar_ev,
                            phonology, phonology_ev, music, music_ev, geo){
  #' Retains k PCs/PCos which account for at least var_th percent
  #' of the explained variance
  #' @param var_th: the variance threshold for each PC/PCo
  #' @param genetics: the PCos for genetics
  #' @param genetics_ev: contains information on the explained variance for genetics
  #' @param grammar: the PCs for grammar
  #' @param grammar_ev: contains information on the explained variance for grammar
  #' @param phonology: the PCs for phonology
  #' @param phonology_ev: contains information on the explained variance for phonology
  #' @param music: the PCos for music
  #' @param music_ev: contains information on the explained variance for music
  #' @param geo: the dbMEMs
  #' @return a list comprising all factors

  genetics_pco_rel <- genetics[, which(genetics_ev >= var_th)]
  grammar_pc_rel <- grammar[, which(grammar_ev >= var_th*100)]
  phonology_pc_rel <- phonology[, which(phonology_ev >= var_th*100), drop=F]
  music_pco_rel <- music[, which(music_ev >= var_th)]

  factors = list(genetics = genetics_pco_rel[order(rownames(genetics_pco_rel)), ],
                 music = music_pco_rel[order(rownames(music_pco_rel)), ],
                 grammar = grammar_pc_rel[order(rownames(grammar_pc_rel)), ],
                 phonology = phonology_pc_rel[order(rownames(phonology_pc_rel)), ,drop=F],
                 geo = geo)}

get_all_factor_combinations <- function(factor_name){
  #' This helper function create a data frame from all combinations of factor names
  #' @param factor_names: the names of all factors
  #' @return a data frame with all combinations of factor names

  # All possible combinations
  all_comb <- expand.grid(factor_names, factor_names)
  comb <- all_comb[!all_comb$Var1 == all_comb$Var2, ]
  comb <- as.data.frame(t(comb), stringsAsFactors=FALSE)

  return(comb)}
```

```r
rda_to_correlation_matrix <- function(rda, adjust_significance=TRUE){
  #' This helper function turns the results from an RDA into a correlation matrix
  #' Moreover, it adjusts the significance values using False Discovery Rate (FDR)
  #' @param rda: the RDA results
  #' @param adjust_significance: Use FDR to adjust significance values?

  # Simplify RDA results
  rda_mat <- sapply(rda, function(x){
                simp <- c(r2=x$r2, r2_adj=x$r2_adj, sig=x$sig,
                          explanatory=x$explanatory, response=x$response)
                return(simp)})

  rda_mat <- data.frame(t(rda_mat))
  rda_mat <- transform(rda_mat, sig=as.numeric(as.character(sig)),
             r2=as.numeric(as.character(r2)), r2_adj=as.numeric(as.character(r2_adj)))

  # Adjusting significance (False discovery rate)

  if (adjust_significance==T)
    rda_mat$sig <- p.adjust(rda_mat$sig, method = "fdr")

  # Significance
  rda_mat[rda_mat$sig > 0.05, "sig_level"] <- ""
  rda_mat[rda_mat$sig <= 0.01 , "sig_level"] <- "**"
  rda_mat[rda_mat$sig > 0.01 & rda_mat$sig <= 0.05 , "sig_level"] <- "*"

  return(rda_mat)}

partial_rda_to_z_val <- function(rda_result, r2_type) {
  #' Flattens the partial RDA results and computes the z-value for the
  #' difference between the observed and permuted R-squares.
  #' Then retrieves the probability density distribution
  #' for both the observed and permuted R-squares.
  #' @param rda_results: the results from a partial RDA analysis over different paramaters
  #' @param r_2_type: the R-squared type that should be retained (r2_semi, r2_adj_semi,
  #'                  r2_partial, r2_adj_partial)
  #' @param n_lang: include the number of societies in the ouput data frame
  #' @return a list comprising the flattened values, z-standardized values and densities

  perm_type = paste("perm_", r2_type, sep="")

  # Difference between observed and permuted R-squares
  rda_diff <- lapply(rda_result, function(x)
    unlist(sapply(x, function(y)
      y[[r2_type]] - mean(y[[perm_type]]))))

  rda_diff_df <- ldply(rda_diff, data.frame)
  colnames(rda_diff_df) <- c("Association", "Values")

  # Z-normalize difference between permuted and observed R-squares
  rda_diff_z <- lapply(rda_result, function(x)
    unlist(sapply(x, function(y)
    (y[[r2_type]] - mean(y[[perm_type]]))/sd(y[[perm_type]]))))
```

```r
  rda_diff_z_df <- ldply(rda_diff_z, data.frame)
  colnames(rda_diff_z_df) <- c("Association", "z")

  # Observed
  rda_obs <- lapply(rda_result, function(x)
    unlist(sapply(x, function(y)
    y[[r2_type]])))

  rda_obs_df <- ldply(rda_diff_z, data.frame)
  colnames(rda_obs_df) <- c("Association", "Values")

  # Permuted
  rda_perm <- lapply(rda_result, function(x)
    unlist(sapply(x, function(y)
      sample(y[[perm_type]], 1))))

  rda_perm_df <- ldply(rda_perm, data.frame)
  colnames(rda_perm_df) <- c("Association", "Values")

  # Probability densities
  densities <- lapply(names(rda_obs), function(x) {

    # Get min and max values for both distributions
    min <- min(c(rda_obs[[x]], rda_perm[[x]]))
    max <- max(c(rda_obs[[x]], rda_perm[[x]]))

    # Estimate densities
    obs_d <- density(rda_obs[[x]], bw = "nrd", from=min, to=max)
    perm_d <- density(rda_perm[[x]], bw = "nrd", from=min, to=max)

    return (list(observed=obs_d,
                 permuted=perm_d))})

  names(densities) <- names(rda_obs)

  return(list(val=rda_diff_df, z_val=rda_diff_z_df, densities=densities))}

add_proportion_and_kld <- function(rda_flat, diff_th){
  #' Adds the proportions of samples larger than 0 and 1
  #' and the Kullback Leibler divergence (KLD) to the flattened RDA results
  #' @param rda_flat: the flattened RDA results
  #' @param diff_th: the threshold value
  #' @return a list comprising the flattened rda results (with KLD and proportions)
  #' and a summary of the same results (both are needed for plotting)

  # Which proportion of points is larger than zero?
  proportion_pos <- sapply(unique(rda_flat$z_val$Association), function(x) {

    sum(rda_flat$z_val$z[rda_flat$z_val$Association==x] >= 0.0)/
      sum(rda_flat$z_val$Association==x) })
```

```r
# Which proportion of points is larger than diff_th?
proportion_th <- sapply(unique(rda_flat$z_val$Association), function(x) {

  sum(rda_flat$z_val$z[rda_flat$z_val$Association==x] >= diff_th)/
    sum(rda_flat$z_val$Association==x) })

# Compute Kullback-Leibler divergence from the observed (px)
# to the permuted (py) distribution
kld <- sapply(rda_flat$densities, function (a) {
   kld_results <- KLD(px=a[["observed"]]$y,
                      py=a[["permuted"]]$y)
   div_obs_perm <- kld_results$sum.KLD.px.py
   })

# Add labels (for plotting)
assoc_labels <- as.vector(sapply(unique(rda_flat$z_val$Association), function (s) {
  s_split <- strsplit(s[1], "_")[[1]]
  s_label <- s_split
  label = paste(s_split[1], "\u2192", s_split[2])},
  simplify = TRUE))

summary_df <- data.frame(Association=assoc_labels,
                         Proportion_pos=proportion_pos,
                         Proportion_th=proportion_th,
                         kld=kld)

# Add associations, proportions and KLD to flat results (z-normalized/not normalized)
mean_associations <- ddply(rda_flat$val, "Association", function(x) mean(x$Values))

# Associations
rda_flat$val$Association <- factor(rda_flat$val$Association,
    levels=mean_associations$Association[order(mean_associations$V1)])

rda_flat$z_val$Association <- factor(rda_flat$z_val$Association,
    levels=mean_associations$Association[order(mean_associations$V1)])

# Proportions larger than zero
rda_flat$val$prop_pos <- sapply(rda_flat$val$Association,
                                function(x) proportion_pos[as.character(x)])

rda_flat$z_val$prop_pos <- rda_flat$val$prop_pos

# Proportions larger than diff_th
rda_flat$val$prop_th <-  sapply(rda_flat$val$Association,
                                function(x) proportion_th[as.character(x)])
rda_flat$z_val$prop_th <- rda_flat$val$prop_th

 # KLD
rda_flat$val$kld <-  sapply(rda_flat$val$Association,
                                function(x) kld[as.character(x)])
rda_flat$z_val$kld <- rda_flat$val$kld

return(list(summary = summary_df, rda_df=rda_flat))}
```

### S2 Data

#### S2.1 Lexical data

We load Levenshtein distances derived from the ASJP data (Wichmann et al. 2015):

```
lex_dist <- sort_dist_mat(as.dist(
  read.csv('data/lexicon/ASJP13PopDist.csv', header=T, row.names = 1)))

attr(lex_dist, 'Labels')[10] <- 'West Greenlandic'
```

#### S2.2 Genetics and Music

We read the distance matrices for genetics and music. Table S1 shows how these data are matched to unique identifiers in the language data. The UULID concatenates IDs from the ISO 639.3 standard ("i-"), from the AUTOTYP (Bickel et al. 2017) database ("a-"), and from the GLOTTOLOG (Hammarström, Forkel, and Haspelmath 2017) catalogue ("g-").

```
# Read the gentic data
genetics <- read.csv("data/genetics/SNPs14PopDist.csv",sep=",",
                     header=TRUE, row.names=1)

# Read the music data
music <- read.csv("data/music/Music14PopDist.csv",sep=",",
                  header=TRUE, row.names=1)
```

Table S1: Music and genome-wide SNP data, with sample sizes, references and match to languages.

| Population | SNPs | Music | SNP Sources | Language | UULID |
|---|---|---|---|---|---|
| Korean | 6 | 30 | Lazaridis et al. (2014) | Korean | [i-kor][a-141][g-kore1280] |
| Japanese (Mainland) | 45 | 30 | Abecasis et al. (2012) | Japanese | [i-jpn][a-118][g-nucl1643] |
| Ainu (Hokkaido) | 25 | 30 | Jinam et al. (2012) | Ainu | [i-ain][a-12][g-ainu1240] |
| Koryak | 24 | 30 | Rasmussen et al. (2010) | Koryak | [i-kpy][a-1808][g-kory1246] |
| Chukchi | 20 | 14 | Lazaridis et al. (2014) | Chukchi | [i-ckt][a-56][g-chuk1273] |
| Yakut | 21 | 8 | Lazaridis et al. (2014) | Yakut | [i-sah][a-2662][g-yaku1245] |
| Even | 14 | 17 | Lazaridis et al. (2014), Fedorova et al. (2013) | Even | [i-eve][a-738][g-even1260] |
| Yukagir | 5 | 12 | Lazaridis et al. (2014) | Yukagir (Tundra) | [i-yux][a-2797][g-sout2750] |
| Evenk | 15 | 28 | Rasmussen et al. (2010) | Evenki | [i-evn][a-527][g-even1259] |
| Buryat | 19 | 30 | Rasmussen et al. (2010) | Buriat | [i-bua][a-1095][g-buri1258] |

| Population | SNPs | Music | SNP Sources | Language | UULID |
|---|---|---|---|---|---|
| West Greenlandic (Inuit) | 5 | 8 | Rasmussen et al. (2010) | West Greenlandic | [i-kal][a-511][g-kala1399] |
| Selkup | 18 | 12 | Rasmussen et al. (2010), Lazaridis et al. (2014) | Selkup | [i-sel][a-2393][g-selk1253] |
| Nganasan | 13 | 15 | Rasmussen et al. (2010) | Nganasan | [i-nio][a-2172][g-ngan1291] |
| Nivkh | 15 | 19 | Fedorova et al. (2013), Matsumae et al., this study | Nivkh | [i-niv][a-433][g-gily1242] |

#### S2.3 Grammar and Phonology

Our data comprise numerical and categorical variables for grammar and phonology, distance matrices for genetics and music, and the geographical locations of peoples in Northern Asia and Greenland in the form of language polygons.

Data on grammar and phonology are aggregated from the following sources:

- AUTOTYP (Bickel et al. 2017)
- WALS (Dryer and Haspelmath 2013), enriched by recodings (Bickel and Zakharko 2018)
- ANU Phonotactics database (Donohue et al. 2013)
- PHOIBLE (Moran, McCloy, and Wright 2014)

The geographical polygon locations of the languages are taken from the Ethnologue (Simons and Fennig 2018); as there is no polygon for Ainu, we drew one ourselves.

The split into phonology vs. grammar is based on the broad definition in `categorization_of_variables.csv` which includes phonotactic and morphophonological data under phonology.[1] We extract a subset from the data comprising languages from fourteen different societies. Note that in AUTOTYP and WALS, Buriat is represented by ISO code `bua` (Buriat in general), while in PHOIBLE it is represented by ISO code `bxr` and in the ANU data by `bxm`. We map all of them below to `bua`.

```
# Define single languages and family-related languages
single_lgs <- c("Ainu", "Buryat", "Japanese", "Korean", "Nivkh", "West Greenlandic",
                "Yakut", "Yukagir")
uralic <- c("Selkup", "Nganasan")
chkkat <- c("Chukchi", "Koryak")
tungus <- c("Even", "Evenki")

# Read categorization of variables
var_categories <- read.csv("data/typology/categorization_of_variables.csv",
                           stringsAsFactors = F)

# Read the grammar, phonology and typlogy data (meta data)
typology.list <- readRDS("data/typology/typology.list.RDS")
phonological_vars <- unlist(as.character(sapply(typology.list, function(x) {
        var_categories[var_categories$Variable == x[1, "variable.ID"],
                       "Broad_Phon_Definition_Binary"] == "Phonology" }) ))
phonology_list <- typology.list[phonological_vars == "TRUE"]
grammar_vars <- unlist(as.character(sapply(typology.list, function(x) {
        var_categories[var_categories$Variable == x[1, "variable.ID"],
                       "Broad_Phon_Definition_Binary"] == "Grammar" })))
```

[1] Available at https://www.geo.uzh.ch/microsite/MusicGenesLanguages

```r
grammar_list <- typology.list[grammar_vars == "TRUE"]
typology_coverage_df <- readRDS("data/typology/typology.coverage.RDS")

# Define subset
siberia_sample <- c(
 "[i-ain][a-12][g-ainu1240]",    # Ainu
 "[i-bua][a-1095][g-buri1258]",  # Buriat
 "[i-bxm][a-][g-mong1330]",      # Buriat (Mongolia)
 "[i-bxr][a-][g-russ1264]",      # Buriat (Russia)
 "[i-ckt][a-56][g-chuk1273]",    # Chukchi
 "[i-eve][a-738][g-even1260]",   # Even
 "[i-evn][a-527][g-even1259]",   # Evenki
 "[i-kal][a-511][g-kala1399]",   # West Greenlandic
 "[i-jpn][a-118][g-nucl1643]",   # Japanese
 "[i-kor][a-141][g-kore1280]",   # Korean
 "[i-kpy][a-1808][g-kory1246]",  # Koryak
 "[i-nio][a-2172][g-ngan1291]",  # Nganasan
 "[i-niv][a-433][g-gily1242]",   # Nivkh
 "[i-sel][a-2393][g-selk1253]",  # Selkup
 "[i-sah][a-2662][g-yaku1245]",  # Yakut
 "[i-ykg][a-423][g-nort2745]")   # Yukagir (Tundra)

# Extract meta data for the above subset
siberia_metadata_all <- subset(typology_coverage_df, UULID %in% siberia_sample)
siberia_metadata <- subset(siberia_metadata_all, !isocode %in% c('bxm','bxr'))

counts <- xtabs(~autotyp.Stock, siberia_metadata, drop.unused.levels = T)
rownames(siberia_metadata) <- with(siberia_metadata,
                                   ifelse(autotyp.Stock %in% names(counts[counts>1]),
                                          paste(autotyp.Stock, autotyp.Language, sep="/"),
                                          paste(autotyp.Language)))

rownames(siberia_metadata) <- gsub('Yukagir/','', rownames(siberia_metadata))
```

We only use variables with one data point per language, and only variables with non-constant values (which otherwise can't deliver a distance signal). At the same time, we also remap `bxr` and `bxm` to `bua` (cf. above). We trim the linguistic data accordingly.

```r
# Trim the data
siberia_grammar_list <- trim_data(grammar_list,extra.coverage = 0.8)
siberia_phonology_list <- trim_data(phonology_list,extra.coverage = 0.8)
```

The following lists the phonological variables we captured. For full definitions and descriptions of the variables, see the source databases listed above.

```
## - WALS$FrontRndV.Presence
## - WALS$GlottalizedC.Presence
## - WALS$Laterals.Presence
## - WALS$Tone.Presence
## - WALS$Uvulars.Presence
## - WALS$VoicingC.Presence
## - WALS$11A Front Rounded Vowels
## - WALS$13A Tone
## - WALS$19A Presence of Uncommon Consonants
## - WALS$1A Consonant Inventories
```

```
## - WALS$2A Vowel Quality Inventories
## - WALS$3A Consonant-Vowel Ratio
## - WALS$4A Voicing in Plosives and Fricatives
## - WALS$6A Uvular Consonants
## - WALS$7A Glottalized Consonants
## - WALS$8A Lateral Consonants
## - PHOIBLE$syllabic_count
## - PHOIBLE$short_count
## - PHOIBLE$long_count
## - PHOIBLE$consonantal_count
## - PHOIBLE$sonorant_count
## - PHOIBLE$continuant_count
## - PHOIBLE$delayedRelease_count
## - PHOIBLE$approximant_count
## - PHOIBLE$trill_count
## - PHOIBLE$nasal_count
## - PHOIBLE$lateral_count
## - PHOIBLE$labial_count
## - PHOIBLE$round_count
## - PHOIBLE$labiodental_count
## - PHOIBLE$coronal_count
## - PHOIBLE$anterior_count
## - PHOIBLE$distributed_count
## - PHOIBLE$strident_count
## - PHOIBLE$dorsal_count
## - PHOIBLE$high_count
## - PHOIBLE$low_count
## - PHOIBLE$front_count
## - PHOIBLE$back_count
## - PHOIBLE$tense_count
## - PHOIBLE$retractedTongueRoot_count
## - PHOIBLE$periodicGlottalSource_count
## - PHOIBLE$spreadGlottis_count
## - PHOIBLE$constrictedGlottis_count
## - PHOIBLE$vowels_count
## - PHOIBLE$vowels.syllabic.consonants_count
## - PHOIBLE$long.vowels_count
## - PHOIBLE$glides_count
## - PHOIBLE$liquids_count
## - PHOIBLE$nasals_count
## - PHOIBLE$fricatives_count
## - PHOIBLE$affricates_count
## - PHOIBLE$stops_count
## - PHOIBLE$stops.affricates_count
## - PHOIBLE$liquids.glides_count
## - PHOIBLE$liquids.glides.nasals_count
## - PHOIBLE$creaky.breathy_count
## - PHOIBLE$aspirated.fricatives_count
## - PHOIBLE$voiceless.stops_count
## - PHOIBLE$velar.nasals_count
## - PHOIBLE$short_presence
## - PHOIBLE$long_presence
## - PHOIBLE$trill_presence
## - PHOIBLE$lateral_presence
```

```
## - PHOIBLE$labiodental_presence
## - PHOIBLE$distributed_presence
## - PHOIBLE$back_presence
## - PHOIBLE$retractedTongueRoot_presence
## - PHOIBLE$spreadGlottis_presence
## - PHOIBLE$constrictedGlottis_presence
## - PHOIBLE$long.vowels_presence
## - PHOIBLE$liquids_presence
## - PHOIBLE$affricates_presence
## - PHOIBLE$creaky.breathy_presence
## - PHOIBLE$aspirated.fricatives_presence
## - PHOIBLE$velar.nasals_presence
## - ANU$CVC language
## - ANU$CCVC language
## - ANU$Onset second C is preferentially glide
## - ANU$Onset second C = y
## - ANU$Onset second C = w
## - ANU$Palatals/Affricates (ts, c etc.) allowed as coda
## - ANU$Glottal stops in onsets?
## - ANU$Glottal stops in codas?
## - ANU$Velar nasals in onsets?
## - ANU$Velar nasals in codas?
```

And these the grammar variables:

```
## - autotyp$NP_per_language$AdjAttrAgr.Presence
## - autotyp$NP_per_language$AdjAttrConstr.Presence
## - autotyp$NP_per_language$AdjAttrGvt.Presence
## - autotyp$NP_per_language$AdjAttrMarking.overt.Presence
## - autotyp$NP_per_language$NPAgr.Presence
## - autotyp$NP_per_language$NPConstr.Presence
## - autotyp$NP_per_language$NPGvt.Presence
## - autotyp$NP_per_language$NPMarking.overt.Presence
## - WALS$AdjNounOrder
## - WALS$CaseMarking.Presence
## - WALS$CaseMarkingTypes.v1
## - WALS$InflAffixPositions
## - WALS$NegationType
## - WALS$PossAffixes.Presence
## - WALS$SOOrder
## - WALS$VInitialOrder
## - WALS$VerbMedialOrder
## - WALS$112A Negative Morphemes
## - WALS$12A Syllable Structure
## - WALS$26A Prefixing vs. Suffixing in Inflectional Morphology
## - WALS$51A Position of Case Affixes
## - WALS$57A Position of Pronominal Possessive Affixes
## - WALS$69A Position of Tense-Aspect Affixes
## - WALS$73A The Optative
## - WALS$87A Order of Adjective and Noun
## - WALS$88A Order of Demonstrative and Noun
```

For each society we compute the coverage, i.e. the percentage of available variables per society.

```
# Compute the coverage for each variable
grammar_coverage <- compute_coverage(siberia_grammar_list) %>%
```

Table S2: Language data coverage for all fourteen societies.

| Language | Grammar (%) | Phonology (%) |
|---|---|---|
| Chukchi | 100 | 100 |
| Evenki | 100 | 100 |
| Japanese | 100 | 100 |
| West Greenlandic | 100 | 100 |
| Koryak | 100 | 100 |
| Nivkh | 100 | 100 |
| Yakut | 100 | 100 |
| Ainu | 96 | 100 |
| Buryat | 96 | 81 |
| Nganasan | 96 | 100 |
| Even | 92 | 100 |
| Korean | 92 | 100 |
| Selkup | 88 | 98 |
| Yukagir | 88 | 100 |

```
  dplyr::select(Language, Coverage)
phonology_coverage <- compute_coverage(siberia_phonology_list) %>%
  dplyr::select(Language, Coverage)
```

We simplify and standardize the language names and visualize the coverage in a table. Finally, we flatten the nested linguistic data and convert them to data frames.

```
# Flatten the data
grammar <- flatten(siberia_grammar_list)
phonology <- flatten(siberia_phonology_list)
```

#### S2.4 Geographic locations

We import the language polygons and 15,000 samples of point locations taken randomly from these. The random point samples were generated in PostGIS with the function `ST_GeneratePoints` For further details see the SQL code `generate_random_point_samples.sql` at https://www.geo.uzh.ch/microsite/MusicGenesLanguages/.

```
# Fetch the language polygons
geo_polygons <- readRDS("data/geo/geo_polygons.RDS")

# Fetch random spatial points in the polygons
geo_random_points <- readRDS("data/geo/geo_random_points.RDS")
```

Since the data are gathered from different sources, the names used for the fourteen societies differ. We standardize all names.

```
# We update all non-matching names using the names in geo_random_points as a template
# Genetics
colnames(genetics)[colnames(genetics) == 'westGreenland'] <- 'West Greenlandic'
rownames(genetics)[rownames(genetics) == 'westGreenland'] <- 'West Greenlandic'
colnames(genetics)[colnames(genetics) == 'Evenk'] <- 'Evenki'
rownames(genetics)[rownames(genetics) == 'Evenk'] <- 'Evenki'
```

```r
# Music
colnames(music)[colnames(music) == 'WestGreenland'] <- 'West Greenlandic'
rownames(music)[rownames(music) == 'WestGreenland'] <- 'West Greenlandic'
colnames(music)[colnames(music) == 'Nganasa'] <- 'Nganasan'
rownames(music)[rownames(music) == 'Nganasa'] <- 'Nganasan'
colnames(music)[colnames(music) == 'Evenk'] <- 'Evenki'
rownames(music)[rownames(music) == 'Evenk'] <- 'Evenki'

# Grammar
rownames(grammar)[rownames(grammar) == 'Chukchi-Kamchatkan/Chukchi'] <-'Chukchi'
rownames(grammar)[rownames(grammar) == 'Tungusic/Evenki'] <- 'Evenki'
rownames(grammar)[rownames(grammar) == 'Greenlandic Eskimo (West)'] <-'West Greenlandic'
rownames(grammar)[rownames(grammar) == 'Uralic/Selkup'] <- 'Selkup'
rownames(grammar)[rownames(grammar) == 'Yukagir (Tundra)'] <- 'Yukagir'
rownames(grammar)[rownames(grammar) == 'Tungusic/Even'] <- 'Even'
rownames(grammar)[rownames(grammar) == 'Buriat'] <- 'Buryat'
rownames(grammar)[rownames(grammar) == 'Uralic/Nganasan'] <- 'Nganasan'
rownames(grammar)[rownames(grammar) == 'Chukchi-Kamchatkan/Koryak'] <- 'Koryak'

# Phonology
rownames(phonology)[rownames(phonology) == 'Chukchi-Kamchatkan/Chukchi'] <-'Chukchi'
rownames(phonology)[rownames(phonology) == 'Tungusic/Evenki'] <- 'Evenki'
rownames(phonology)[rownames(phonology) == 'Greenlandic Eskimo (West)'] <-'West Greenlandic'
rownames(phonology)[rownames(phonology) == 'Uralic/Selkup'] <- 'Selkup'
rownames(phonology)[rownames(phonology) == 'Yukagir (Tundra)'] <- 'Yukagir'
rownames(phonology)[rownames(phonology) == 'Tungusic/Even'] <- 'Even'
rownames(phonology)[rownames(phonology) == 'Buriat'] <- 'Buryat'
rownames(phonology)[rownames(phonology) == 'Uralic/Nganasan'] <- 'Nganasan'
rownames(phonology)[rownames(phonology) == 'Chukchi-Kamchatkan/Koryak'] <- 'Koryak'
```

### S3 Dimensionality reduction

#### S3.1 Factorial analysis of mixed data (FAMD) of Grammar and Phonology

In view of the fact that the grammar and phonology data are partly numerical and partly categorical, we use a balanced mix of PCA and MCA (Lê, Josse, and Husson 2008) to bring out the main patterns in the data. Empty values are imputed using the methods developed by Josse and Husson (2016).

```r
# Impute empty values
grammar_imputed <- imputeFAMD(grammar[,-1])
phonology_imputed <- imputeFAMD(phonology[,-1])

# Perform FAMD
grammar_famd <- FAMD(grammar[,-1],
                     tab.disj=grammar_imputed$tab.disj,
                     ncp=10,
                     graph=F)

phonology_famd <- FAMD(phonology[,-1],
                       ncp=10,
                       tab.disj=phonology_imputed$tab.disj,
                       graph=F)
```

We rescale the dimensions obtained through FAMD in relation to the explained variance:

```
for(i in 1:ncol(phonology_famd$ind$coord)) {
  phonology_famd$ind$coord[,i] <-
    scale(phonology_famd$ind$coord[,i])*
    phonology_famd$eig[i,"percentage of variance"]}

for(i in 1:ncol(grammar_famd$ind$coord)) {
  grammar_famd$ind$coord[,i]<-
    scale(grammar_famd$ind$coord[,i])*
    grammar_famd$eig[i,"percentage of variance"]}
```

#### S3.2 Principal Coordinates Analysis (PCoA) of Music and Genes

We perform a principal coordinate analysis (PCoA) on the distance matrices for genetics and music. Similar to a PCA, a PCoA produces a set of orthogonal axes whose importance is measured by eigenvalues (Dray, Legendre, and Peres-Neto 2006). However, in contrast to the PCA, non-Euclidean distance matrices can be used. We correct for negative eigenvalues using the Cailliez procedure.

```
# Convert the matrices into dist objects
genetics_dist <- as.dist(genetics, diag = FALSE, upper = FALSE)
music_dist <- as.dist(music, diag = FALSE, upper = FALSE)

# Perform PCoA
genetics_pcoa <- pcoa(genetics_dist, correction = "cailliez")
music_pcoa <- pcoa(music_dist, correction = "cailliez")
```

We rescale the PCoA components in relation to the explained variance.

```
for(i in 1:ncol(genetics_pcoa$vectors)) {
genetics_pcoa$vectors[,i] <- scale(genetics_pcoa$vectors[,i])*
  genetics_pcoa$values$Rel_corr_eig[i]}

for(i in 1:ncol(music_pcoa$vectors)) {
music_pcoa$vectors[,i] <- scale(music_pcoa$vectors[,i])*
  music_pcoa$values$Rel_corr_eig[i]}
```

#### S3.3 Distance-based Moran's Eigenvector Map Analysis (dbMEM) of the spatial locations

We take 1,000 random point locations from the language polygons (as represented in Figure S1) and compute the spherical distance between them. Then we perform a distance-based Moran's eigenvector map analysis (`dbMEM`) where we decompose the spatial structure of each of the resulting 1,000 distance matrices (Borcard and Legendre 2002). Similar to a PCoA, dbMEM reveals the principal coordinates of the spatial locations from which the distance matrix was generated. However, in contrast to PCoA, dbMEM is primarily concerned with the interaction between spatial neighbors. Thus, only distances below a certain threshold feed directly into constructing the principal coordinates, whereas distances above the threshold are "truncated" (i.e. they are set to four times the threshold value). In the dbMEM, we use the length of the longest edge in the minimum spanning tree as a truncation threshold. Moreover, we only return those eigenfunctions that correspond to positive autocorrelation.

```
# Some random points are not complete. We remove them.
incomplete_samples <- as.data.frame(geo_random_points) %>%
```

```r
  dplyr::group_by(sample_id) %>%
  dplyr::summarise (n=n()) %>%
  dplyr::filter(n!=14)

incomplete_samples <- as.vector(incomplete_samples$sample_id)
geo_random_points <- geo_random_points[!geo_random_points$sample_id
                                       %in% incomplete_samples, ]

# We take 1000 samples from the random points and compute dbMEMs for each sample
n_geo_samples = 1000
choose_p <- sample(unique(geo_random_points$sample_id), n_geo_samples, replace=F)
geo_mem <- random_points_to_dbmem(geo_random_points[geo_random_points$sample_id
                                                    %in% choose_p, ])
```

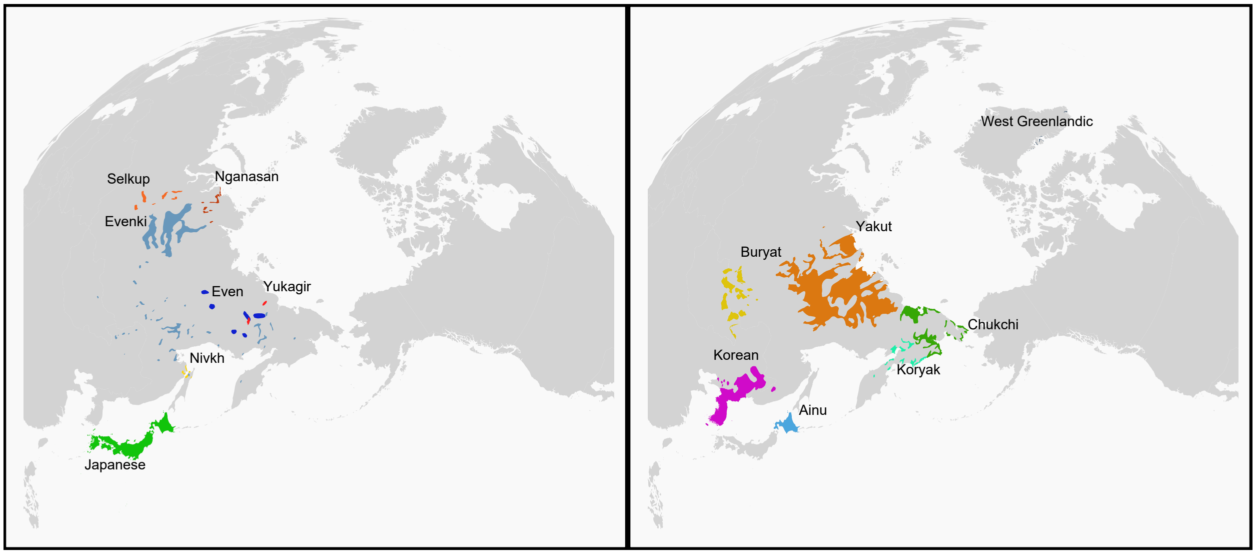

Figure S1: Geographical locations of the fourteen languages. Language polygons are plotted on two panels because they partly overlap in spatial distributions. Similarly-colored pairs of languages belong to the same family: Even and Evenki belong to the Tungusic family, Selkup and Nganasan to the Uralic family, and Koryak and Chukchi to the Chukotko-Kamchatkan family.

#### S3.4 Visualizing the explained variance

We visualize the results of the PCA and PCoA in a scree plot. The figures below show the fraction of total variance in the data as explained by each PC/PCo in decreasing order. We extract the eigenvalues from the PCos/PCs and visualize the explained variance.

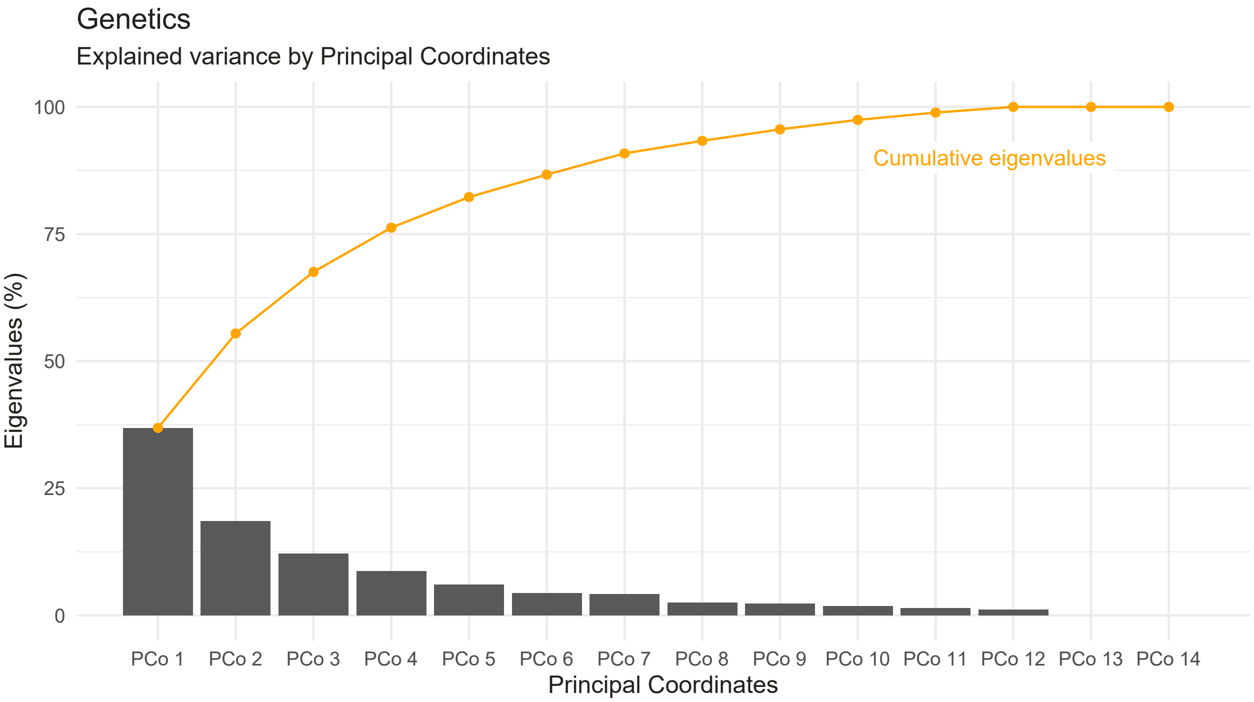

Figure S2: Scree plot of explained variance for Genetics

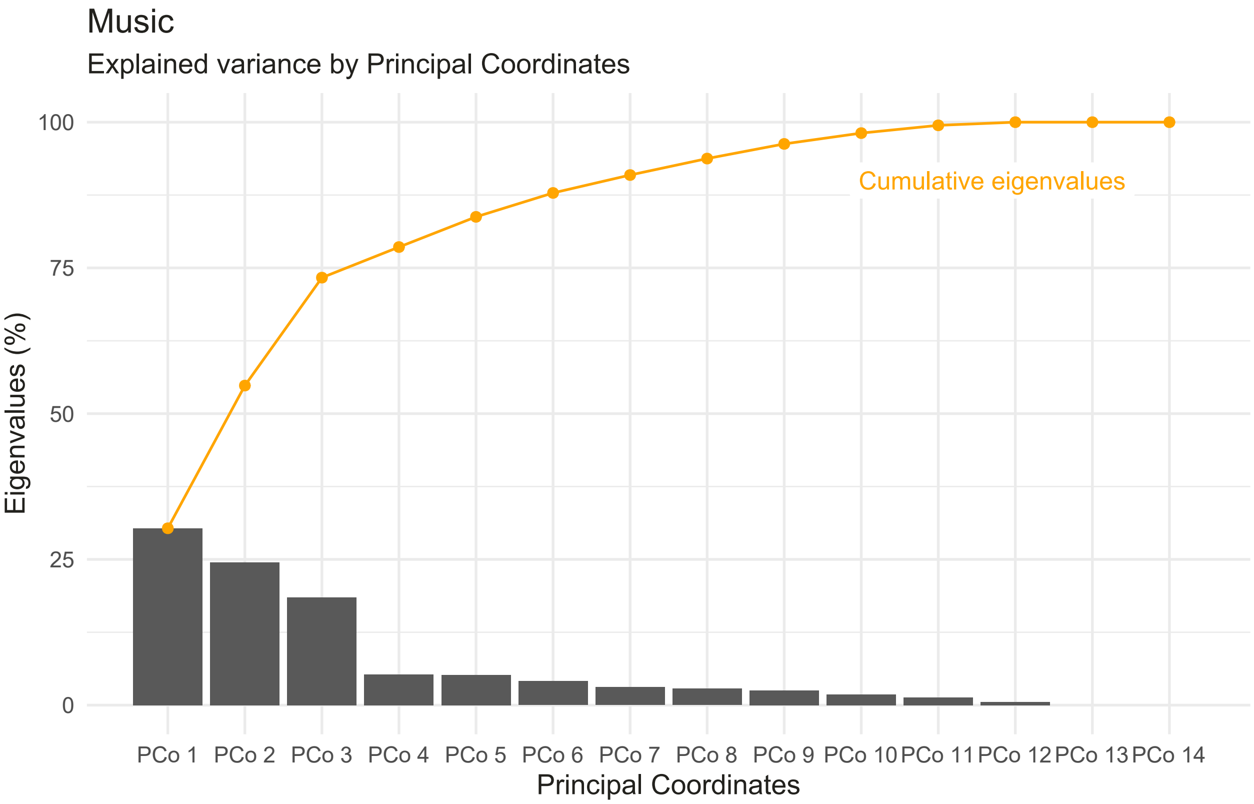

Figure S3: Scree plot of explained variance for Music

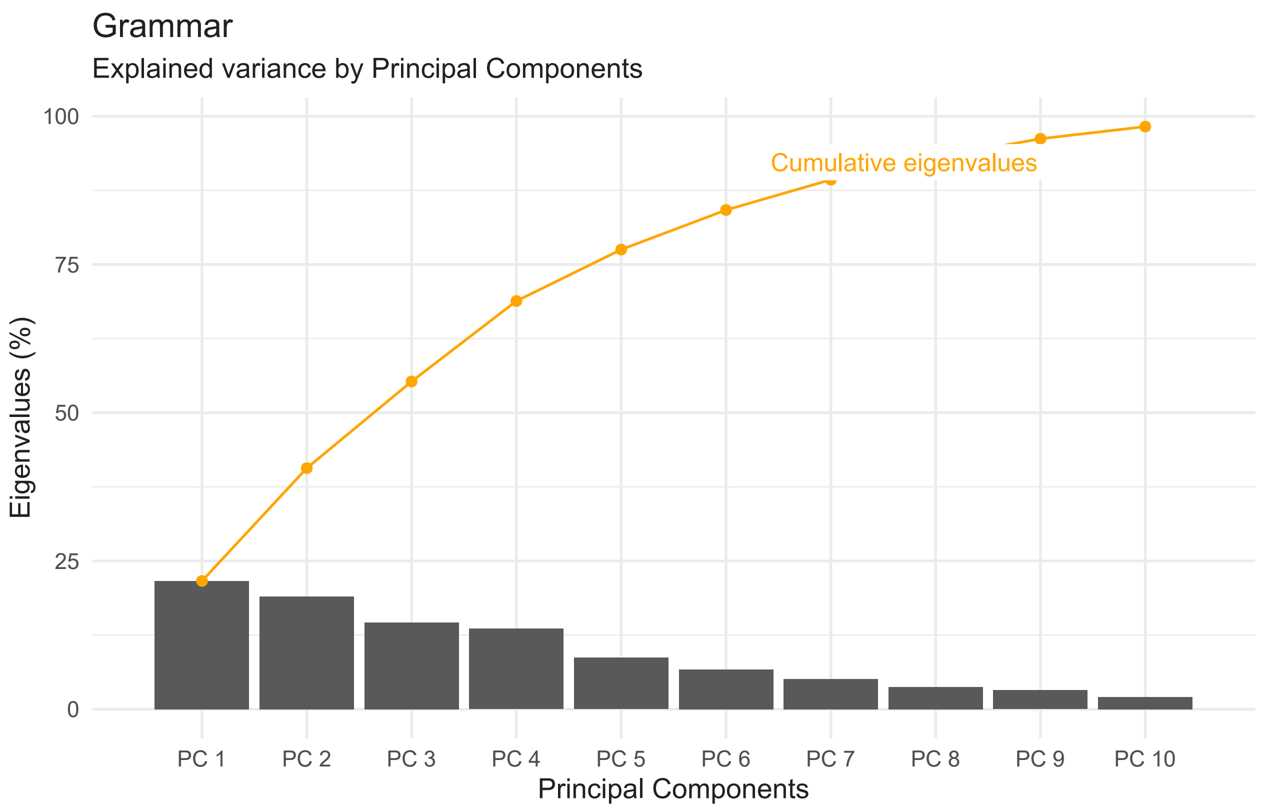

Figure S4: Scree plot of explained variance for Grammar

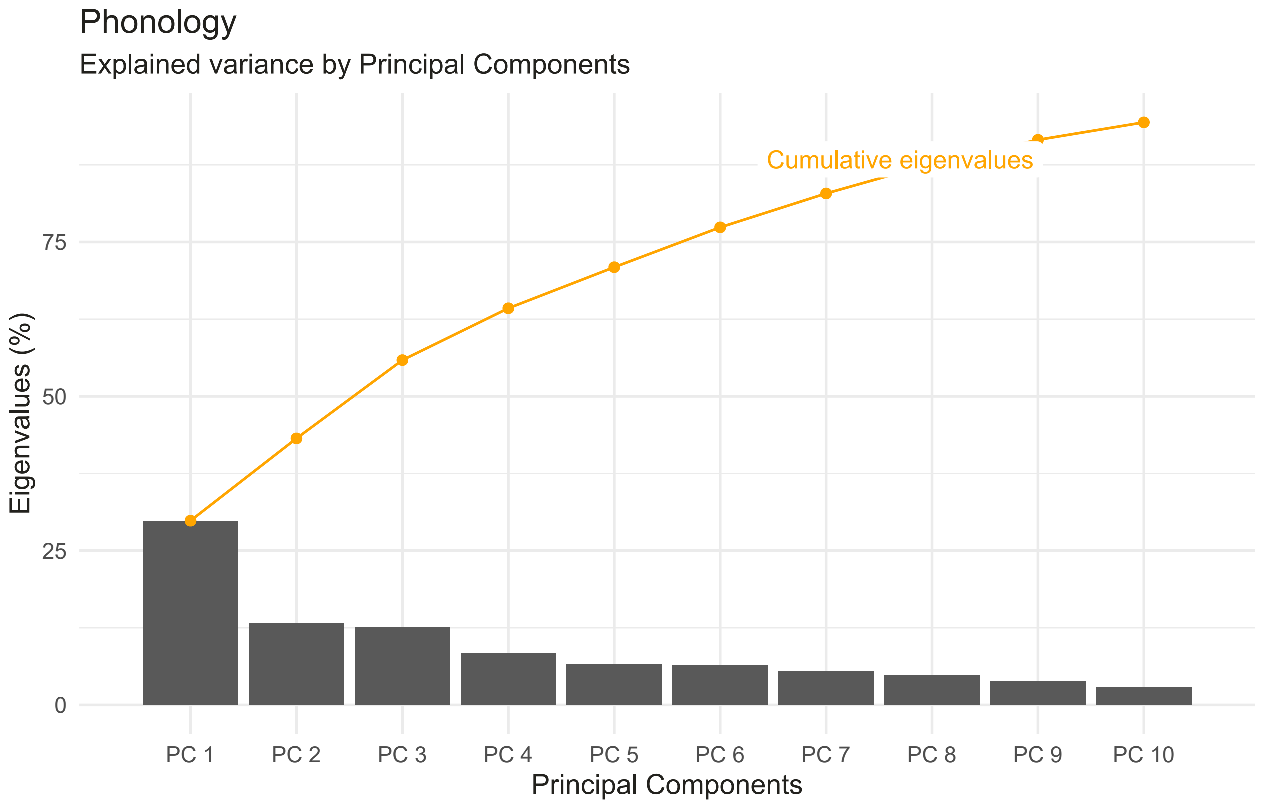

Figure S5: Scree plot of explained variance for Phonology

#### S3.5 Heatmaps of PCs and PCos

We extract the principal components/coordinates for all factors: genetics, grammar, music and phonology. We normalize the PCs/PCos to a range from 0 to 1 and then plot a heatmap for each factor.

```
# Extract the PCs and the PCos from the PCA and PCoA results
genetics_pco <- genetics_pcoa$vectors
music_pco <- music_pcoa$vectors
grammar_pc <- grammar_famd$ind$coord
phonology_pc <- phonology_famd$ind$coord

# Change the column names of all PCs and PCoAs
colnames(genetics_pco)<- paste("genetics_pco_", seq(1,ncol(genetics_pco)), sep="")
colnames(music_pco) <- paste("music_pco_", seq(1,ncol(music_pco)), sep="")
colnames(grammar_pc) <- paste("grammar_pc_", seq(1,ncol(grammar_pc)), sep="")
colnames(phonology_pc) <- paste("phonology_pc_", seq(1,ncol(phonology_pc)), sep="")
```

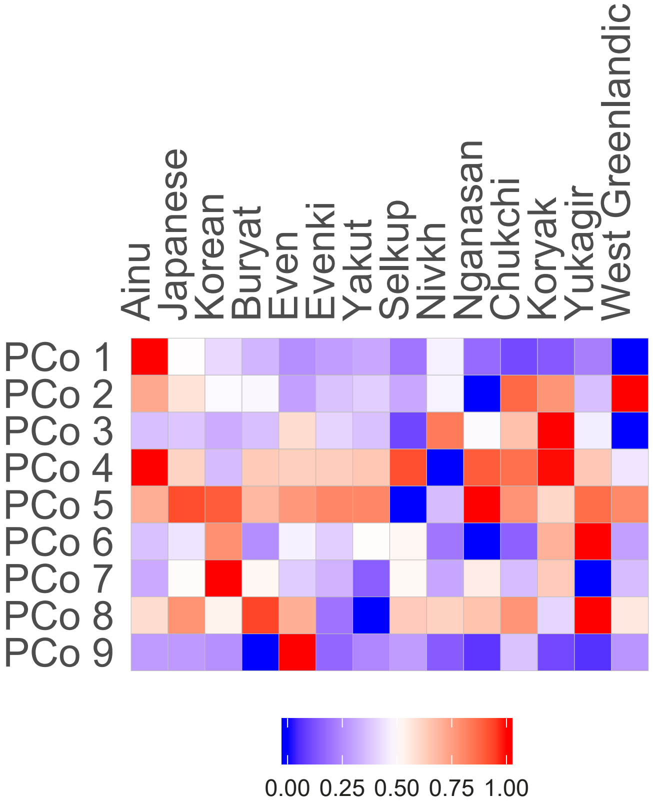

Figure S6: Heat plot of the first nine PCos (normalized) of Genetics

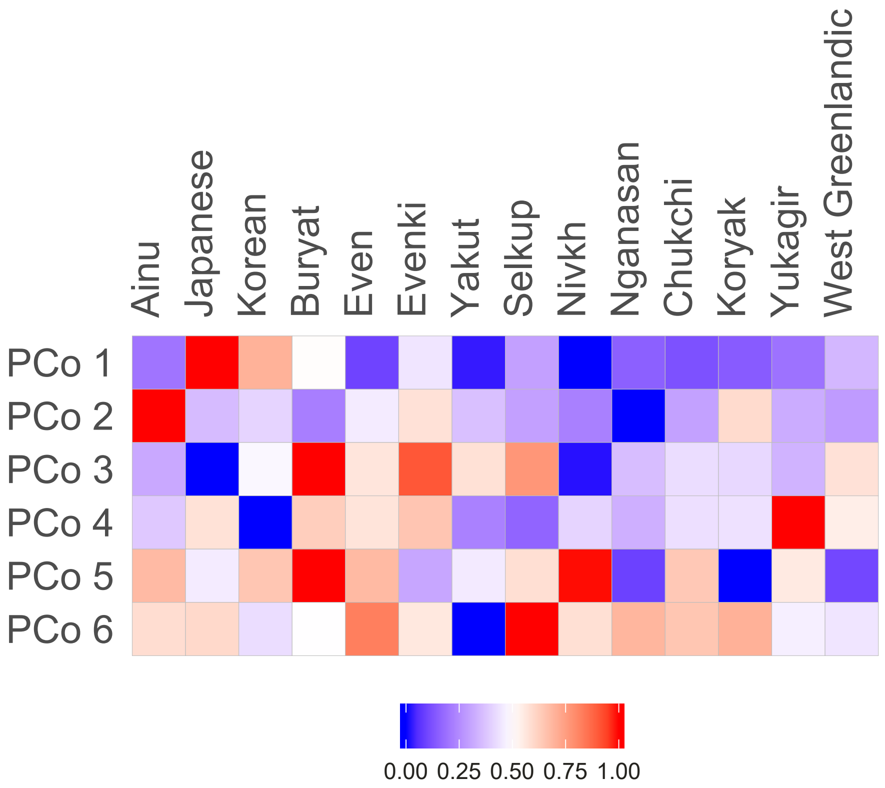

Figure S7: Heat plot of the first six PCos (normalized) of Music

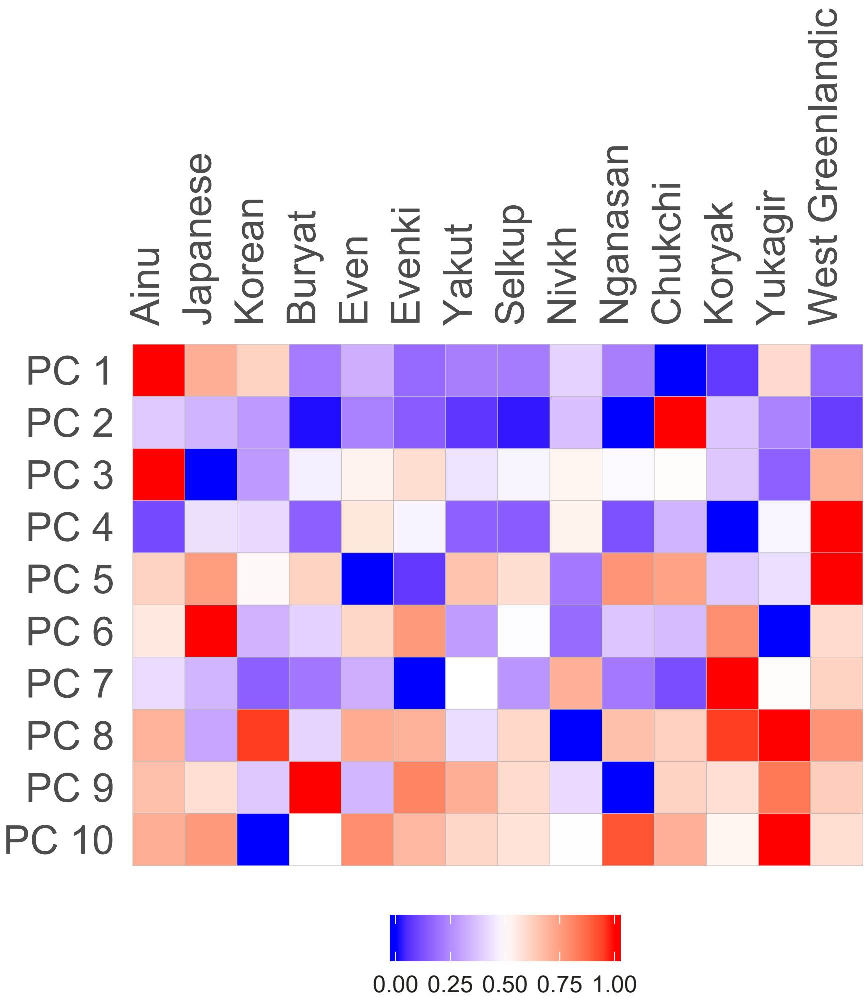

Figure S8: Heat plot of the first ten PCs (normalized) of Grammar

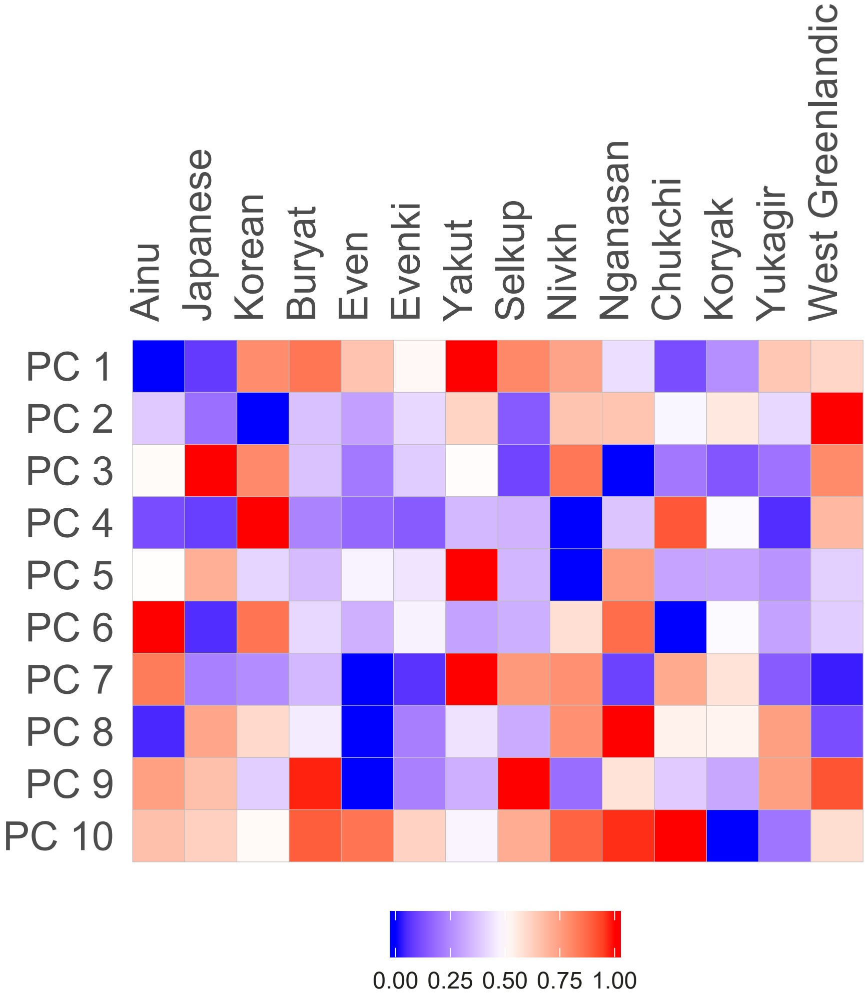

Figure S9: Heat plot of the first ten PCs (normalized) of Phonology

### S4 Distance visualization (NeighborNets)

Lexical distances are visualized directly. For nonlexical data, we compute Euclidean distances from the dimensionality-reduced data. All NeighborNets were created with SplitsTree (Huson and Bryant 2006).

Table S3: Lexical distances

| | Ainu | Buryat | Chukchi | Even | Evenki | Japanese | Korean | Koryak | Nivkh | WstGrnln | WstGrnln | Yakut |
|---|---|---|---|---|---|---|---|---|---|---|---|---|
| Buryat | 9788 | | | | | | | | | | | |
| Chukchi | 10093 | 9766 | | | | | | | | | | |
| Even | 9754 | 9258 | 9509 | | | | | | | | | |
| Evenki | 9880 | 9157 | 9757 | 6009 | | | | | | | | |
| Japanese | 9670 | 10178 | 9993 | 10036 | 10023 | | | | | | | |
| Korean | 9963 | 9979 | 9788 | 9881 | 9490 | 10054 | | | | | | |
| Koryak | 10093 | 9632 | 5809 | 9527 | 9852 | 10185 | 9806 | | | | | |
| Nivkh | 9919 | 9783 | 9859 | 9717 | 9508 | 9923 | 9644 | 9660 | | | | |
| WstGrnln | 9849 | 10229 | 10030 | 9737 | 9766 | 9879 | 9891 | 9857 | 10126 | | | |
| WstGrnln | 10145 | 10217 | 10012 | 9765 | 9710 | 10077 | 9975 | 10006 | 9745 | 9909 | | |
| Yakut | 10091 | 9527 | 10129 | 10009 | 10015 | 10437 | 10113 | 10132 | 9561 | 9868 | 10277 | |
| Yukagir | 10236 | 10108 | 9846 | 10005 | 10027 | 9672 | 9875 | 9708 | 10066 | 9759 | 10087 | 9561 |

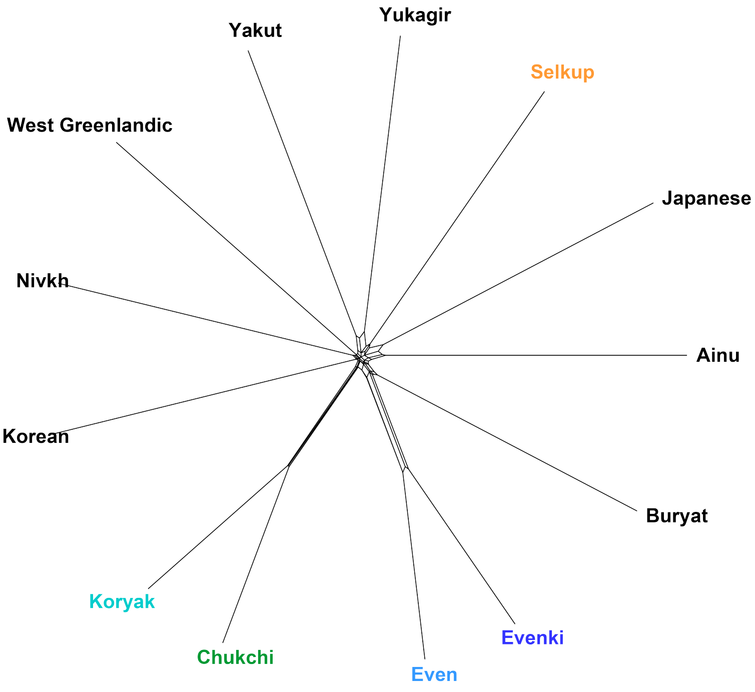

Figure S10: Lexical (ASJP) distances (Nganasan missing)

Table S4: Genetic distances

| | Ainu | Buryat | Chukchi | Even | Evenki | Japanese | Korean | Koryak | Nganasan | Nivkh | Selkup | WstGrnln | Yakut |
|---|---|---|---|---|---|---|---|---|---|---|---|---|---|
| Buryat | 1.027 | | | | | | | | | | | | |
| Chukchi | 1.374 | 0.482 | | | | | | | | | | | |
| Even | 1.189 | 0.240 | 0.482 | | | | | | | | | | |
| Evenki | 1.121 | 0.145 | 0.478 | 0.146 | | | | | | | | | |
| Japanese | 0.777 | 0.266 | 0.663 | 0.446 | 0.366 | | | | | | | | |
| Korean | 0.940 | 0.203 | 0.612 | 0.350 | 0.273 | 0.201 | | | | | | | |
| Koryak | 1.348 | 0.496 | 0.221 | 0.454 | 0.479 | 0.660 | 0.608 | | | | | | |
| Nganasan | 1.364 | 0.446 | 0.638 | 0.293 | 0.341 | 0.658 | 0.560 | 0.617 | | | | | |
| Nivkh | 0.918 | 0.369 | 0.692 | 0.439 | 0.418 | 0.335 | 0.341 | 0.648 | 0.668 | | | | |
| Selkup | 1.274 | 0.337 | 0.533 | 0.325 | 0.297 | 0.568 | 0.471 | 0.557 | 0.381 | 0.630 | | | |
| WstGrnln | 1.558 | 0.665 | 0.384 | 0.697 | 0.660 | 0.849 | 0.763 | 0.573 | 0.807 | 0.915 | 0.627 | | |
| Yakut | 1.083 | 0.144 | 0.496 | 0.187 | 0.062 | 0.335 | 0.258 | 0.500 | 0.387 | 0.410 | 0.315 | 0.669 | |
| Yukagir | 1.230 | 0.264 | 0.447 | 0.176 | 0.176 | 0.479 | 0.382 | 0.442 | 0.342 | 0.510 | 0.308 | 0.622 | 0.197 |

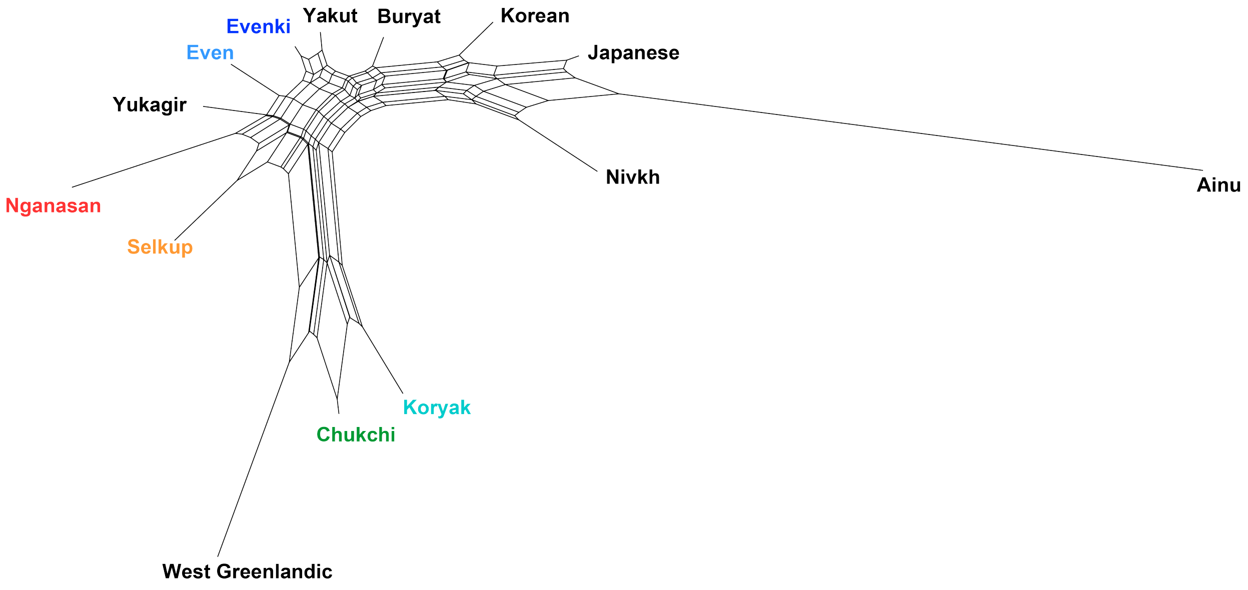

Figure S11: Genetic distances

Table S5: Music distances

| | Ainu | Buryat | Chukchi | Even | Evenki | Japanese | Korean | Koryak | Nganasan | Nivkh | Selkup | WstGrnln | Yakut |
|---|---|---|---|---|---|---|---|---|---|---|---|---|---|
| Buryat | 0.994 | | | | | | | | | | | | |
| Chukchi | 0.752 | 0.565 | | | | | | | | | | | |
| Even | 0.609 | 0.586 | 0.200 | | | | | | | | | | |
| Evenki | 0.650 | 0.404 | 0.556 | 0.460 | | | | | | | | | |
| Japanese | 1.123 | 0.852 | 0.990 | 1.051 | 0.871 | | | | | | | | |
| Korean | 0.840 | 0.464 | 0.644 | 0.674 | 0.455 | 0.471 | | | | | | | |
| Koryak | 0.454 | 0.692 | 0.333 | 0.218 | 0.458 | 0.999 | 0.652 | | | | | | |
| Nganasan | 1.064 | 0.630 | 0.333 | 0.520 | 0.776 | 1.019 | 0.747 | 0.630 | | | | | |
| Nivkh | 0.871 | 0.839 | 0.314 | 0.452 | 0.846 | 1.095 | 0.846 | 0.523 | 0.398 | | | | |
| Selkup | 0.811 | 0.321 | 0.300 | 0.319 | 0.377 | 0.918 | 0.502 | 0.439 | 0.449 | 0.597 | | | |
| WstGrnln | 0.799 | 0.362 | 0.279 | 0.349 | 0.393 | 0.805 | 0.436 | 0.410 | 0.396 | 0.548 | 0.204 | | |
| Yakut | 0.728 | 0.627 | 0.208 | 0.213 | 0.567 | 1.129 | 0.744 | 0.333 | 0.458 | 0.420 | 0.375 | 0.380 | |
| Yukagir | 0.731 | 0.564 | 0.162 | 0.260 | 0.537 | 0.905 | 0.604 | 0.338 | 0.382 | 0.346 | 0.363 | 0.261 | 0.31 |

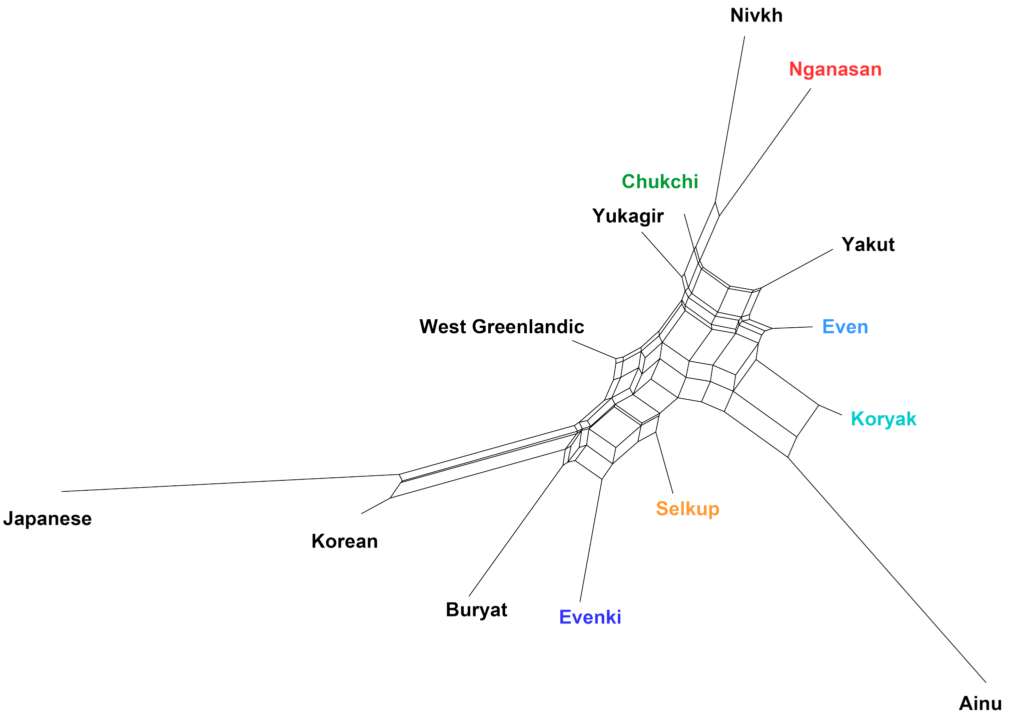

Figure S12: Music distances

Table S6: Grammar distances

| | Ainu | Buryat | Chukchi | Even | Evenki | Japanese | Korean | Koryak | Nganasan | Nivkh | Selkup | WstGrnln | Yakut |
|---|---|---|---|---|---|---|---|---|---|---|---|---|---|
| Buryat | 75.220 | | | | | | | | | | | | |
| Chukchi | 95.900 | 75.958 | | | | | | | | | | | |
| Even | 68.160 | 35.847 | 68.124 | | | | | | | | | | |
| Evenki | 76.030 | 29.244 | 69.000 | 17.370 | | | | | | | | | |
| Japanese | 69.450 | 58.393 | 81.867 | 52.571 | 61.778 | | | | | | | | |
| Korean | 57.587 | 43.093 | 73.714 | 34.628 | 45.264 | 29.168 | | | | | | | |
| Koryak | 82.598 | 37.421 | 55.411 | 42.185 | 39.999 | 62.134 | 52.438 | | | | | | |
| Nganasan | 74.860 | 14.944 | 77.064 | 38.646 | 34.601 | 59.738 | 44.042 | 39.344 | | | | | |
| Nivkh | 62.072 | 40.996 | 61.581 | 21.449 | 33.740 | 49.455 | 31.379 | 44.318 | 43.910 | | | | |
| Selkup | 73.925 | 6.698 | 75.155 | 33.505 | 27.743 | 57.602 | 41.948 | 34.862 | 10.786 | 40.090 | | | |
| WstGrnln | 84.779 | 49.393 | 79.275 | 43.243 | 42.201 | 71.120 | 57.977 | 63.575 | 49.978 | 47.573 | 48.775 | | |
| Yakut | 73.734 | 8.920 | 72.031 | 34.521 | 31.277 | 55.966 | 40.337 | 33.107 | 13.791 | 36.660 | 8.826 | 48.733 | |
| Yukagir | 66.960 | 45.379 | 78.453 | 38.534 | 49.454 | 33.230 | 17.989 | 55.337 | 47.646 | 34.173 | 45.269 | 60.144 | 41.82 |

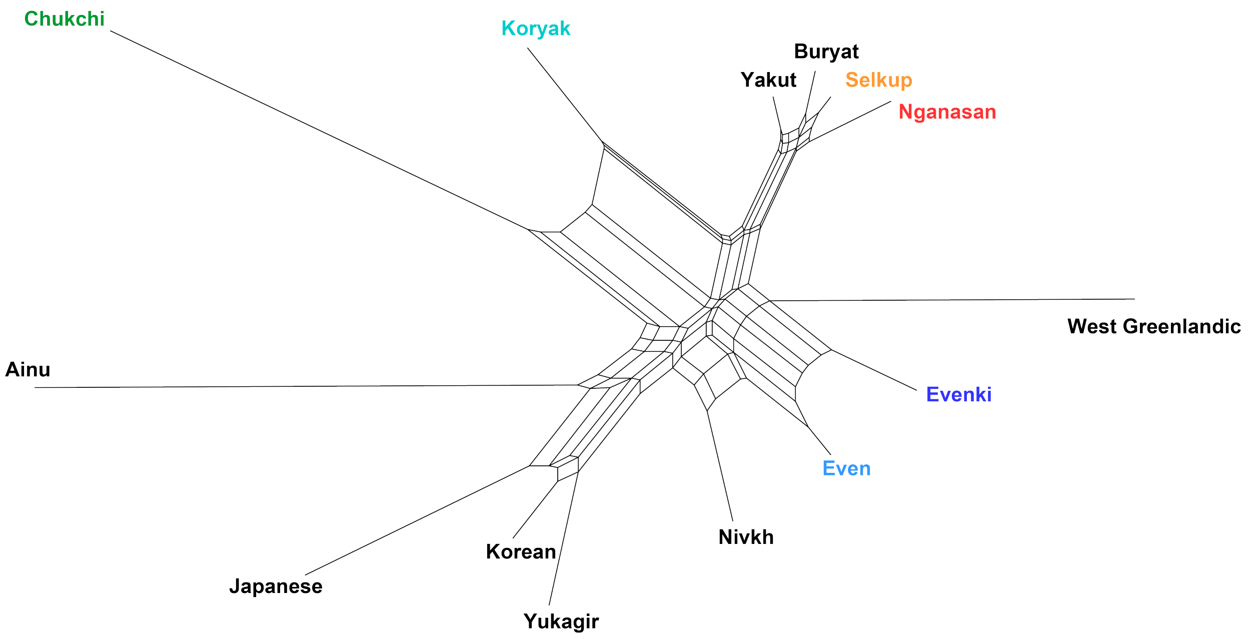

Figure S13: Grammar distances (scaled)

Table S7: Phonology distances

| | Ainu | Buryat | Chukchi | Even | Evenki | Japanese | Korean | Koryak | Nganasan | Nivkh | Selkup | WstGrnln | Yakut |
|---|---|---|---|---|---|---|---|---|---|---|---|---|---|
| Buryat | 82.345 | | | | | | | | | | | | |
| Chukchi | 36.977 | 73.211 | | | | | | | | | | | |
| Even | 67.390 | 24.833 | 58.435 | | | | | | | | | | |
| Evenki | 52.798 | 33.258 | 47.047 | 17.398 | | | | | | | | | |
| Japanese | 35.192 | 78.968 | 44.218 | 65.949 | 52.242 | | | | | | | | |
| Korean | 83.512 | 35.771 | 75.984 | 41.625 | 45.517 | 75.865 | | | | | | | |
| Koryak | 36.199 | 59.463 | 23.452 | 44.754 | 31.865 | 47.943 | 67.093 | | | | | | |
| Nganasan | 52.651 | 48.477 | 43.839 | 37.855 | 29.075 | 61.058 | 61.919 | 27.810 | | | | | |
| Nivkh | 75.912 | 31.724 | 71.033 | 39.891 | 36.044 | 72.145 | 46.918 | 57.168 | 51.927 | | | | |
| Selkup | 81.036 | 18.197 | 70.397 | 25.116 | 36.494 | 79.689 | 37.838 | 58.044 | 50.368 | 44.731 | | | |
| WstGrnln | 71.476 | 46.143 | 61.818 | 47.718 | 39.616 | 70.313 | 58.211 | 51.039 | 46.261 | 36.069 | 58.322 | | |
| Yakut | 98.633 | 30.588 | 89.043 | 45.252 | 52.336 | 94.340 | 49.350 | 75.574 | 62.311 | 41.750 | 40.555 | 51.101 | |
| Yukagir | 66.971 | 23.014 | 57.305 | 18.950 | 19.547 | 65.372 | 45.701 | 41.085 | 33.642 | 34.369 | 26.143 | 44.265 | 45.787 |

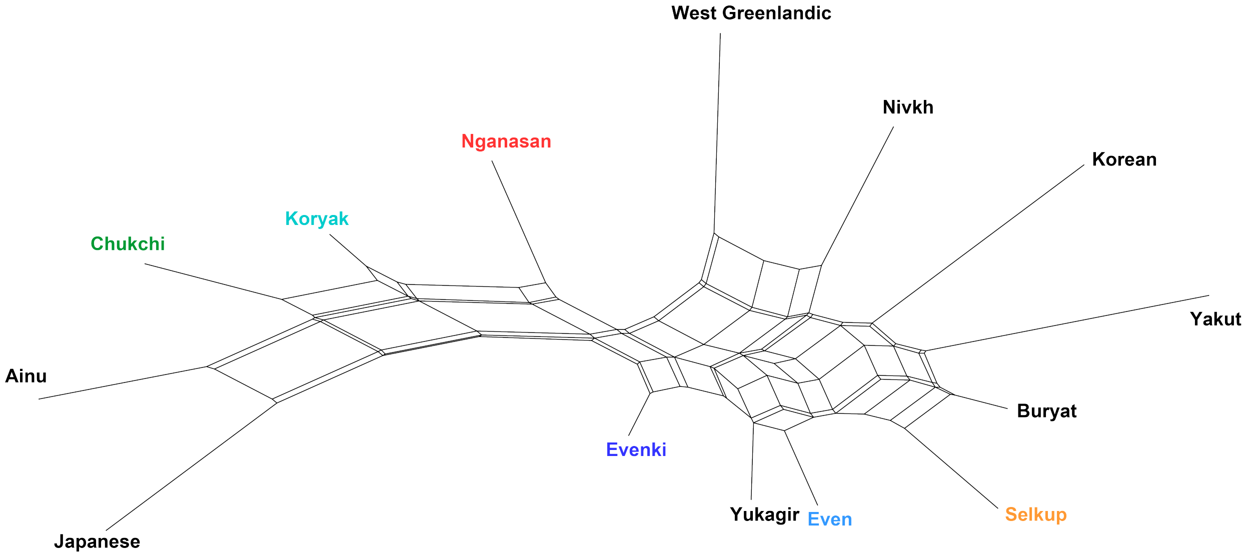

Figure S14: Phonology distances (scaled)

### S5 Redundancy Analysis (RDA)

Redundancy Analysis (RDA) extracts the variation in a set of explanatory variables that can be explained by a set of response variables (Legendre and Legendre 2012). In our case the response and explanatory variables comprise the principle components (coordinates) of a factor, e.g. the explanatory variable comprises the PCs of grammar and the response those of phonology. RDA carries out a multiresponse multiple linear regression, that is a regression of multiple response variables on multiple explanatory variables (Van Den Wollenberg 1977). We perform RDA for three different setups and report the associations found in the data:

- **Setup 0: RDA:** In this setup we do not control for potential confounding factors.
- **Setup 1: Partial RDA controlling for spatial autocorrelation:** In this setup we account for the influence of space. We take random point locations from the language polygons in turn and partial out their effect on the response before running the RDA. Thus, we explore to what degree the associations are affected by different spatial neighborhood scenarios.
- **Setup 2: Partial RDA controlling for spatial autocorrelation and genealogy**: In addition to spatial neighborhood we also control for genalogy. We perform a partial RDA for random samples of non-related languages only and explore to what degree the associations are driven by phylogenetic inheritance.

For each setup we report the observed explained variance (adjusted $R^2$) and compare it to the explained variance of random permutations. For associations with non-random, consistently high explained variance, we perform three types of sensitivity analysis to explore how changes in the model affect the results:

- **Sensitivity 1: Influence of Principal components/coordinates** Given the small number of observations (fourteen societies) using all principal components/coordinates would yield an over-determined model. Hence, we only retain those PCs/PCos per factor which account for at least $k$% of the explained variance. We run the partial RDA with different values for $k$ and compare the results.
- **Sensitivity 2: Influence of sampling** Some societies might have a larger contribution to the associations than others. We exclude each society in turn and run a partial RDA to explore the influence of each society on the associations.
- **Senisitivity 3: Spatial distribution of $R^2$** We explore the spatial locations of samples for which the adjusted $R^2$ is particularly low/high. If locations with low and high (adjusted) $R^2$ differ in space, this might indicate that a particular neighborhood scenario (low $R^2$) explains the variance in the response, while another one does not (high $R^2$).

#### S5.1 RDA with different setups

Before the analysis, we match the order of societies for each factor and combine all factors in a list. We find all combinations of factors by expanding the factor names to a grid. We iterate over this grid, such that each of factor once functions as a predictor and once as a response in the RDA.

```
# Retain PCs/PCos
factors <- pcos_to_factors(var_th=0.15,
                           genetics=genetics_pco,
                           genetics_ev=genetics_pcoa$values$Rel_corr_eig,
                           grammar=grammar_pc, grammar_ev=grammar_famd$eig[, 2],
                           phonology=phonology_pc, phonology_ev=phonology_famd$eig[, 2],
                           music=music_pco, music_ev=music_pcoa$values$Rel_corr_eig,
                           geo=geo_mem)

# Find all factor combinations
factor_names <- c("genetics", "music", "grammar","phonology")
factor_combinations <- get_all_factor_combinations(factor_names)
```

##### S5.1.1 Setup 0: RDA

In this setup we perform an RDA without controlling for potential confounding factors. We use each of the factors as either the response or the explanatory variable in turns. This yields twelve RDA pairs for which we report the coefficient of determination (adjusted $R^2$) and the significance of the correlation determined by an ANOVA like permutation test. We perform the RDA with $k = 15\%$, i.e. we retain all $k$ PCs/PCos which account for at least 15% of the explained variance.

```
# Iterate over all factor combination
rda_setup_0 <- lapply(factor_combinations, function (x) {
  rda_wrapper_setup_0(factors[x[1]][[1]],
                      factors[x[2]][[1]], n_perm=10000)})
```

We simplify the RDA results and adjust the significance for multiple testing using False Discovery Rate. Finally, we plot the explained variance and significance for each association.

```
rda_correlation_matrix <- rda_to_correlation_matrix(rda_setup_0, adjust_significance=T)
plot_rda_as_matrix(rda_correlation_matrix)
```

We observe two significant associations, between grammar and genetics on the one hand, and genetics and grammar on the other. We will now explore if these associations hold in a partial RDA, i.e. in an RDA which accounts for the influence of spatial autocorrelation and genealogy.

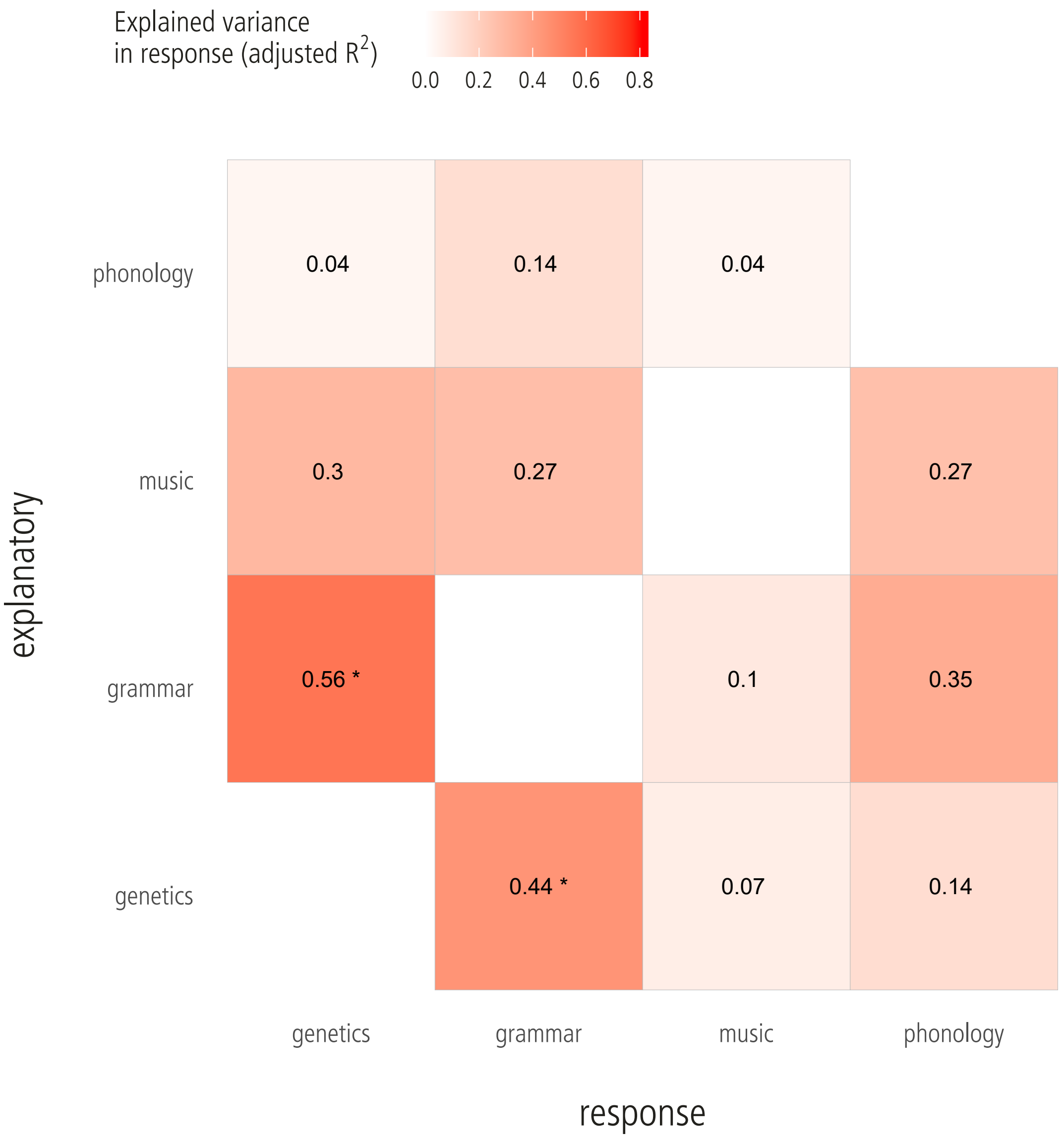

Figure S15: RDA: Variance in the response explained by each explanatory variable; * indicates a significant association ($p \leq 0.05$).

##### S5.1.2 Setup 1: Partial RDA controlling for spatial autocorrelation

Next, we perform a partial RDA where we explore the influence of space on the observed associations. Partial RDA removes the effects of one or more explanatory variables on the response prior to an ordinary RDA (Borcard, Legendre, and Drapeau 1992). In our case, the influence of space is removed. Since most languages in the sample occupy relatively large territories (Fig. S1), choosing a central point location might yield a misleading picture of the possible spatial interactions. Instead, we randomly sample 1,000 spatial locations from the language polygons and perform a partial RDA for each. With each sample we remove the influence for one possible scenario of spatial neighborhood or spatial autocorrelation.

```
rda_setup_1 <- lapply(factor_combinations, function (x) {
  rda_wrapper_setup_1(factors[x[1]][[1]],
                      factors[x[2]][[1]],
                      factors$geo, n_perm=100)})
# Rename the list entries
names(rda_setup_1) <- sapply(rda_setup_1, function (q){
  return (paste(q[[1]]$explanatory, q[[1]]$response, sep="_"))})
```

We compare the distribution of the adjusted $R^2$ across the 1,000 spatial locations against a distribution of adjusted $R^2$ based on random permutations ($N = 100$), i.e. samples for which the rows of the explanatory variable were randomly permuted for each run of the partial RDA. The two density plots belwo show the difference between observed and permuted adjusted $R^2$ for the association between genetics and grammar and between grammar and genetics.

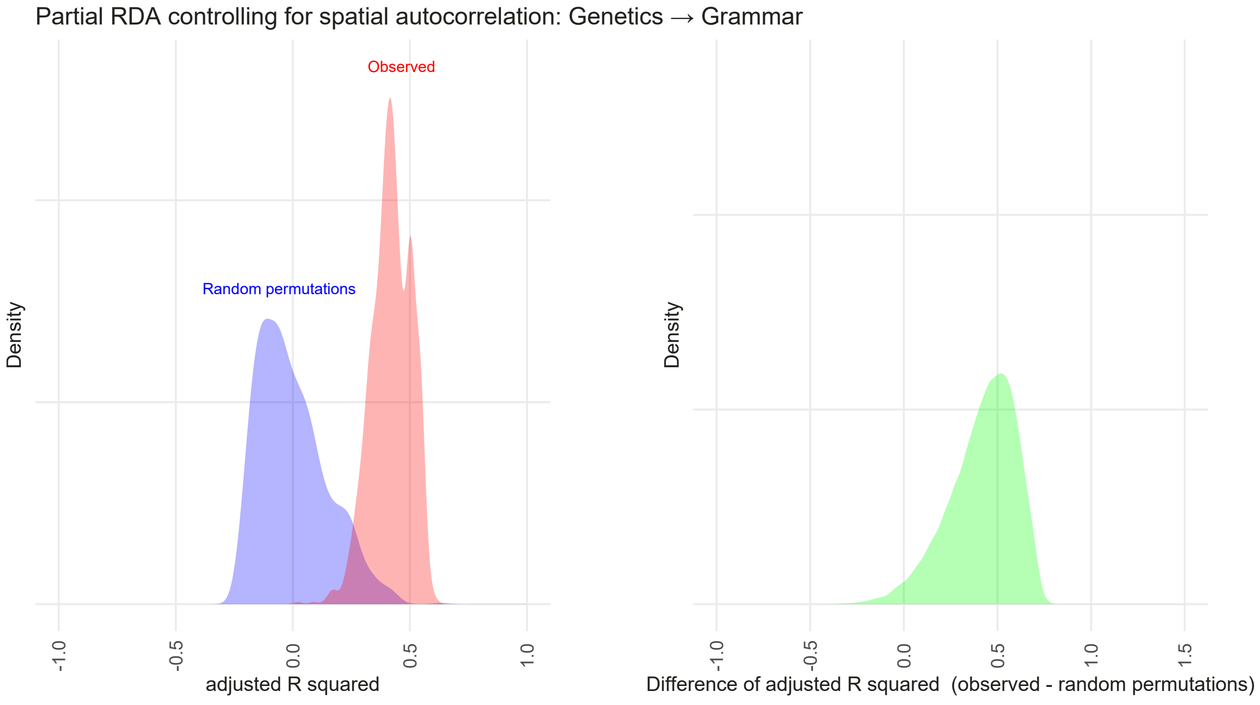

Figure S16: Partial RDA controlling for spatial autocorrelation: The association between Genetics (explanatory variable) and Grammar (response). In the figure on the left, the red curve shows the observed (adjusted) $R^2$, the blue curve the $R^2$ for random permutations. In the figure on the right, the green curve shows the difference of $R^2$ (observed - random permutations).

The difference between the observed and the permuted adjusted $R^2$ needs to be statistically evaluated to determine whether the distribution of observed adjusted $R^2$ is likely to produce values equal or larger than

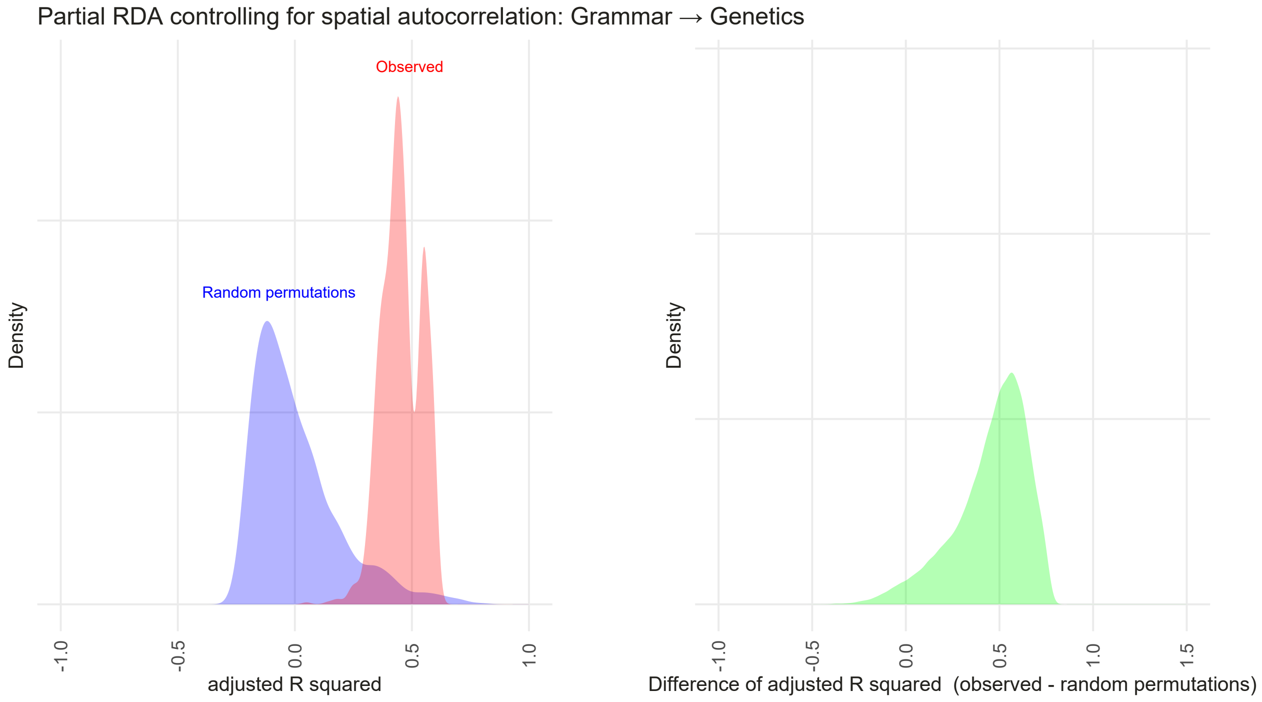

Figure S17: Partial RDA controlling for spatial autocorrelation: The association between Grammar (explanatory variable) and Genetics (response). In the figure on the left, the red curve shows the observed (adjusted) $R^2$, the blue curve the $R^2$ for random permutations. In the figure on the right, the green curve shows the difference of $R^2$ (observed - random permutations).

the permuted one. To do so, we asses the $z$-score of each sample, defined as $z = \Delta R^2/\text{sd}(R^2_{permuted})$, where $\Delta R^2 = R^2_{observed} - E[R^2_{permuted}]$ and $R^2$ refers to the adjusted $R^2$. The expected values and standard deviations are estimated on the sample of all $1,000$ randomly sampled spatial location.

In order to assess how robust the results are across the 1,000 samples we report:

- all samples with positive z-scores ($P(z > 0)$ SD), i.e. samples where the observed adjusted $R^2$ is larger than the permuted
- all samples with strong positive z-scores ($P(z > 1)$ SD) i.e. samples, where the observed adjusted $R^2$ is one standard deviation larger than the permuted
- the Kullback-Leibler divergence (KLD) between the distribution of observed adjusted $R^2$ and permuted adjusted $R^2$.

The KLD allows to assess the overall divergence of the two distributions, $P(z > 0)$ SD and $P(z > 1)$ SD report the the proportion of (strongly) positive differences.

```
rda_setup_1_z <- partial_rda_to_z_val(rda_setup_1, r2_type="r2_adj_partial")
proportion_rda_setup_1 <- add_proportion_and_kld(rda_setup_1_z, diff_th = 1.)
```

We visualize the difference between observed and permuted adjusted $R^2$ in a ridge plot.

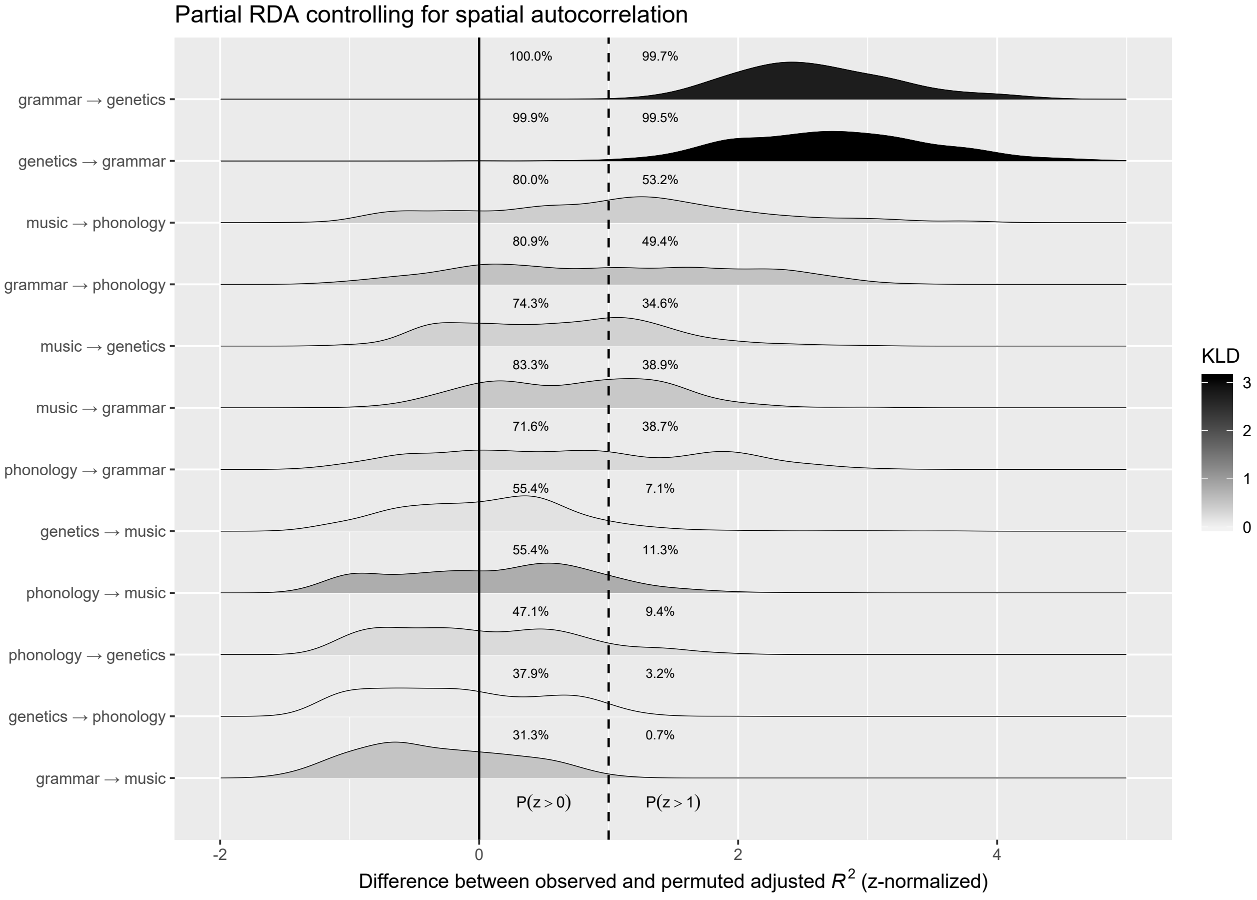

Figure S18: Partial RDA controlling for spatial autocorrelation: Densities of the difference between observedand permuted adjusted $R^2$ (z-normalized). All input components contribute at least 15% to the explained variance. Numbers right to the solid and dashed line show the proportion of samples with a positive difference – $P(z > 0)$ – and a strong positive difference – $P(z > 1$ SD), respectively. Grey shading reflects the Kullback-Leibler divergence (KLD) between the observed and permuted adjusted $R^2$.

##### S5.1.3 Setup 2: Partial RDA controlling for spatial autocorrelation and genealogy

In addition to space, we now also control for the possible influence of phylogenetic inheritance in the three cases where we have data from the same language families (Even and Evenki from the Tungusic, Selkup and Nganasan from the Uralic, and Koryak and Chukchi from the Chukotko-Kamchatkan family).

```
rda_setup_2 <- lapply(factor_combinations, function (x) {
  rda_wrapper_setup_2 (factors[x[1]][[1]],
                       factors[x[2]][[1]],
                       factors$geo, n_perm=100)})

# Rename the list entries
names(rda_setup_2 ) <- sapply(rda_setup_2 , function (q){
  return (paste(q[[1]]$explanatory, q[[1]]$response, sep="_"))})
```

Again, we compare the distribution of the adjusted $R^2$ across the 1,000 spatial locations against the adjusted $R^2$ based on random permutations ($N = 100$ per sample). The two density plots below show the difference between observed and permuted adjusted $R^2$ for the association between genetics and grammar and between grammar and genetics.

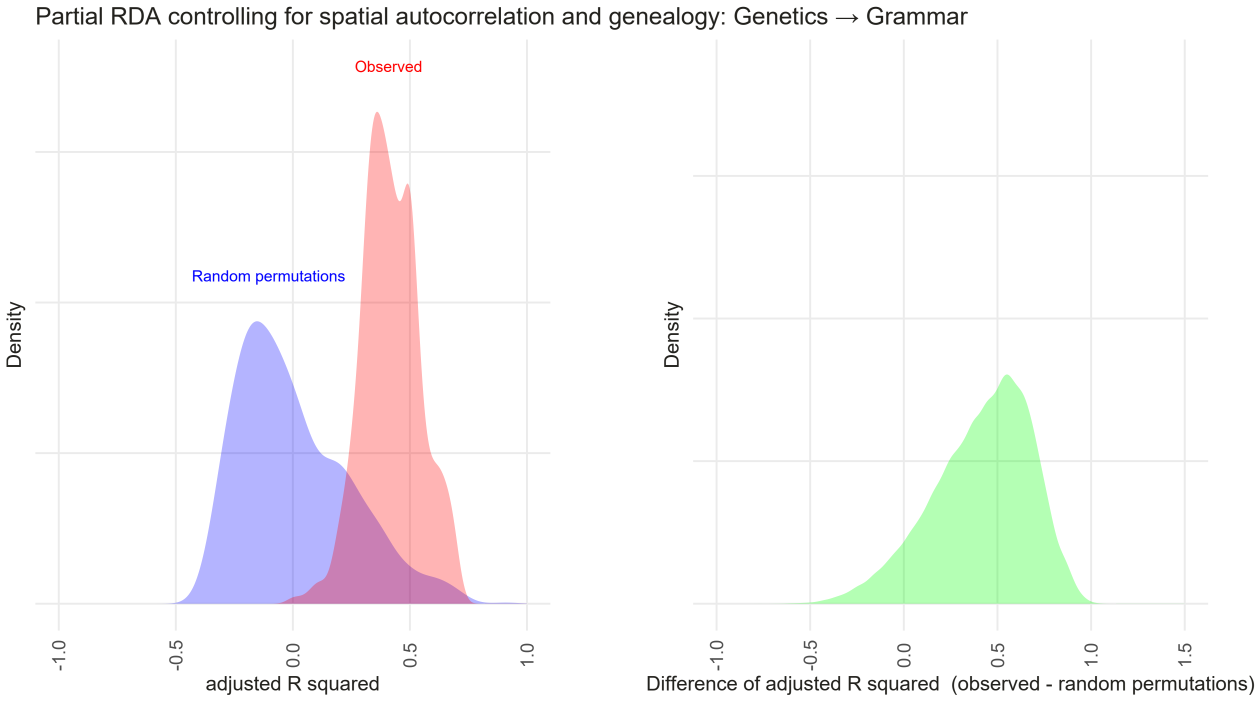

Figure S19: Partial RDA controlling for spatial autocorrelation and genealogy: The association between Genetics (explanatory variable) and Grammar (response). In the figure on the left, the red curve shows the observed (adjusted) $R^2$, the blue curve the $R^2$ for random permutations. In the figure on the right, the green curve shows the difference of $R^2$ (observed - random permutations).

We statistically evaluate the difference between the observed and the permuted adjusted $R^2$ and visualize it in a ridge plot.

```
rda_setup_2_z <- partial_rda_to_z_val(rda_setup_2, r2_type="r2_adj_partial")
proportion_rda_setup_2 <- add_proportion_and_kld(rda_setup_2_z, diff_th = 1.)
```

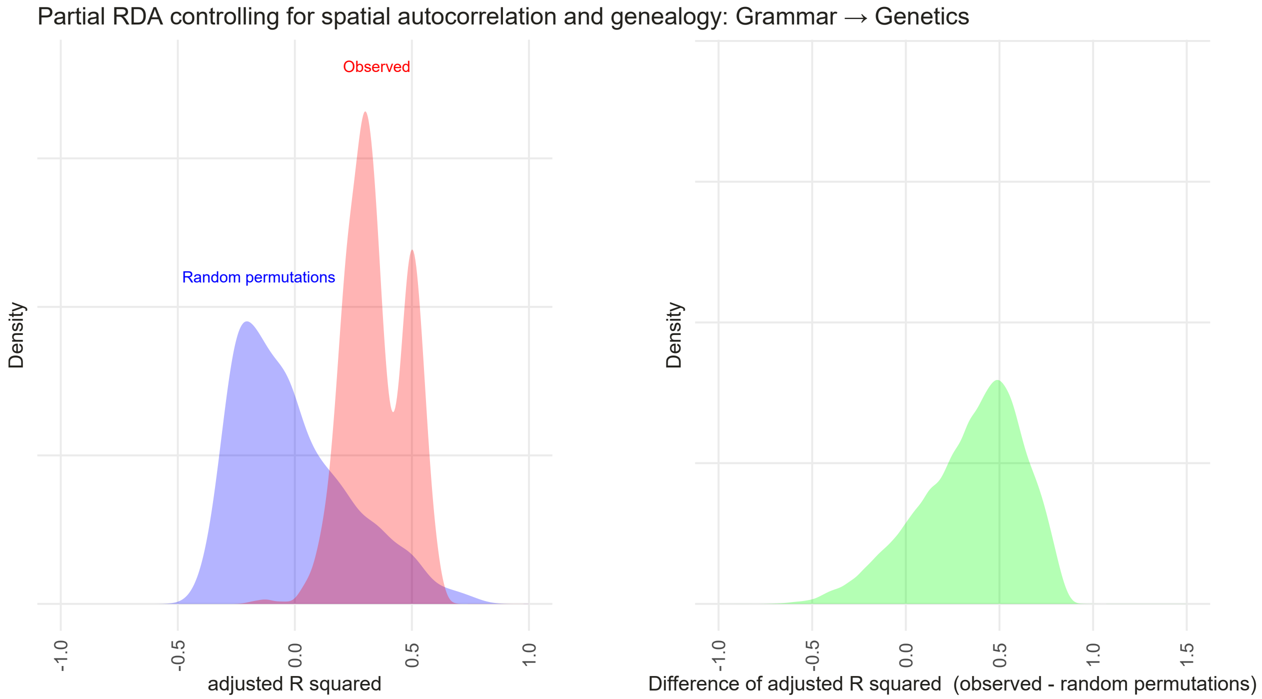

Figure S20: Partial RDA controlling for spatial autocorrelation and genealogy: The association between Grammar (explanatory variable) and Genetics (response). In the figure on the left, the red curve shows the observed (adjusted) $R^2$, the blue curve the $R^2$ for random permutations. In the figure on the right, the green curve shows the difference of $R^2$ (observed - random permutations).

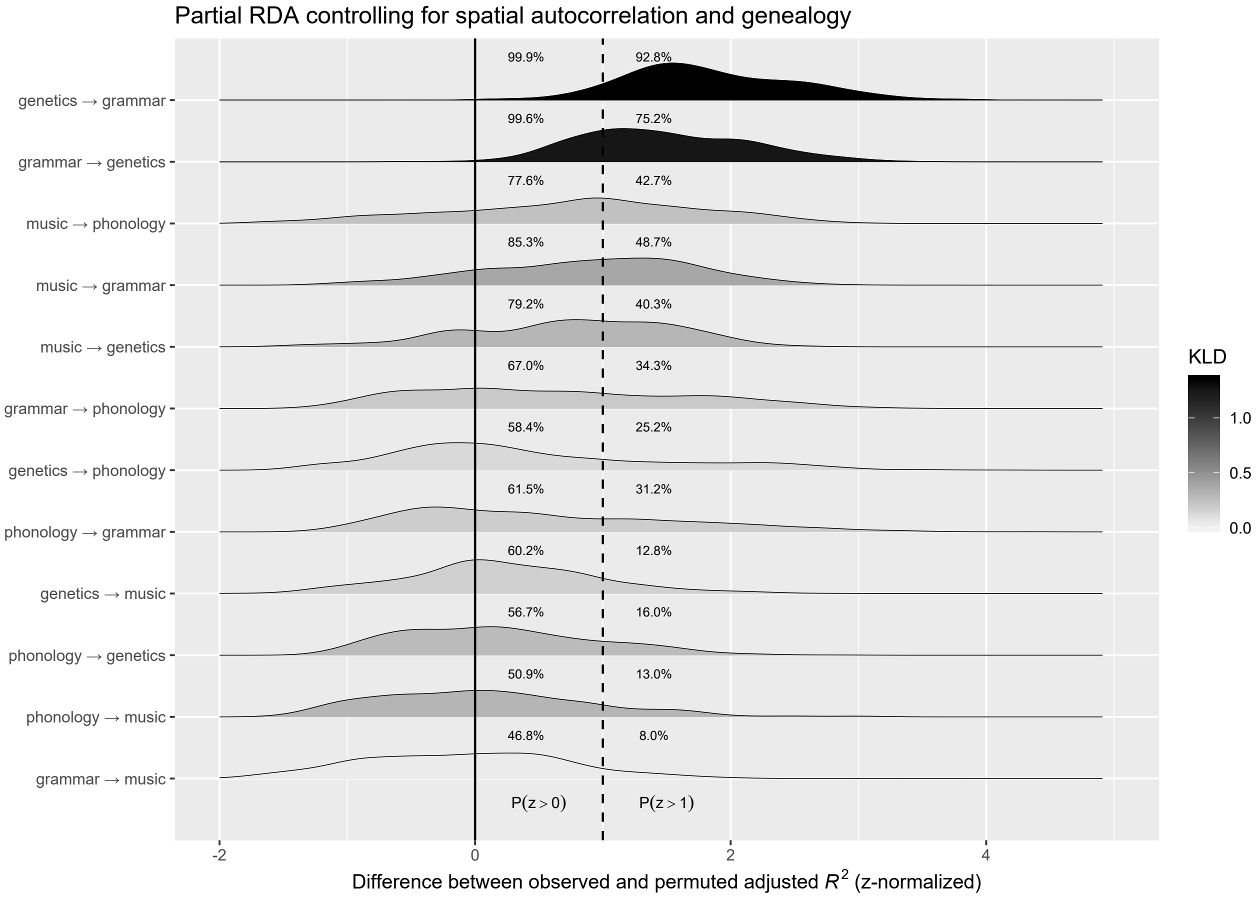

Figure S21: Partial RDA controlling for spatial autocorrelation and genealogy: Densities of the difference between observed and permuted adjusted $R^2$ (z-normalized) in the partial RDA. All input components contribute at least 15% to the explained variance. Numbers right to the solid and dashed line show the proportion of samples with a positive difference – $P(z > 0)$ – and a strong positive difference – $P(z > 1$ SD), respectively. Grey shading reflects the Kullback-Leibler divergence (KLD) between the observed and permuted adjusted $R^2$.

#### S5.2 Sensitivity Analysis

##### S5.2.1 Sensitivity 1: Influence of principal components/coordinates

First we explore to what extent the association between grammar and genetics and genetics and grammar are sensitive to the number of principal components/coordinates used as input. We vary the parameter $k$, which controls the number of PCs/Pcos, such that $k = 0.1$, $k = 0.15$, $k = 0.18$. As before, we run a partial RDA and account for both spatial autocorrelation and genealogy.

```
ks = c(0.1, 0.15, 0.18)

factor_names <- c("genetics", "grammar")
factor_combinations <- get_all_factor_combinations(factor_names)

rda_sens_pc <- list()

for (k in ks) {
  # Retain PCs/PCos which explain at least k%
  factors <- pcos_to_factors(var_th=k,
                              genetics=genetics_pco,
                              genetics_ev=genetics_pcoa$values$Rel_corr_eig,
                              grammar=grammar_pc, grammar_ev=grammar_famd$eig[, 2],
                              phonology=phonology_pc, phonology_ev=phonology_famd$eig[, 2],
                              music=music_pco, music_ev=music_pcoa$values$Rel_corr_eig,
                              geo=geo_mem)
  # Run Partial RDA
  rda_result <- lapply(factor_combinations, function (x) {
    rda_wrapper_setup_2 (factors[x[1]][[1]],
                          factors[x[2]][[1]],
                          factors$geo, n_perm=100)})

  # Rename the list entries
  names(rda_result) <- sapply(rda_result , function (q){
    return (paste(q[[1]]$explanatory, q[[1]]$response, sep="_"))})

  for (n in names(rda_result)){
    rda_sens_pc[[n]][[as.character(k)]] <- rda_result[[n]]
  }}
```

For each $k$, we statistically evaluate the difference between the observed and the permuted adjusted $R^2$ and visualize it in a ridge plot.

```
proportion_rda_sens_pc <- list()

for (n in names(rda_sens_pc)){
    z <- partial_rda_to_z_val(rda_sens_pc[[n]], r2_type="r2_adj_partial")
    proportion_rda_sens_pc[[n]] <- add_proportion_and_kld(z, diff_th = 1.)}
```

##### S5.2.2 Sensitivity 2: Influence of single societies

We explore the influence of sampling on the model. Some societies might have a larger effect on the associations than others. We exclude each society in turn and run a partial RDA. We control for spatial autocorrelation and both spatial autocorrelation and genealogy.

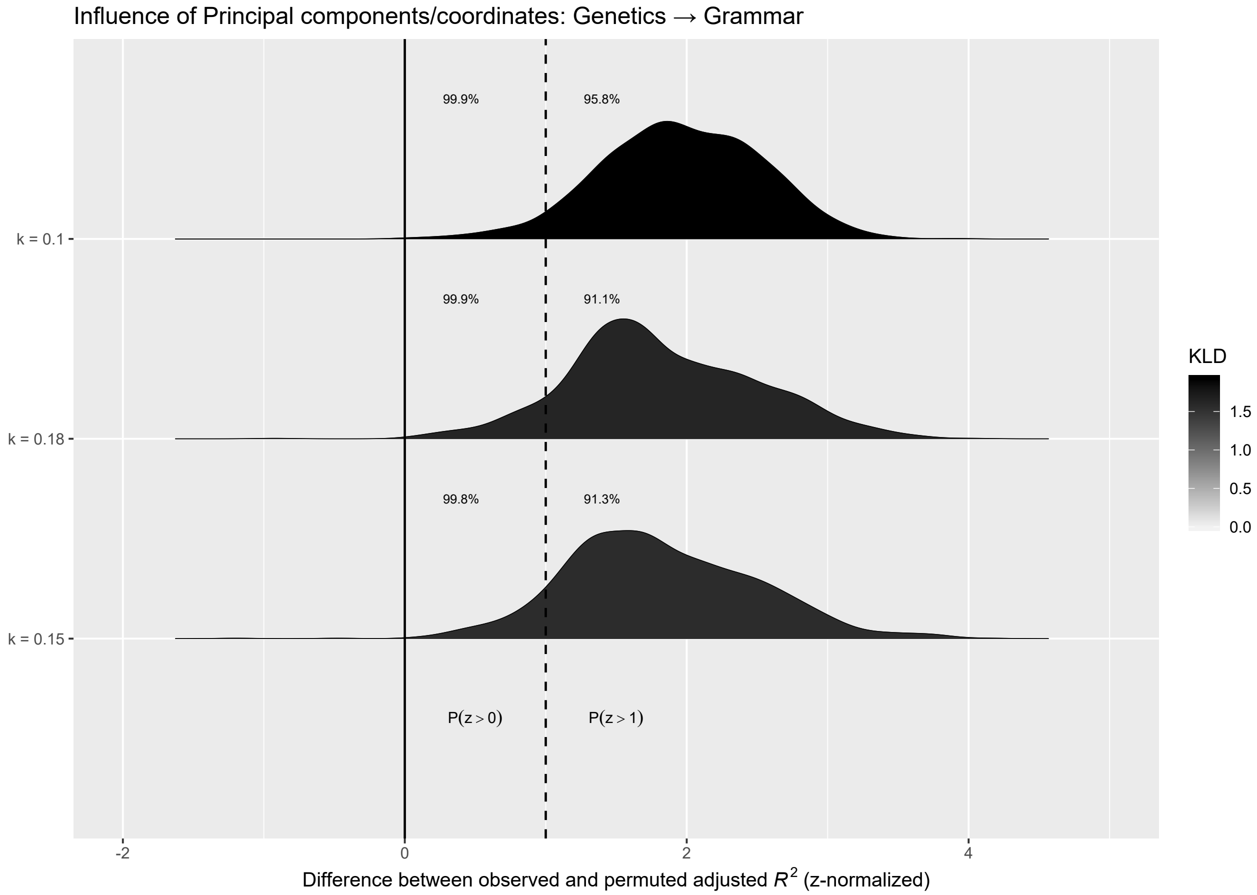

Figure S22: The influence of the number of principal components/coordinates on the association between Genetics (explanatory variable) and Grammar (response). PCs/Pcos must account for at least $k$% of the explained variance. Densities of the difference between observed and permuted adjusted $R^2$ (z-normalized) in the partial RDA. Numbers right to the solid and dashed line show the proportion of samples with a positive difference – $P(z > 0)$ – and a strong positive difference – $P(z > 1$ SD), respectively. Grey shading reflects the Kullback-Leibler divergence (KLD) between the observed and permuted adjusted $R^2$.

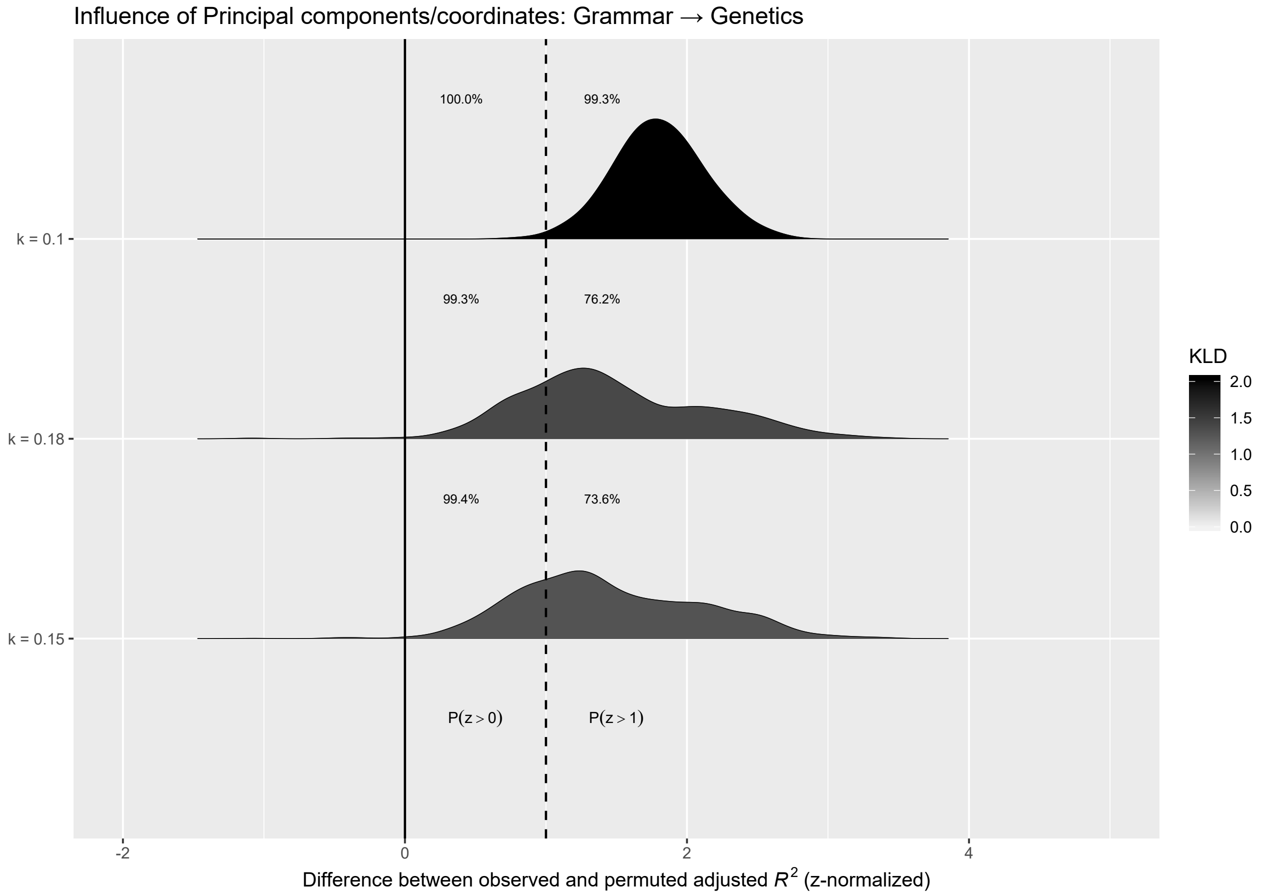

Figure S23: The influence of the number of principal components/coordinates on the association between Grammar (explanatory variable) and Genetics (response). PCs/Pcos must account for at least $k$% of the explained variance. Densities of the difference between observed and permuted adjusted $R^2$ values (z-normlalized) in the partial RDA. Numbers right to the solid and dashed line show the proportion of samples with a positive difference – $P(z > 0)$ – and a strong positive difference – $P(z > 1$ SD), respectively. Grey shading reflects the Kullback-Leibler divergence (KLD) between the observed and permuted adjusted $R^2$.

```r
rda_sens_sites_1 <- list()
rda_sens_sites_2 <- list()

factors <- pcos_to_factors(var_th=0.15,
                           genetics=genetics_pco,
                           genetics_ev=genetics_pcoa$values$Rel_corr_eig,
                           grammar=grammar_pc, grammar_ev=grammar_famd$eig[, 2],
                           phonology=phonology_pc, phonology_ev=phonology_famd$eig[, 2],
                           music=music_pco, music_ev=music_pcoa$values$Rel_corr_eig,
                           geo=geo_mem)
```

```r
for (l in languages) {

  rda_result_1 <- lapply(factor_combinations, function (x) {
    rda_wrapper_sensitivity_1(factors[x[1]][[1]],
                              factors[x[2]][[1]],
                              exclude_site=l,
                              factors$geo, n_perm=100)})

  # Rename the list entries
  names(rda_result_1) <- sapply(rda_result_1 , function (q){
    return (paste(q[[1]]$explanatory, q[[1]]$response, sep="_"))})

  for (n in names(rda_result_1)){
    rda_sens_sites_1[[n]][[as.character(l)]] <- rda_result_1[[n]]}

  rda_sens_sites_1[["n_sites"]][[as.character(l)]] <- rda_result_1[[1]][[1]]$n_sites
}
```

```r
for (l in languages) {

  rda_result_2 <- lapply(factor_combinations, function (x) {
    rda_wrapper_sensitivity_2(factors[x[1]][[1]],
                               factors[x[2]][[1]],
                               exclude_site=l,
                               factors$geo, n_perm=100)})

  # Rename the list entries
  names(rda_result_2) <- sapply(rda_result_2 , function (q){
    return (paste(q[[1]]$explanatory, q[[1]]$response, sep="_"))})

  for (n in names(rda_result_2)){
    rda_sens_sites_2[[n]][[as.character(l)]] <- rda_result_2[[n]]}

  rda_sens_sites_2[["n_sites"]][[as.character(l)]] <- rda_result_2[[1]][[1]]$n_sites
}
```

For each society removed we statistically evaluate the difference between the observed and permuted adjusted $R^2$ and visualize it in a ridge plot.

```
proportion_rda_sens_sites_1<- list()
proportion_rda_sens_sites_2 <- list()

for (n in names(rda_sens_sites_1)[names(rda_sens_sites_1) != "n_sites"]){
    z <- partial_rda_to_z_val(rda_sens_sites_1[[n]], r2_type="r2_adj_partial")
    proportion_rda_sens_sites_1[[n]] <- add_proportion_and_kld(z, diff_th = 1.)}

for (n in names(rda_sens_sites_2)[names(rda_sens_sites_2) != "n_sites"]){
    z <- partial_rda_to_z_val(rda_sens_sites_2[[n]], r2_type="r2_adj_partial")
    proportion_rda_sens_sites_2[[n]] <- add_proportion_and_kld(z, diff_th=1.)}
```

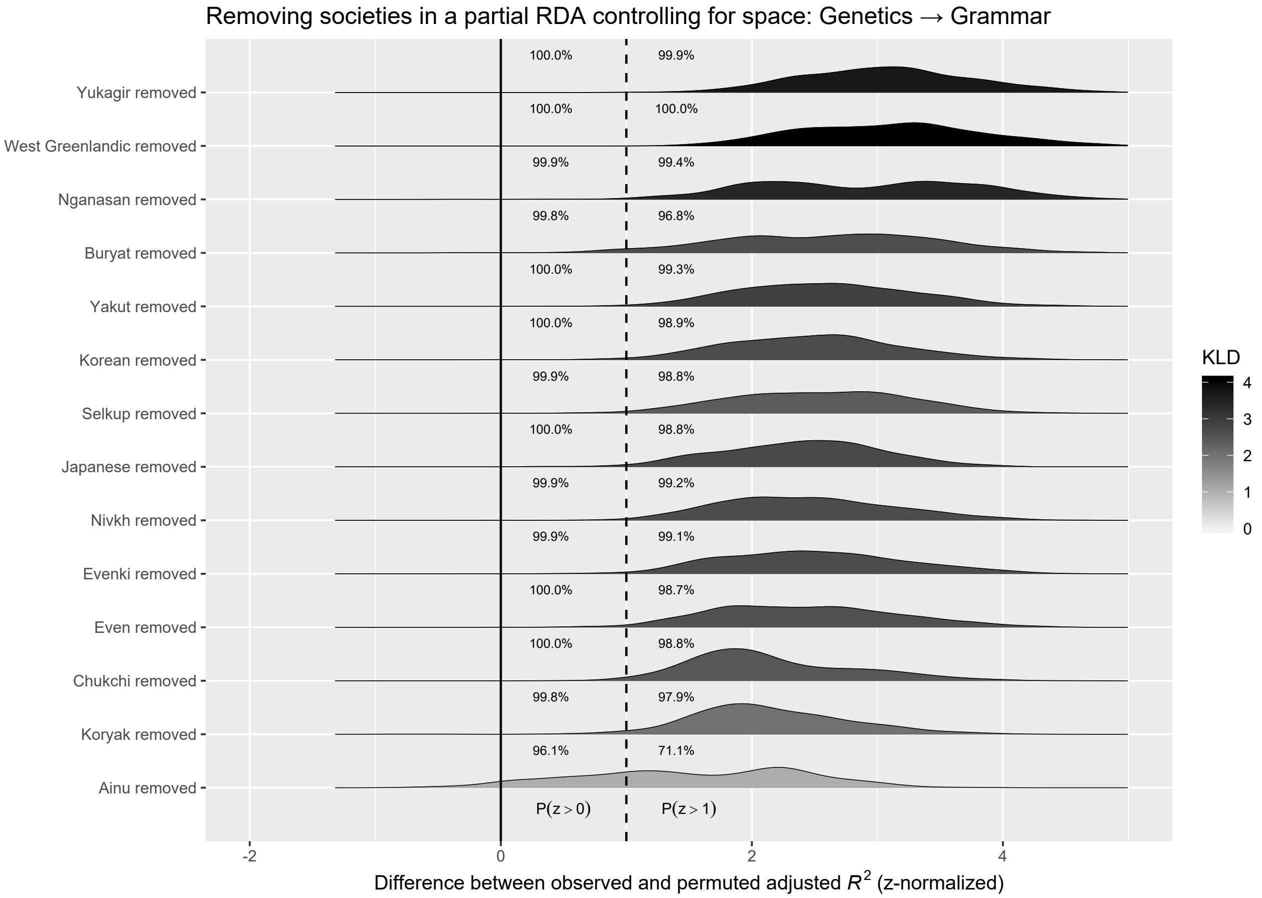

Figure S24: The influence of single societies on the association between Genetics (explanatory variable) and Grammar (response). Densities of the difference between observed and permuted adjusted $R^2$ (z-normalized) in the partial RDA controlling for space but not for genealogy. Numbers right to the solid and dashed line show the proportion of samples with a positive difference – $P(z > 0)$ – and a strong positive difference – $P(z > 1$ SD), respectively. Grey shading reflects the Kullback-Leibler divergence (KLD) between the observed and permuted adjusted $R^2$.

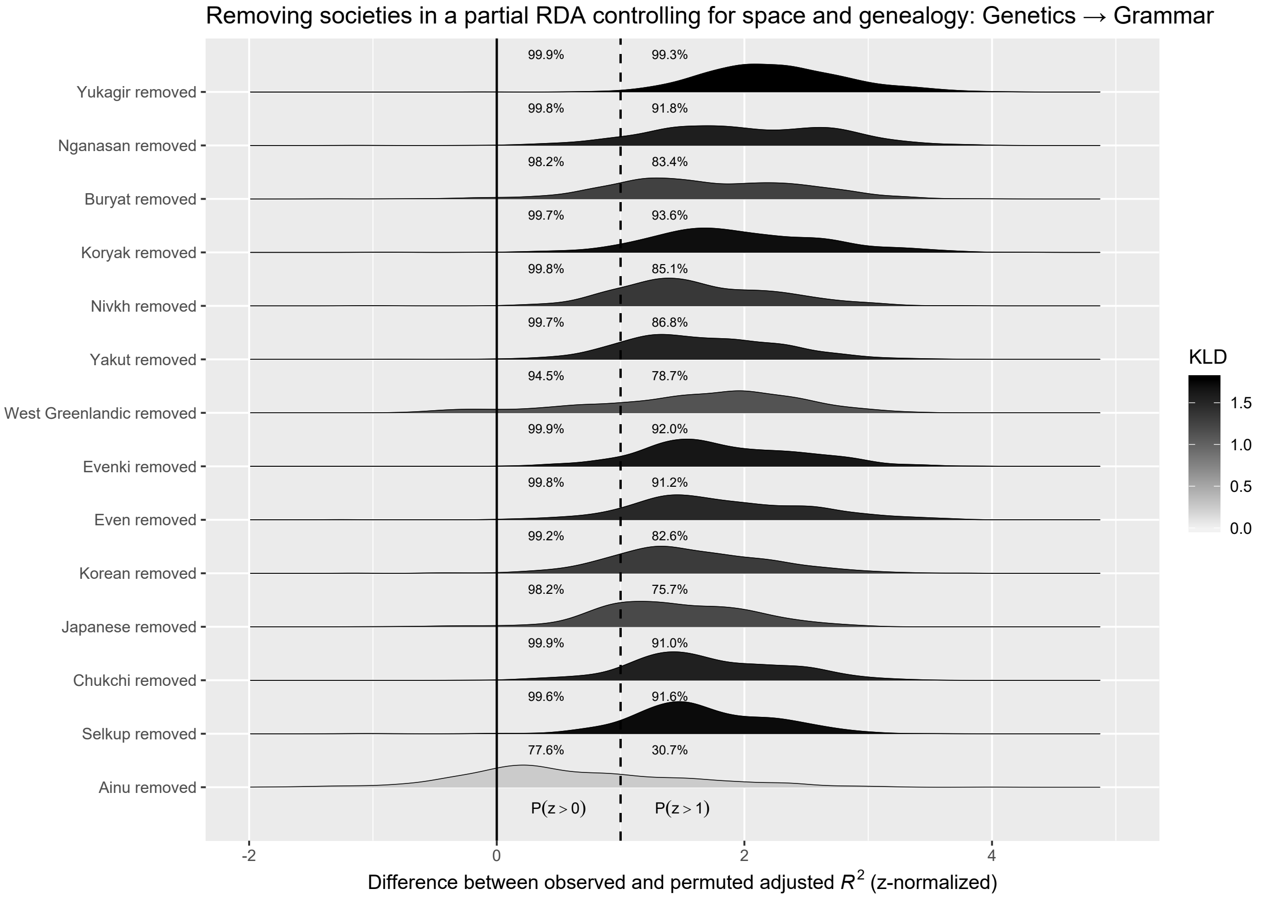

Figure S25: The influence of potential outlier societies on the association between Genetics (explanatory variable) and Grammar (response). Densities of the difference between observed and permuted adjusted $R^2$ (z-normalized) in the partial RDA controlling for space and for genealogy. Numbers right to the solid and dashed line show the proportion of samples with a positive difference – $P(z > 0)$ – and a strong positive difference – $P(z > 1\text{ SD})$, respectively. Grey shading reflects the Kullback-Leibler divergence (KLD) between the observed and permuted adjusted $R^2$.

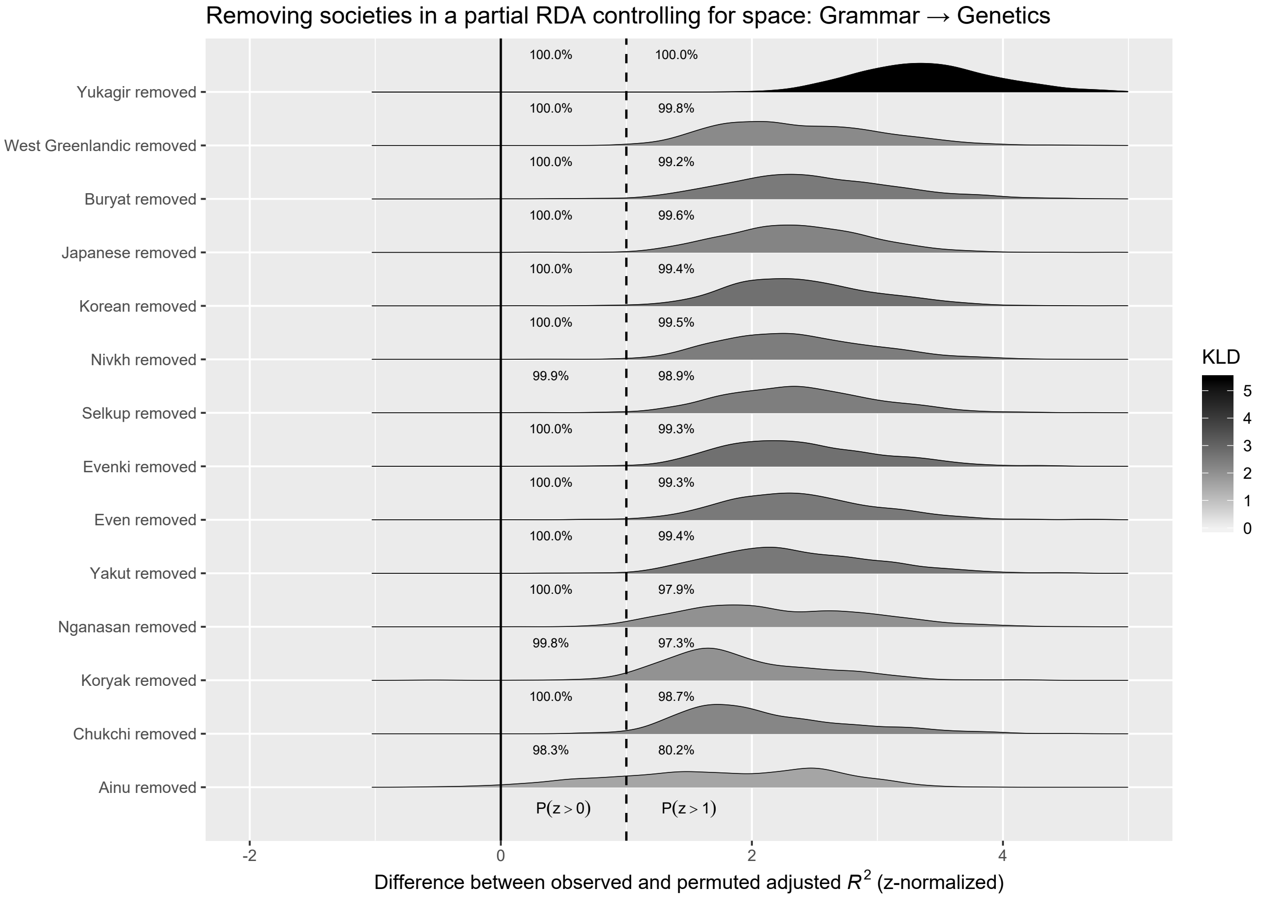

Figure S26: The influence of potential outlier societies on the association between Grammar (explanatory variable) and Genetics (response). Densities of the difference between observed and permuted adjusted $R^2$ (z-normalized) in the partial RDA controlling for space but not for genealogy. Numbers right to the solid and dashed line show the proportion of samples with a positive difference – $P(z > 0)$ – and a strong positive difference – $P(z > 1\text{ SD})$, respectively. Grey shading reflects the Kullback-Leibler divergence (KLD) between the observed and permuted adjusted $R^2$.

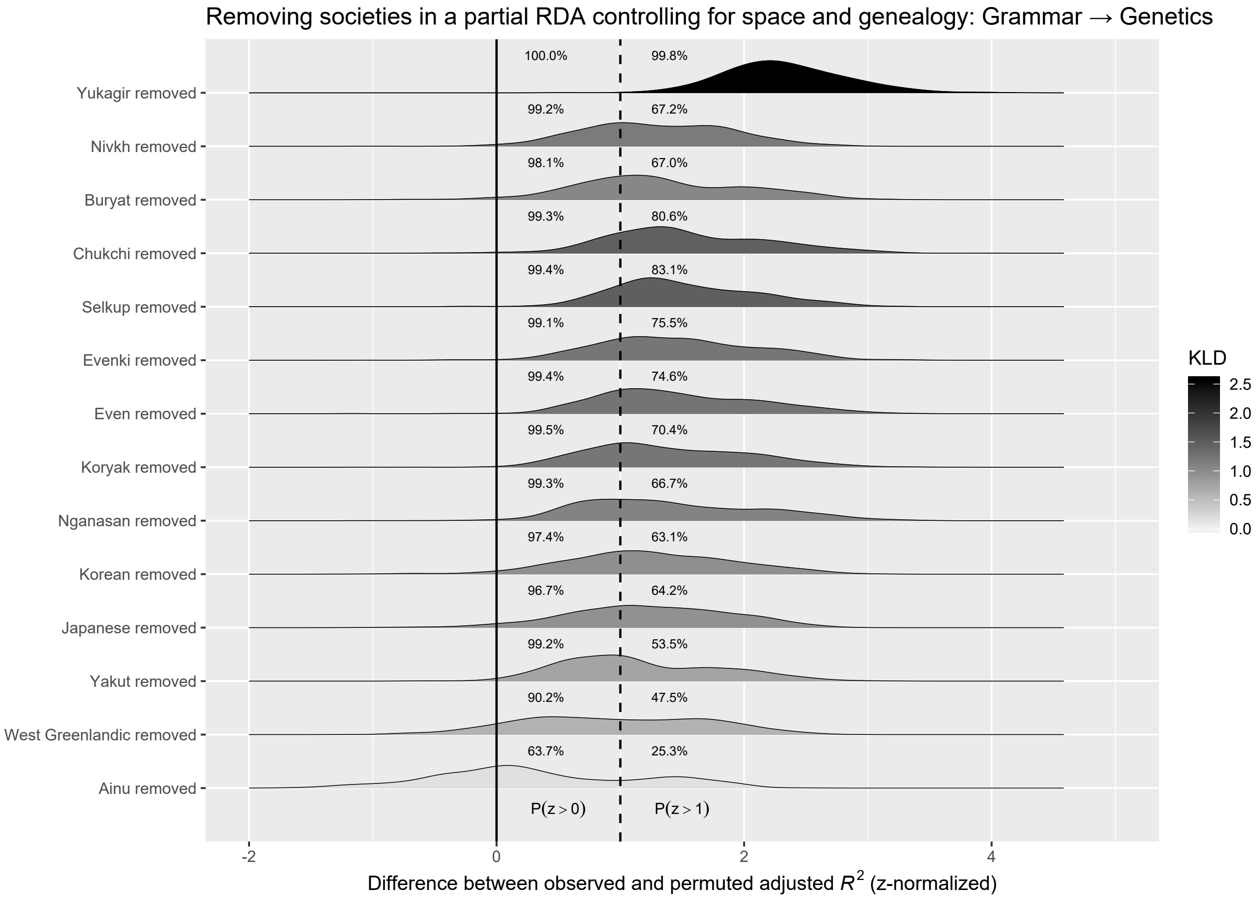

Figure S27: The influence of potential outlier societies on the association between Grammar (explanatory variable) and Genetics (response). Densities of the difference between observed and permuted adjusted $R^2$ (z-normalized) in the partial RDA. Numbers right to the solid and dashed line show the proportion of samples with a positive difference – $P(z > 0)$ – and a strong positive difference – $P(z > 1 \text{ SD})$, respectively. Grey shading reflects the Kullback-Leibler divergence (KLD) between the observed and permuted adjusted $R^2$.

##### S5.2.3 Senisitivity 3: Spatial distribution of $R^2$

Spatial neighborhood is unlikely to explain the observed relationship between factors if the adjusted $R^2$ for all spatial locations is well above zero and if it differs from the adjusted $R^2$ under random permutations, both of which we have shown above. However, the correlations we find might still be an artefact of space, in particular if locations with low adjusted $R^2$ (for example the lower tail of observed distributions in Figures S21) cluster together in different parts of the language polygons than those with particularly high adjusted $R^2$ (e.g. the upper tail of observed distributions in Figures S21) . To rule this out, we map all samples for which the adjusted $R^2$ is in the .2 percentile and the 0.8 percenile, respectivily, and compare both spatial patterns. The following figures suggest that for both the association between grammar and genetics and genetics and grammar locations with low and high $R^2$ randomly distribute in the polygons and do not mass up in specific regions.

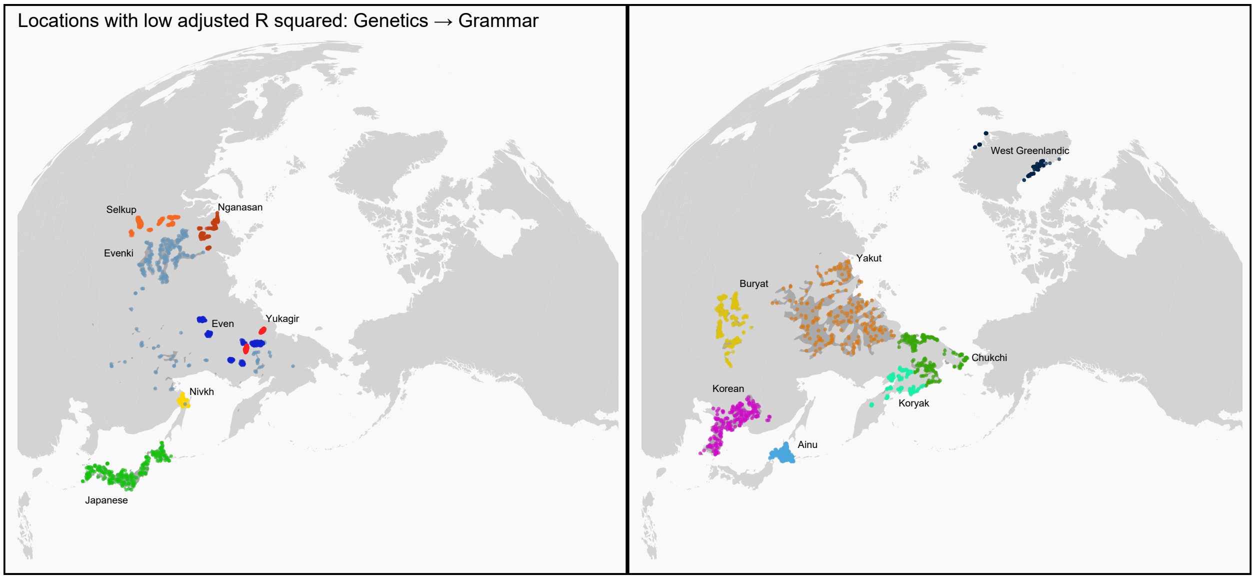

Figure S28: Location samples used for removing the influence of space in the partial RDA between Genetics (explanatory variable) and Grammar (response) with a low adjusted $R^2$

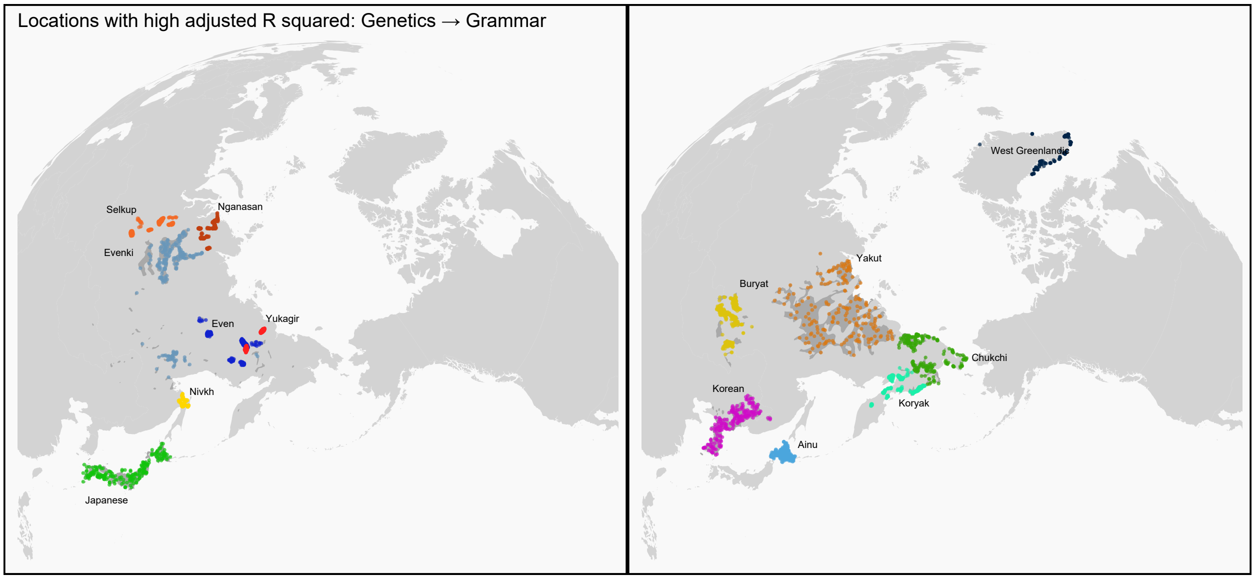

Figure S29: Location samples used for removing the influence of space in the partial RDA between Genetics (explanatory variable) and Grammar (response) with a high adjusted $R^2$

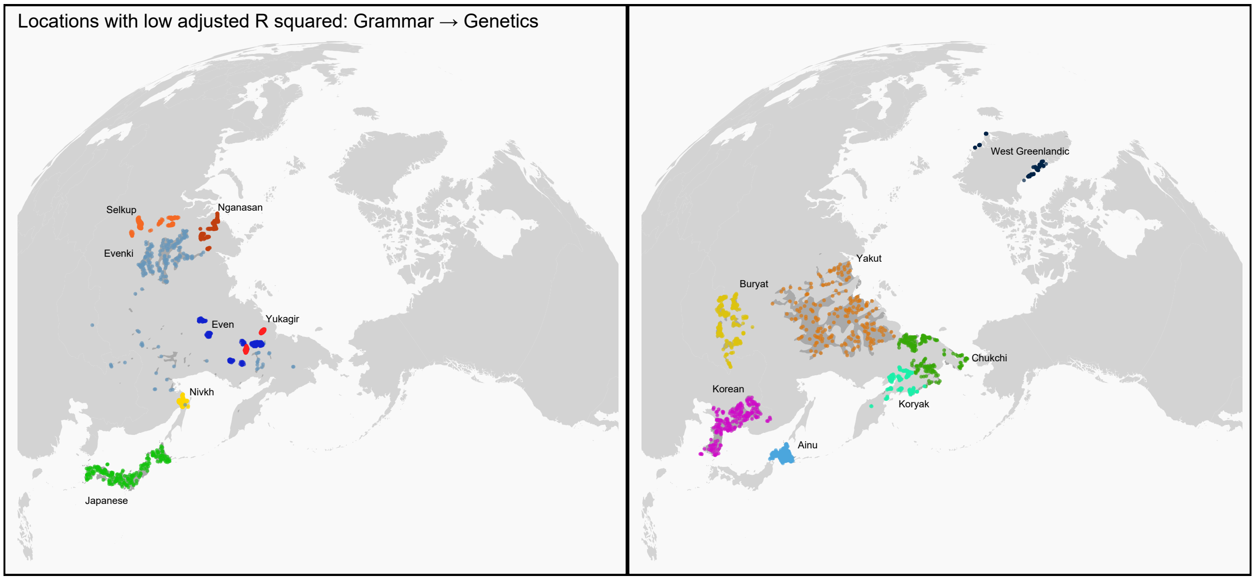

Figure S30: Location samples used for removing the influence of space in the partial RDA between Grammar (explanatory variable) and Genetics (response) with a low adjusted $R^2$

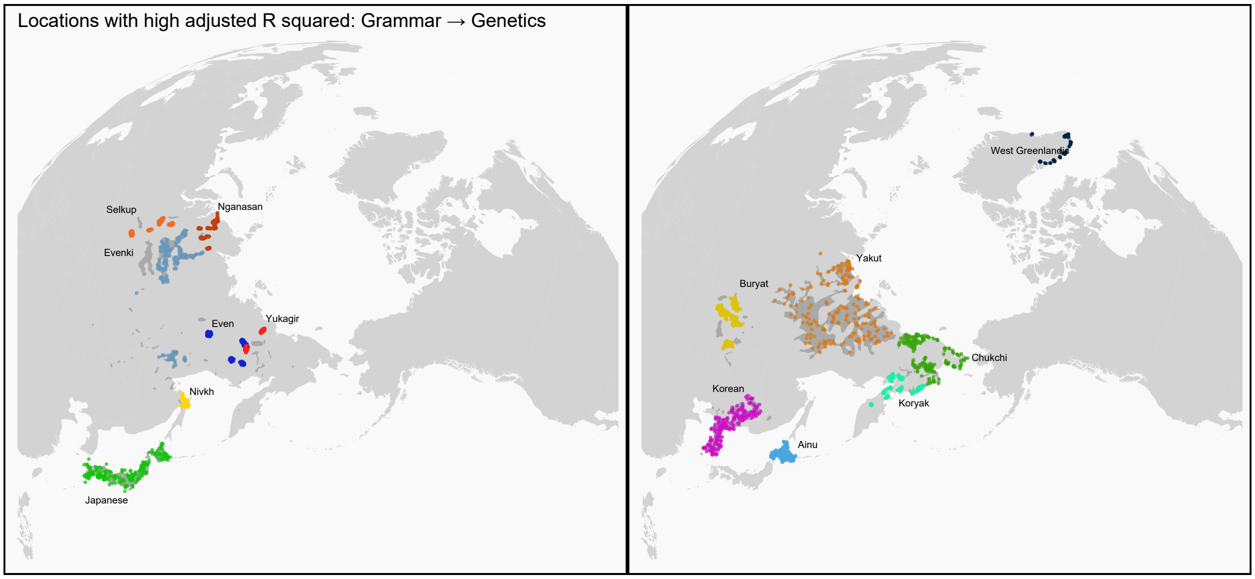

Figure S31: Location samples used for removing the influence of space in the partial RDA between Grammar (explanatory variable) and Genetics (response) with a high adjusted $R^2$

Hammarström, Harald, Robert Forkel, and Martin Haspelmath. 2017. *Glottolog 3.0*. Jena: Jena: Max Planck Institute for the Science of Human History.

Huson, Daniel H, and David Bryant. 2006. "Application of Phylogenetic Networks in Evolutionary Studies." *Molecular Biology and Evolution* 23 (2). Oxford University Press: 254–67.

Jinam, Timothy, Nao Nishida, Momoki Hirai, Shoji Kawamura, Hiroki Oota, Kazuo Umetsu, Ryosuke Kimura, et al. 2012. "The history of human populations in the Japanese Archipelago inferred from genome-wide SNP data with a special reference to the Ainu and the Ryukyuan populations." *Journal of Human Genetics* 57 (12): 787–95. https://doi.org/10.1038/jhg.2012.114.

Josse, Julie, and François Husson. 2016. "missMDA: A Package for Handling Missing Values in Multivariate Data Analysis." *Journal of Statistical Software* 70 (1): 1–31. https://doi.org/10.18637/jss.v070.i01.

Lazaridis, Iosif, Nick Patterson, Alissa Mittnik, Gabriel Renaud, Swapan Mallick, Karola Kirsanow, Peter H. Sudmant, et al. 2014. "Ancient human genomes suggest three ancestral populations for present-day Europeans." *Nature* 513 (7518): 409–13. https://doi.org/10.1038/nature13673.

Legendre, Pierre, and Loic FJ Legendre. 2012. *Numerical Ecology*. Vol. 24. Elsevier.

Lê, Sébastien, Julie Josse, and François Husson. 2008. "FactoMineR: An R Package for Multivariate Analysis." *Journal of Statistical Software* 25: 1–18.

Moran, Steven, Daniel McCloy, and Richard Wright, eds. 2014. *PHOIBLE Online*. Munich: Max Planck Digital Library [http://phoible.org/].

Rasmussen, Morten, Yingrui Li, Stinus Lindgreen, Jakob Skou Pedersen, Anders Albrechtsen, Ida Moltke, Mait Metspalu, et al. 2010. "Ancient human genome sequence of an extinct Palaeo-Eskimo." *Nature* 463 (7282): 757–62. https://doi.org/10.1038/nature08835.

Simons, Gary F., and Charles D. Fennig. 2018. "Ethnologue: Languages of the World, Twenty-First Edition." Dallas, Texas: SIL International. http://www.ethnologue.com.

Van Den Wollenberg, Arnold L. 1977. "Redundancy Analysis an Alternative for Canonical Correlation Analysis." *Psychometrika* 42 (2). Springer: 207–19.

Wichmann, Søren, André Müller, Annkathrin Wett, Viveka Velupillai, Julia Bischoffberger, Cecil H. Brown, Eric W. Holman, et al. 2015. *The ASJP Database (Version 16)*. Repository at http://asjp.clld.org.
